# The new kid on the block: A dominant-negative mutation of phototropin1 enhances carotenoid content in tomato fruits

**DOI:** 10.1101/2020.09.13.295121

**Authors:** Himabindu Vasuki Kilambi, Alekhya Dindu, Kapil Sharma, Narasimha Rao Nizampatnam, Neha Gupta, Nikhil Padmanabhan Thazath, Ajayakumar Jaya Dhanya, Kamal Tyagi, Sulabha Sharma, Sumit Kumar, Rameshwar Sharma, Yellamaraju Sreelakshmi

## Abstract

Phototropins, the UVA-blue light photoreceptors, endow plants to detect the direction of light and optimize photosynthesis by regulating chloroplasts positioning and stomatal gas exchange. Little is known about their functions in other developmental responses. A tomato *Non-phototropic seedling1* (*Nps1*) mutant, bearing an Arg495His substitution in the vicinity of LOV2 domain in phototropin1, dominant-negatively blocks phot1 and phot2 responses. The fruits of *Nps1* mutant were enriched in carotenoids, particularly lycopene, than its parent, Ailsa Craig. Contrarily, CRISPR/CAS9-edited loss of function *phototropin1* mutants displayed subdued carotenoids than the parent. The enrichment of carotenoids in *Nps1* fruits is genetically linked with the mutation and exerted in a dominant-negative fashion. *Nps1* also altered volatile profiles with high levels of lycopene-derived 6-methyl 5-hepten2-one. The transcript levels of several MEP and carotenogenesis pathways genes were upregulated in *Nps1*. *Nps1* fruits showed altered hormonal profiles with subdued ethylene emission and reduced respiration. Proteome profiles showed a causal link between higher carotenogenesis and increased levels of protein protection machinery, which may stabilize proteins contributing to MEP and carotenogenesis pathways. Given the enhancement of carotenoid content by *Nps1* in a dominant-negative fashion, it offers a potential tool for high lycopene-bearing hybrid tomatoes.

**One-sentence summary:** A dominant-negative phototropin1 mutation enhances carotenoid levels, alters metabolite homeostasis, and protein quality control machinery in tomato fruits.

## Introduction

Light is an omnipresent environmental signal which is very effectively utilized by plants to modulate a variety of growth and developmental responses. The capacity to sense light endows plants to monitor seasonal changes, detect the canopy shade, and orient their growth towards or away from light (Nemhauser and Chory, 2002; Kharshiing et al., 2019). In young germinating seedlings, the availability of light is necessary to initiate photomorphogenic development, wherein seedlings become photoautotrophic by transforming plastids to fully functional chloroplasts. (Jenkins et al., 1983; Waters and Langdale, 2009). The formation of chloroplasts is preceded by light-triggered activation of chlorophyll and carotenoid biosynthesis (Kasemir, 1983; von Wettstein et al., 1995; Bartley and Scolnik, 1995; Llorente et al., 2017).

Extensive studies on the light-mediated formation of photosynthetic pigments in seedlings revealed that specific photoreceptors activate key genes involved in chlorophyll and carotenoid biosynthesis (Ilag et al., 1994; Yuan et al., 2017; Huq et al., 2004; Von Lintig et al., 1997). At least five different classes of photoreceptors have been reported in plants viz, phytochromes, cryptochromes, phototropins, zeitlupe/FKF1/LKP1, and UVR8 (Galvão and Fankhauser, 2015; Kharshiing et al., 2019). Among these, phytochrome specifically influences the chlorophyll and carotenoids formation in young seedlings (Huq et al 2004; Von Lintig et al., 1997). The synthesis of chlorophylls and carotenoids in photosynthetic tissues are tightly coupled and regulated in a stoichiometric fashion. Carotenoids acts as photosynthetic accessory pigments and protect chlorophylls from photooxidation (Frank and Cogdell, 1996; Bartley and Scolnik, 1995). Consequently, inhibition of carotenoid biosynthesis leads to photooxidation of chloroplasts with albino or pale green seedlings (Mayfield and Taylor, 1984; Mayfield et al., 1986).

While chlorophyll accumulation is obligatorily linked with carotenoid biosynthesis (Mayfield and Taylor, 1984; Bartley and Scolnik, 1995), the carotenoid biosynthesis can occur independently of chlorophylls. The organs like carrot roots or tomato fruits make a high amount of carotenoids in the chromoplasts (Llorente et al., 2017). Given the diversity of genetic resources available, the tomato has emerged as a model system to study the regulation of carotenogenesis (Barry, 2014; Seymour et al., 2013). The molecular-genetic studies established that carotenogenesis in tomato employs the same biosynthetic pathway as in leaves, albeit with modification. The first enzyme of pathway phytoene synthase is replaced by a chromoplast-specific phytoene synthase 1 (PSY1) and conversion of lycopene to β-carotene is mediated by chromoplast-specific lycopene β-cyclase (CYCB; Hirschberg, 2001). The carotenoids complement of tomato fruits is also distinctly different from leaves, with lycopene and upstream precursors constituting principal carotenoids, rather than xanthophylls, and lutein.

Besides, the initiation of carotenogenesis in tomato fruits is obligatorily coupled with the induction of ripening. Consequently, the non-ripening mutants, such as *Rin*, *Nor*, and *Cnr* fail to accumulate carotenoids in fruits (Seymour et al., 2013). Downstream to ripening regulators, the accumulation of carotenoids is also linked with the climacteric rise of plant hormone ethylene. Therefore, ethylene insensitive mutant such as *Nr* too does not accumulate carotenoids (Barry et al., 2005). Even total suppression of ethylene biosynthesis by transgenic means leads to fruits bereft of carotenoids (Oeller et al., 1991). While ripening regulators and ethylene are essential for the induction of carotenogenesis in fruits, other hormones too modulate the carotenoid biosynthesis. The ABA deficient mutants of tomato show higher carotenoids levels in fruits (Galpaz et al., 2008). Similarly, jasmonic acid also influences carotenoid biosynthesis in fruits (Liu et al., 2012). The level of carotenoid accumulation in fruits seems to be also dependent on the auxin-ethylene balance of fruits, indicating an antagonistic role of IAA (Su et al., 2015), even second messengers like NO affect accumulation of carotenoids in fruits (Bodanapu et al., 2016).

Though multiple factors modulate carotenoid levels in tomato fruits, the light too retains a modulatory role in this process. Tomato fruits grown or incubated in darkness develop normal complement of carotenoids, albeit at lower levels than light-grown plants (Raymundo et al., 1976; Gupta et al., 2014). The mutants defective in light signaling such as *hp1* and *hp2* accumulate a higher amount of carotenoids in fruits, indicating a negative influence of light on carotenogenesis (Kilambi et al., 2013). The influence of the above mutants on the increased carotenoid level seems to be indirect. The higher level of carotenoids in these mutants is ascribed to an increased number of plastids (Azari et al., 2010) and a higher level of carotenoid sequestration proteins in ripened fruits (Kilambi et al., 2013). Among the plant photoreceptors, only cryptochromes, phytochromes, and UVR8 seem to influence carotenogenesis in tomato fruits. The loss of function mutants of cryptochromes and phytochromes and silencing of UVR8 reduces carotenoid levels in fruits (Fantini et al., 2019; Gupta et al., 2014; Bianchetti et al., 2018; Li et al., 2018). Conversely, the overexpression of the above photoreceptors increases the carotenoids level in fruits (Giliberto et al., 2005; Liu et al., 2018; Li et al., 2018; Alves et al., 2020).

Among the plant photoreceptors, phototropin is considered as a receptor that specifically modulates movement responses (Galvão and Fankhauser, 2015). Among the responses mediated by phototropins include phototropic seedling curvature, negative phototropism of roots, stomatal movements, and chloroplast relocation to adapt to high or low light intensity. Though phototropins regulate chloroplast movements, they do not seem to play a role in chloroplast development and function, as the photosynthetic efficiency of *phot2* mutants is not compromised (Gotoh et al., 2018). The studies concerning phototropin function are limited to vegetative tissues and their role in other organs such as fruits is not known.

The uncovering of physiological functions can be best executed by the use of mutants. In an earlier study, we described the isolation and characterization of the *Nps1* mutant bearing a mutation in phototropin1 (Sharma et al., 2014). *Nps1* mutant bearing an Arg 495 His substitution in the vicinity of LOV2 domain dominant negatively blocked both phot1 and phot2 responses in tomato such as chloroplast movement. In this study, we show that *Nps1* also influenced carotenogenesis in fruits in a dominant-negative fashion. The stimulation of carotenogenesis was specific to *Nps1*, as loss of function *phot1* alleles displayed reduced carotenoids levels. Genetic analysis using the introgression of *Nps1* in two distinct cultivars revealed that enhancement of carotenoids is tightly linked to *Nps1* mutation. The influence of *Nps1* was not limited to carotenogenesis, analysis of proteome, metabolome, and volatiles revealed broad spectrum influence of *Nps1* specifically on protein quality control machinery.

## Results

The carotenogenesis in tomato fruits is governed by a complex interaction between endogenous and environmental factors including light acting via phytochromes and cryptochromes (Liu et al., 2015). Conversely, it is not known whether phototropins too contribute to carotenogenesis and ripening. In this study, we show that a dominant-negative mutation in phototropin1- *Nps1*, which suppresses phototropins function, influences fruit development, and considerably enhances carotenoids in tomato.

### *Nps1* mutation impedes phototropins function in fruits

To monitor, whether phototropins are functional in fruits, we examined chloroplast accumulation and avoidance mediated by phototropin1 and phototropin2 respectively. In dark-adapted *Nps1* and its parent Ailsa Craig (AC) fruits, similar to dark-adapted leaves, chloroplasts were localized on the cell periphery (Figure 1A). Consistent with the pale-green color of *Nps1* fruits (Table 1), the chloroplast fluorescence was weaker than AC. The low-fluence blue-light-mediated chloroplast accumulation in the center of cells was seen only in AC but not in *Nps1*. Consequently, the high-fluence blue-light-mediated chloroplast avoidance too was confined to AC. The absence of chloroplast movement indicated that phototropins function is compromised in *Nps1* fruits (Sharma et al., 2014).

**Figure 1.**
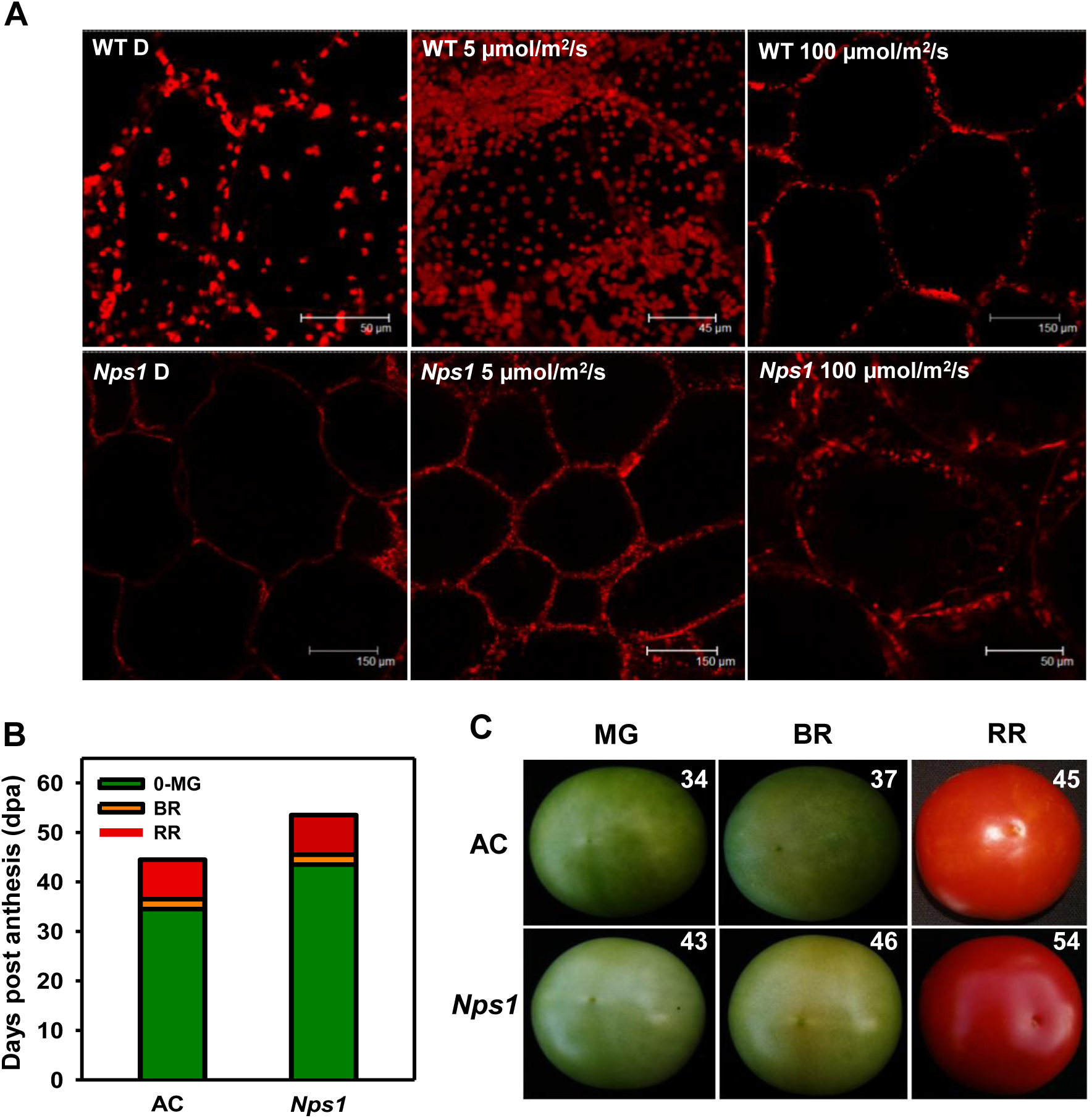
Blue light-induced chloroplast relocations in the fruit pericarp, fruit development and ripening in AC and *Nps1.* **A**, The pericarp slices from dark-adapted MG fruits were exposed to low-fluence blue light (5 µmol/m^2^/s) to induce chloroplast accumulation. Thereafter, slices were exposed to high-fluence (100 µmol/m^2^/s) blue light to induce chloroplast avoidance. Photographs shown are the representative images taken after 30 min of light treatment. **B**, Chronological development of fruits. Note: While *Nps1* is delayed in attaining the MG stage, the duration to attain BR and RR stages was similar to AC (n= 15-20 fruits from 30 plants). **C**, Development of fruit color in AC and *Nps1*. Days post-anthesis are indicated on the top right corner of photos.

**Table 1.**
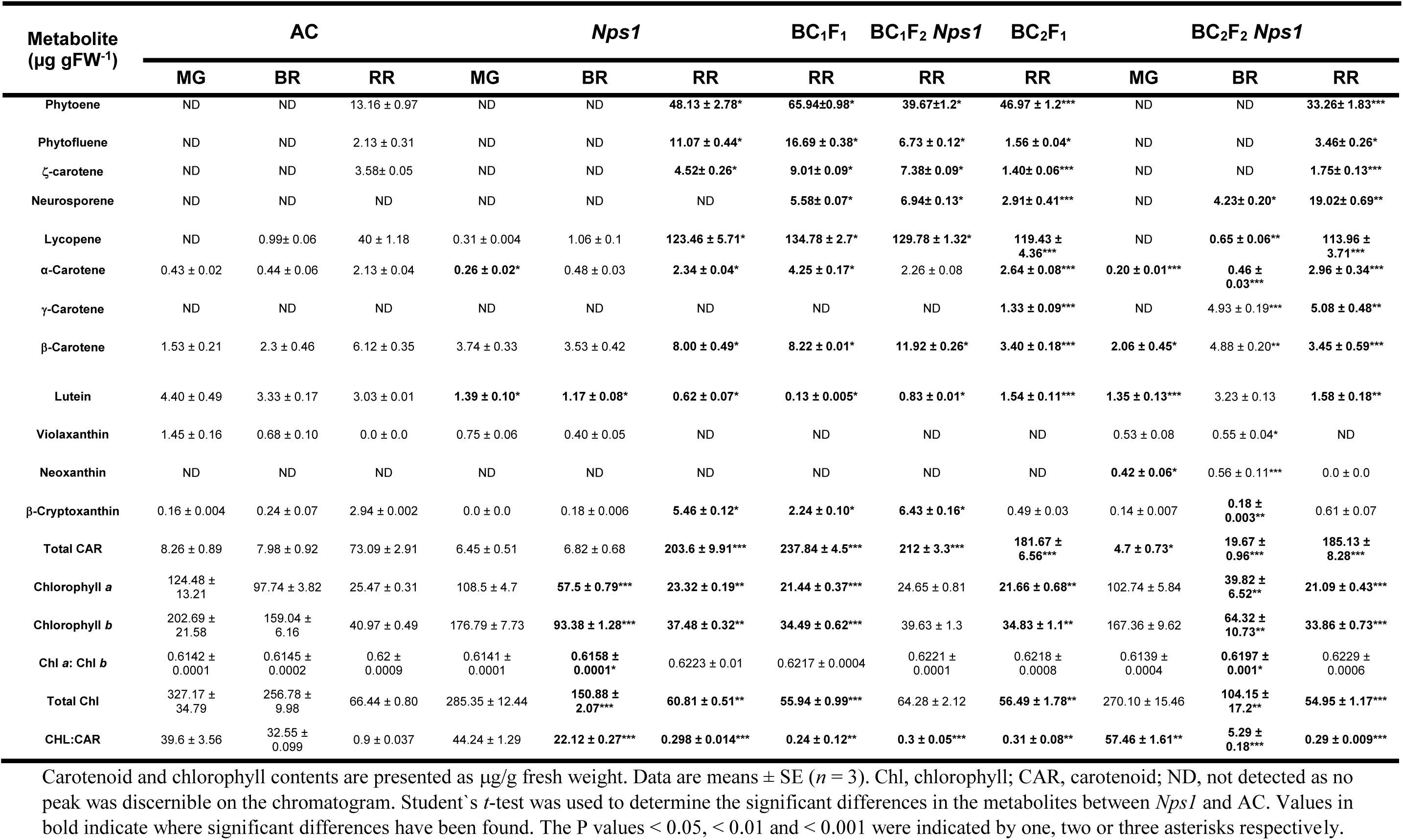
Carotenoid and chlorophyll contents in the fruits of AC and *Nps1* at different ripening stages.

We then examined whether *Nps1* affected fruit development and attainment of ripening stages viz. mature-green (MG), breaker (BR), and red-ripe (RR) compared to AC. Interestingly, the days post-anthesis (dpa) to attain MG was 8 days longer in *Nps1* than AC (34 dpa). However, post-MG, the duration to MG◊BR and BR◊RR in *Nps1* and AC) were nearly similar (Figure 1B).

### *Nps1* fruits accumulate high levels of carotenoids

Though MG fruits of *Nps1* were pale-green, the on-vine ripened RR fruits exhibited intense-red coloration than AC (Figure 1C). The profiling of carotenoids revealed the presence of at least ten carotenoids in AC and *Nps1* fruits (Table 1). Consistent with intense-red coloration, *Nps1* RR fruits had 3.1-fold higher lycopene. Similarly, the levels of precursors to lycopene were high in *Nps1* RR fruits. The increase in lycopene levels in *Nps1* took place during BR to RR transition. In MG and BR fruits, the lycopene levels did not markedly differ in *Nps1* and AC. Although *Nps1* showed higher β-carotene levels, its increase was subdued compared to lycopene. The contribution of carotenoids downstream to lycopene to the carotenoid pool was minor.

### Increased carotenogenesis in fruits is linked to dominant negative function of *Nps1*

Compared to loss-of-function mutants, the dominant-negative mutants are rare. Nonetheless, the dominant-negative mutations are useful to elucidate redundant functions, as mutant protein suppresses the function of its partners. Consequently, the dominant-negative alleles strongly affect the phenotype of heterozygotes, which often resembles a knockout mutation (Meinke, 2013). Considering that *Nps1* RR fruits display a high level of lycopene and its precursors, the above upregulation may be linked to *Nps1* mutation. To ascertain this, we crossed *Nps1* with its parent AC. We also crossed *Nps1* with Arka Vikas (AV), to ascertain whether its dominant-negative effect is retained in distant genetic background.

Akin to reported residual phototropism in F_1_ (Sharma et al., 2014), the F_1_ seedlings showed slight curvature. Remarkably, the F_1_ RR fruits also exhibited high lycopene similar to *Nps1* in consonance with the dominant nature of the *Nps1* mutant (Table 1 and 2). The presence of a mutated copy of the *phot1* gene in F_1_ seedlings was also confirmed by the CELI endonuclease assay (Supplemental Figure 1). The analysis of F_2_ seedling’s phototropism along with the CELI endonuclease assay revealed a typical Mendelian segregation ratio for dominant mutation close to 3:1 (Nonphototropic: Phototropic; Supplemental Table 1). Alike *Nps1*, in BC_1_F_2_ (*Nps1*/*Nps1*) RR fruits, lycopene was considerably higher than either parent. Similarly, BC_2_F_1_ and BC_2_F_2_ (*Nps1*/*Nps1*) progeny too showed elevated levels of lycopene and its precursors in RR fruits (Table 1 and 2). The backcrossed plants retained epinasty-like twisting and downward inclination of expanding apical leaves observed for *Nps1.* The plants were also more robust with increased branching than AC (Figure 2A-B). However, the chlorophyll and carotenoids levels in leaves of *Nps1* and BC_2_F_4_ *Nps1* were akin to AC (Supplemental Table 2). Further studies were carried out using homozygous BC_2_F_2_ (*Nps1/Nps1)* and its F_3_/F_4_ progeny in AC, which altogether for convenience is referred to as *Nps1\**.

**Figure 2.**
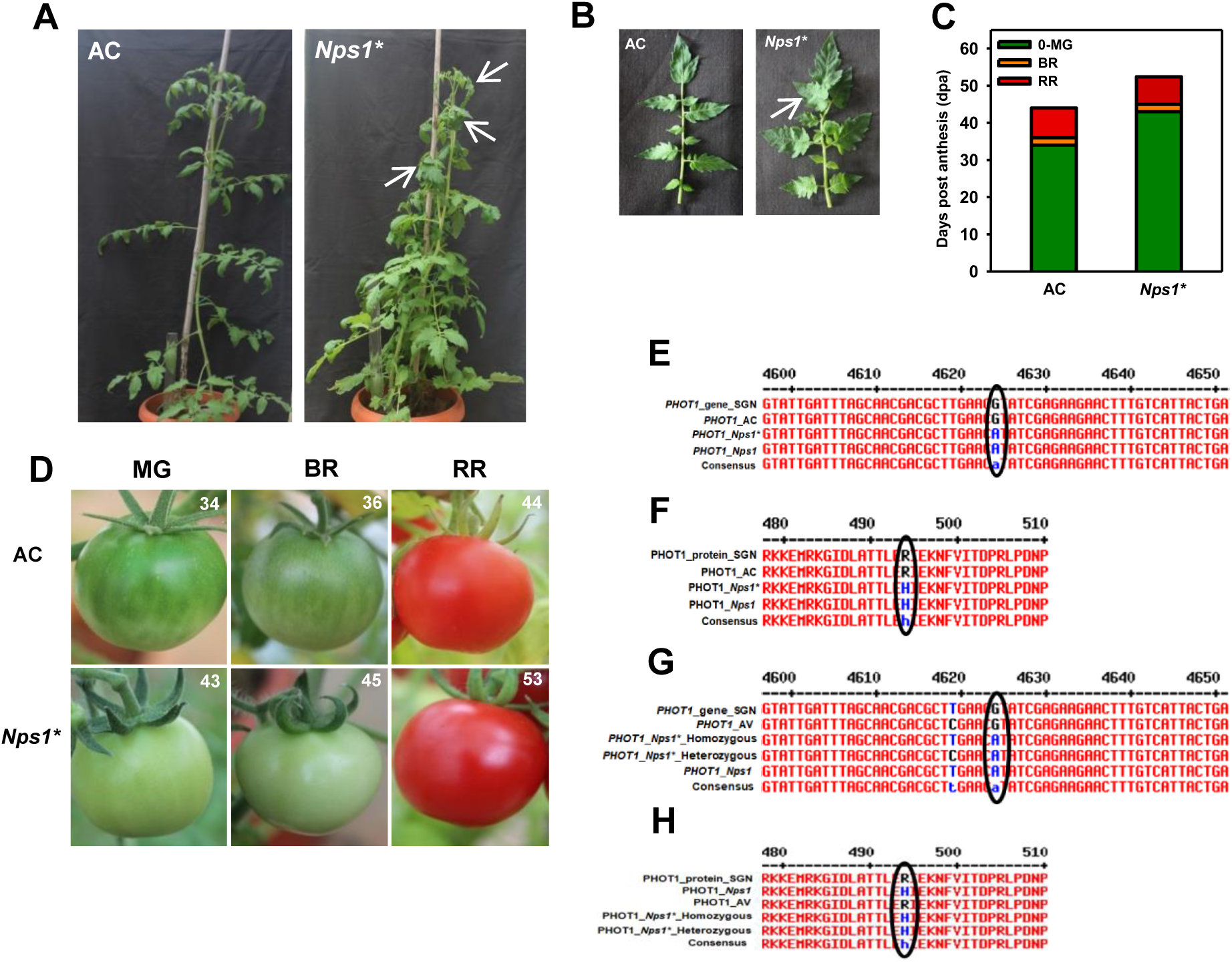
The phenotype of homozygous *Nps1* introgressed plants. **A-B**, Phenotype of AC, and *Nps1\** (BC_2_F_3_) plants (**A**) and leaves (**B**). Note: *Nps1\** shows more biomass. White arrows point to leaf curling and epinasty in *Nps1\** (**A**). Note: *Nps1*\* leaves show twisted rachis (**B**). **C**, Chronological development of fruits (n=15-20 fruits from 30 plants). Note: fruit development and ripening duration in *Nps1*\* is similar to *Nps1*. **D**, Development of fruit color in AC and *Nps1\**. Days post-anthesis are indicated on the top right corner of photos. **E-H**, Confirmation of introgression of *Nps1\** mutation in AC (**E, F**) and AV (**G, H**). **E**, **G**-Alignment of *phot1* gene (Solyc11g072710) sequence. **F, H**-Alignment of PHOT1 amino acid sequences. Note: In recent ITAG3.2 annotation the mutation is located at R494H.

**Table 2.** Carotenoid and chlorophyll contents in the fruits of AV and *Nps1* at different ripening stages.

| Metabolite<br>($\mu\text{g gFW}^{-1}$) | AV | | | *Nps1* | | | BC<sub>1</sub>F<sub>1</sub> | BC<sub>1</sub>F<sub>2</sub> *Nps1* | BC<sub>2</sub>F<sub>1</sub> | BC<sub>2</sub>F<sub>2</sub> *Nps1* |
| --- | --- | --- | --- | --- | --- | --- | --- | --- | --- | --- |
|  | MG | BR | RR | MG | BR | RR | RR | RR | RR | RR |
| Phytoene | ND | ND | 14.46 ± 0.34 | ND | ND | 48.13 ± 2.78* | 55.11 ± 1.84* | 25.59 ± 0.6*** | 39.11 ± 0.06*** | 30.59 ± 0.52*** |
| Phytofluene | ND | ND | 4.7 ± 0.88 | ND | ND | 11.07 ± 0.44* | 14.04 ± 1.44* | 0.21 ± 0.01** | 9.04 ± 0.94* | 8.05 ± 0.79* |
| ζ-carotene | ND | ND | 5.14 ± 0.42 | ND | ND | 4.52 ± 0.26* | 7.24 ± 0.47 | 0.19 ± 0.01*** | 3.24 ± 0.24* | 2.96 ± 0.14** |
| Neurosporene | ND | ND | ND | ND | ND | ND | 3.91 ± 0.48* | 4.3 ± 0.04*** | 2.54 ± 0.19*** | 2.14 ± 0.15*** |
| Lycopene | 0.31 ± 0.007 | 1.05 ± 0.03 | 38.07 ± 4.12 | 0.31 ± 0.004 | 1.06 ± 0.1 | 123.46 ± 5.71* | 153.69 ± 10.26* | 144.68 ± 15.66** | 105.25 ± 9.25** | 91.57 ± 2.3*** |
| α-Carotene | 0.34 ± 0.04 | 0.26 ± 0.02 | 2.16 ± 0.14 | 0.26 ± 0.02* | 0.48 ± 0.03 | 2.34 ± 0.04* | 2.24 ± 0.21 | 3.45 ± 0.23** | 1.85 ± 0.18 | 1.55 ± 0.12* |
| γ-Carotene | ND | ND | ND | ND | ND | ND | ND | ND | 1.33 ± 0.09*** | 5.08 ± 0.48** |
| β-Carotene | 3.93 ± 0.53 | 3.8 ± 0.46 | 3.13 ± 0.36 | 3.74 ± 0.33 | 3.53 ± 0.42 | 8.00 ± 0.49* | 4.27 ± 0.77 | 11.56 ± 1.36** | 6.27 ± 0.35* | 6.78 ± 0.38** |
| Lutein | 4.58 ± 0.41 | 2.33 ± 0.07 | 0.22 ± 0.02 | 1.39 ± 0.10* | 1.17 ± 0.08* | 0.62 ± 0.07* | 0.19 ± 0.003 | 2.14 ± 0.19** | 0.13 ± 0.002** | 0.11 ± 0.001** |
| Violaxanthin | 1.54 ± 0.14 | 0.68 ± 0.10 | ND | 0.75 ± 0.06 | 0.40 ± 0.05 | ND | ND | ND | ND | ND |
| Neoxanthin | 62.77 ± 6.49 | 44.26 ± 0.89 | ND | ND | ND | ND | ND | ND | ND | ND |
| β-Cryptoxanthin | 0.16 ± 0.006 | 0.22 ± 0.04 | 3.36 ± 0.73 | 0.0 ± 0.0 | 0.18 ± 0.006 | 5.46 ± 0.12* | 1.2 ± 0.03* | ND | 0.04 ± 0.003* | 0.026 ± 0.002* |
| Total CAR | 73.63 ± 7.62 | 52.6 ± 1.61 | 71.24 ± 7.01 | 6.45 ± 0.51 | 6.82 ± 0.68 | 203.6 ± 9.91*** | 241.89 ± 15.5*** | 192.12 ± 18.1** | 168.8 ± 11.3** | 148.85 ± 4.88*** |
| Chlorophyll *a* | 109.81 ± 10.43 | 92.07 ± 4.13 | 22.47 ± 0.31 | 108.5 ± 4.7 | 57.5 ± 0.79*** | 23.32 ± 0.19** | 19.78 ± 0.39 | 21.99 ± 0.53 | 18.99 ± 0.24 | 19.42 ± 0.84 |
| Chlorophyll *b* | 190.35 ± 17.69 | 159.70 ± 7.71 | 33.63 ± 1.79 | 176.79 ± 7.73 | 93.38 ± 1.28*** | 37.48 ± 0.32** | 25.16 ± 0.56** | 25.63 ± 2.3* | 28.16 ± 0.93* | 25.19 ± 0.89* |
| Chl *a*: Chl *b* | 0.58 ± 0.003 | 0.616 ± 0.0001 | 0.667 ± 0.03 | 0.6141 ± 0.0001*** | 0.6113 ± 0.0003 | 0.6223 ± 0.0002 | 0.787 ± 0.02* | 0.869 ± 0.05* | 0.675 ± 0.01 | 0.77 ± 0.03 |
| Total Chl | 300.16 ± 28.12 | 242.78 ± 11.84 | 56.11 ± 3.4 | 285.35 ± 12.44 | 150.88 ± 2.07** | 60.81 ± 0.51** | 44.94 ± 0.51* | 47.61 ± 2.86 | 47.15 ± 1.14 | 44.61 ± 1.51* |
| CHL:CAR | 4.07 ± 0.36 | 4.62 ± 0.735 | 0.79 ± 0.048 | 44.24 ± 1.29*** | 22.12 ± 0.27*** | 0.298 ± 0.014*** | 0.186 ± 0.003** | 0.248 ± 0.01*** | 0.28 ± 0.01*** | 0.3 ± 0.03*** |

Carotenoid and chlorophyll contents are presented as $\mu\text{g/g}$ fresh weight. Data are means ± SE ( $n = 3$ ). Chl, chlorophyll; CAR, carotenoid; ND, not detected as no peak was discernible on the chromatogram. Student's *t*-test was used to determine the significant differences in the metabolites between *Nps1* and AV. Values in bold indicate where significant differences have been found. The P values $< 0.05$ , $< 0.01$ and $< 0.001$ were indicated by one, two or three asterisks respectively.

The comparison of the ripening pattern of backcrossed progeny with parental *Nps1* revealed no perceptible differences in duration to attain MG, and subsequent transition to BR, and RR (Figure 2C-D). The on-vine ripening of *Nps1*\* and AC fruits post-RR stage was nearly similar, discounting any role of *Nps1*\* on the regulation of fruit senescence. The mutation was also confirmed by Sanger sequencing in *Nps1*\*/*Nps1*\* plants in both AC and AV backgrounds (Figure 2E-H).

### *phot1* loss of function alleles exhibit reduced carotenoid levels in fruits

To ascertain whether the enriched carotenoids level in *Nps1\** was specific to its dominant-negative action, we generated CRISPR/Cas9-edited loss of function alleles by targeting the first exon of *phot1* gene. Out of eight editing events in T_0_ plants, five were recovered to homozygosity in T_1_ progeny (Figure 3). Among five alleles, PHOT1 protein got prematurely truncated in two alleles at T100Cys* (*phot1^CR1^*) and D117Cys* (*phot1^CR3^*), while three alleles had in-frame deletion of 50 (*phot1^CR2^*), 96 (*phot1^CR4^*) and 21 (*phot1^CR5^*) amino acids. In the T_2_ generation, *phot1^CR2^, phot1^CR3^,* and *phot1^CR5^* became Cas9-free and showed no off-target effects (Supplemental Figure 2). All the alleles showed reduced chloroplast accumulation and avoidance in leaf (Supplemental Figure 3), which confirms that both *phot1* and *phot2* redundantly regulate the chloroplast relocation response.

**Figure 3.**
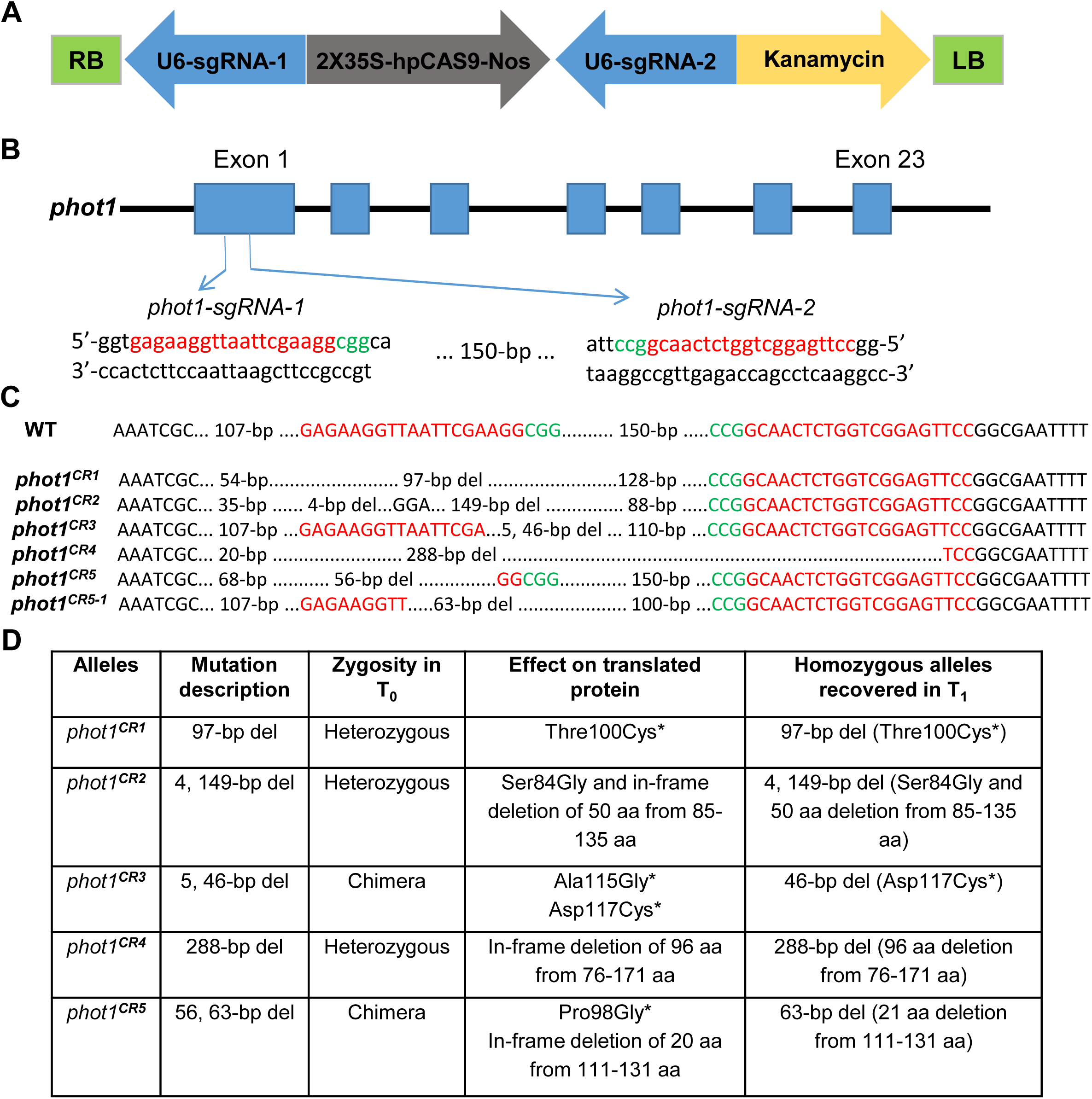
The CRISPR/CAS9-mediated editing in the tomato *phot1* gene. **A**, Schematic representation of binary vectors used in this study. *phot1-sgRNA-1* & *2:* Arabidopsis *U6* promoter and the gRNA sequence; 2X 35S-hpCAS9-Nos: 2X *CaMV35S* promoter sequence, hpCas9: human-codon optimized SpCas9, Nos: nos terminator; Kanamycin: the kanamycin resistant marker expression cassette; RB: right border of T-DNA; LB: left border of T-DNA. **B**, Schematic view of sgRNA1 and sgRNA2 target sites in the *phot1* gene. Boxes indicate exons, red color indicates 19-20 bp target sequences and green indicates the protospacer-adjacent motif (PAM). **C**, Sequence alignment of the target regions. The wild type sequence is shown at the top with the target sequence in red and the PAM in green. Nucleotide variations at the targets of T_0_ mutant lines, ‘del’ -base deletion. **D**, The table shows the alleles identified in T_0_, effect of editing on *phot1* gene and protein sequences and the recovered alleles in T_1_ generation.

Though all the alleles showed delayed attainment of the MG stage similar to *Nps1*\*, it was one-day shorter (7-days). Post-MG, however unlike *Nps1*, the transition to BR was shorter in all alleles (to a lesser extent in *phot1^CR1^*), and BR to RR attainment was marginally extended in all except *phot1^CR1^* than AV (Supplemental Figure 4A). Importantly, opposite to *Nps1*\*, in all the alleles, the fruit colour at RR stage was lighter indicating that the carotenoids accumulation was reduced (Supplemental Figure 4B). Particularly lycopene, phytoene, phytofluene, β-carotene, and α-carotene were significantly lower than AV in all the alleles (Table 3). The opposing influence on carotenoid accumulation in *Nps1*\* vis a vis *phot1^CR1-5^* lines entails that enhancement of carotenoids in *Nps1*\* was specific to its dominant-negative action. We then characterized in detail the influence of *Nps1*\* on phytohormones, metabolome, and proteome of fruits including the transcript levels of selected genes vis-à-vis AC.

**Table 3.** Carotenoid content in the red ripe fruits of homozygous *phot1^CR^* lines in T_2_ generation.

| Metabolite<br>($\mu\text{g/gFW}$) | AV | *phot1<sup>CR1</sup>* | *phot1<sup>CR2</sup>* | *phot1<sup>CR3</sup>* | *phot1<sup>CR4</sup>* | *phot1<sup>CR5</sup>* |
| --- | --- | --- | --- | --- | --- | --- |
| Phytoene | 59.914±4.98 | 31.559±1.74** | 43.815±2.825* | 42.302±4.594* | 26.091±0.526* | 27.609±1.745*** |
| Phytofluene | 7.775±0.4319 | 4.986±0.3166*** | 5.624±0.333* | 5.5±0.7* | 4.242±0.471*** | 3.799±0.359*** |
| Lycopene | 42.374±6.148 | 20.871±1.253*** | 22.036±1.205** | 23.505±2.371* | 17.01±1.709*** | 18.227±2.26** |
| β-carotene | 3.416±0.156 | 2.191±0.113*** | 2.541±0.089*** | 2.503±0.244* | 1.971±0.121*** | 1.887±0.114*** |
| α-carotene | 5.441±0.596 | 2.995±0.178** | 2.652±0.077** | 2.289±0.124** | 1.808±0.165** | 1.851±0.133** |
| Lutein | 1.297±0.124 | 2.897±0.523* | 3.445±0.285*** | 2.63±0.211*** | 2.58±0.198*** | 2.504±0.123** |
| Total<br>carotenoids | 128.8±9.53 | 66.479±1.225* | 74.935±2.774*** | 68.714±6.396* | 51.009±4.461*** | 53.268±1.794*** |

Carotenoid contents are presented as $\mu\text{g/g}$ fresh weight. Student's *t*-test was used to calculate the significant differences between AV and *phot1<sup>CR</sup>* lines. Data are means of 6-8 biological replicates ± SE. The P values < 0.05, < 0.01 and < 0.001 were indicated by one, two or three asterisks respectively.

### *Nps1\** fruits show reduced respiration and ethylene emission

The onset of ripening in tomato fruits is strongly associated with increased ethylene emission, called a system II response. Though ripening duration from MG to RR in AC and *Nps1*\* was nearly similar, the ethylene emission was highly subdued in *Nps1*\* (Figure 4). In tomato, system II ethylene is mainly contributed by elevated expression of ethylene biosynthesis genes *1-aminocyclopropane-1-carboxylate synthase 2* and *4* (*acs2*, *acs4), 1-aminocyclopropane-1-carboxylic acid oxidase 1,* and *3 (aco1, aco3)* (Barry et al., 2000). Compared to AC, *acs2* and *acs4* transcripts were distinctly higher in *Nps1*\* at MG and BR but were nearly similar at RR (barring *acs2* which is lower) (Supplemental Figure 5). Conversely, expression of *aco1* and *aco3* was highly reduced in *Nps1* (barring *aco1* at BR). Seemingly, the reduced expression of *aco* may have contributed to lower ethylene emission from *Nps1*\* fruits. Similar to ethylene, *Nps1*\* fruits also emitted less CO_2_ signifying reduced respiration than AC (Figure 4). *Nps1*\* RR fruits were more firm and had higher °Brix. Though fruit’s acidity differed in AC and *Nps1\** at MG and BR, it was similar at RR (Supplemental Figure 6). Contrasting to *Nps1*\*, in RR fruits of *phot1^CR^* alleles, °Brix and pH was akin to AV (Supplemental Figure 4C-D).

**Figure 4.**
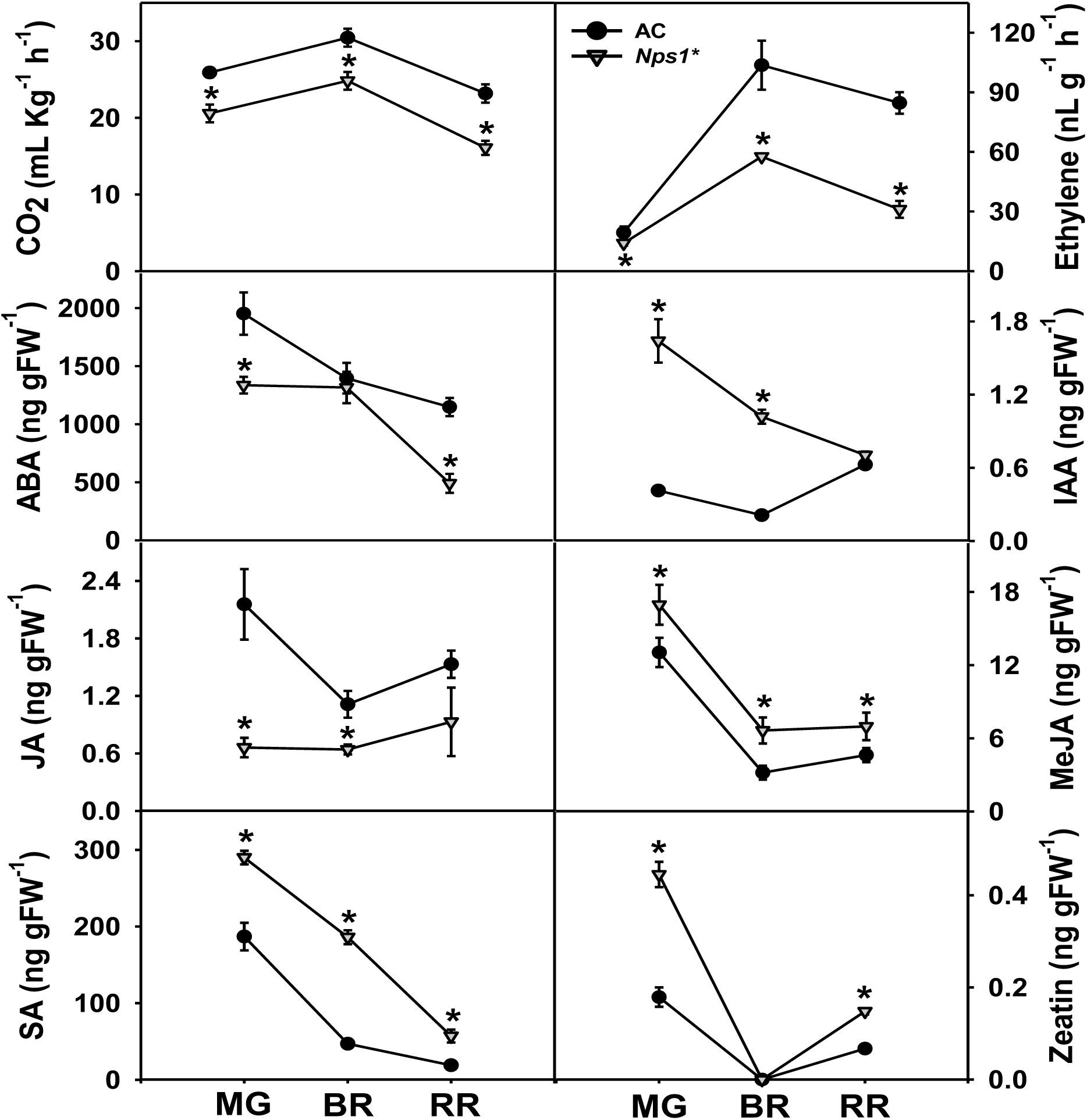
Rate of respiration and hormonal levels in ripening fruits of AC and *Nps1*\*. Data are means ± SE (n=5), *P≤0.05.

### Hormonal balance is altered in *Nps1*\*

Based on analyses of seedling phototropism, it is surmised that phototropins affect the transport of auxins (Fankhauser and Christie, 2015). Barring auxin, the influence of phototropins on other phytohormones is not known. We examined whether, in addition to ethylene, other phytohormones were also modulated by *Nps1*\* (Figure 4). The IAA level was higher in *Nps1*\* at MG and BR, which is consistent with reported antagonistic action of ethylene and auxin. The higher levels were also observed for MeJA, SA, and zeatin (barring BR). In contrast, ABA (barring BR) and JA levels were lower in *Nps1*\* than AC. It is apparent that the influence of *Nps1*\* was not restricted to ethylene, it also influenced the overall hormonal homeostasis of fruits.

### Several MEP and carotenogenesis pathway genes are upregulated in *Nps1*\*

Since *Nps1\** fruits exhibit high lycopene content, it is reasonable to assume that this may be associated with the upregulation of carotenoid biosynthesis genes. The geranylgeranyl diphosphate (GGPP) needed for carotenoid biosynthesis is derived from the methylerythritol-4-phosphate (MEP) pathway. While the first gene of MEP pathway, *deoxy-xylulose 5-phosphate synthase* (*dxs*) had lower expression in *Nps1*\*, the other genes, *deoxy-xylulose 5-phosphate reductase* (*dxr*)*, 4-hydroxy-3-methyl but-2-enyl diphosphate reductase* (*hdr*)*, isopentenyl diphosphate isomerase 5g* (*idi5g*) (BR, RR) and *geranylgeranyl pyrophosphate synthase 2 (ggpps2)* (MG) were upregulated in *Nps1\** (Figure 5).

**Figure 5.**
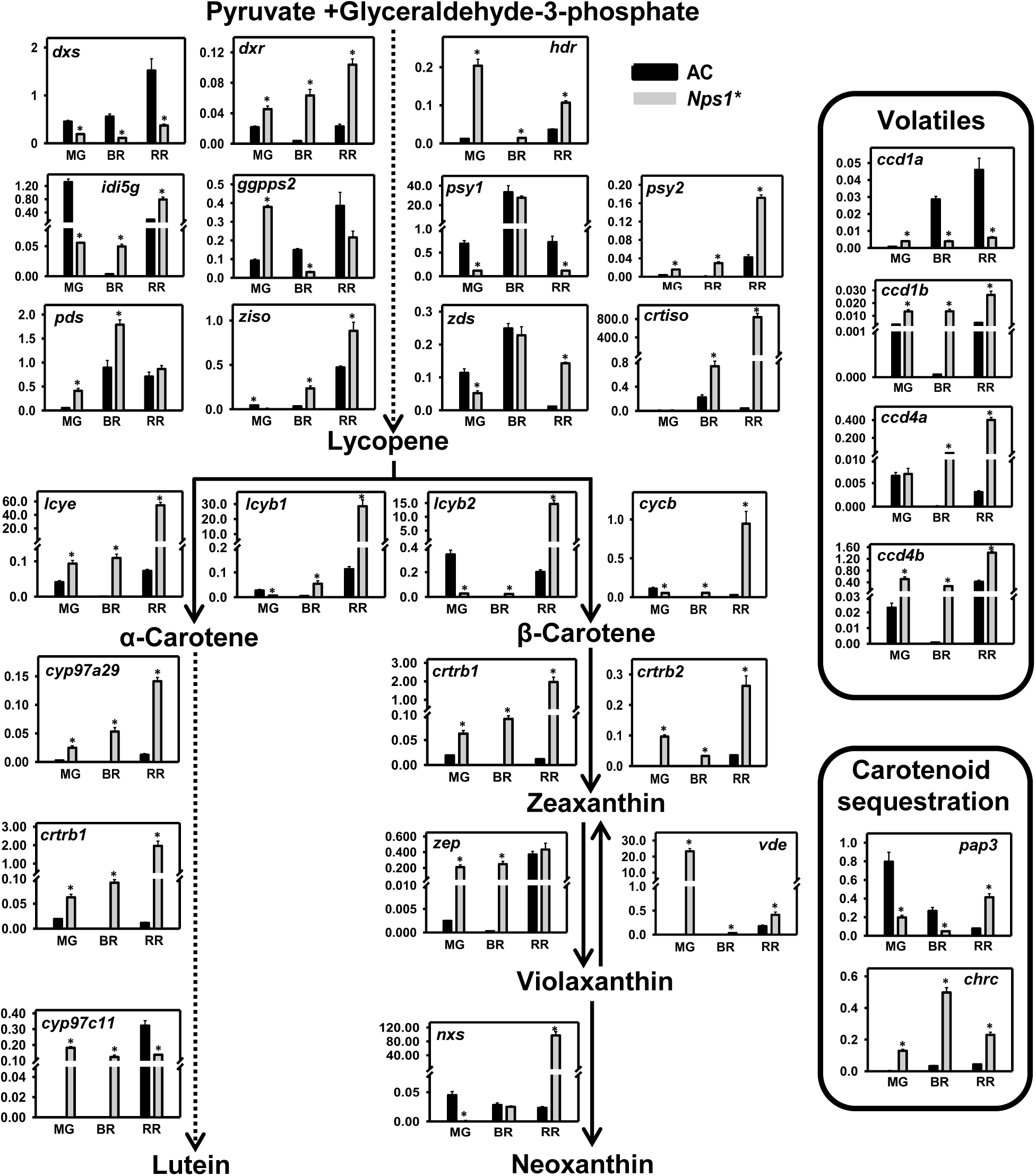
Relative expression of genes mediating carotenoid biosynthesis, sequestration, and carotenoid derived volatile formation in ripening fruits of AC and *Nps1*\*. Dotted arrows indicate multiple steps in the carotenoid biosynthesis pathway. The genes mediating carotenoid sequestration and volatile formation are enclosed in separate rectangles. The y-axis indicates relative expression of genes obtained after normalization with *β-actin* and *ubiquitin*. Data are means ± SE (n=3), *P ≤ 0.05.

Downstream to MEP pathway in tomato, GGPP conversion to phytoene by phytoene synthase1 (PSY1) is considered as a first committed step for fruit-specific carotenogenesis. Surprisingly, *psy1* expression in *Nps1\** was considerably lower (barring BR). Conversely, *psy2*, a leaf carotenogenesis gene, was upregulated in *Nps1\**. Nevertheless, the expression of downstream genes, *phytoene desaturase* (*pds*), *ζ-carotene isomerase* (*ziso*), *ζ-carotene desaturase* (*zds*) and *carotenoid isomerase* (*crtiso*) leading to lycopene were higher in *Nps1\** (barring *pds* at RR, *ziso* at MG, *zds* at BR). Seemingly, desaturation steps are more crucial than phytoene formation for enhanced lycopene levels in *Nps1\** fruits.

Lycopene is converted to β-carotene by the action of lycopene β-cyclases. Remarkably, the expression of all four lycopene cyclases was higher in *Nps1\** at BR and RR stages. *Nps1\** also showed stage-specific higher expression of xanthophyll and lutein pathway genes. The expression of the *chromoplast-specific carotenoid-associated protein* (*chrc),* a carotenoid sequestration gene, was considerably higher in *Nps1\**. Likewise; another sequestration gene-*pap3* was also upregulated but only at RR. Out of 4 *carotenoid cleavage dioxygenases* (*ccd)* genes examined, three (*ccd1b, ccd4a, ccd4b*) were upregulated in *Nps1*\*.

Given the low carotenoid content in *phot1^CR^* alleles, we examined whether the expression of few key genes regulating carotenogenesis differs from *Nps1*\* (Supplemental Figure 7). Though the *dxs* expression varied at MG, BR stages in Cas9-free *phot1^CR2^, phot1^CR3^,* and *phot1^CR5^*, it was higher at RR stage than AV. In *phot1^CR2^, phot1^CR3^,* and *phot1^CR5^* alleles, while *psy1* expression was lower at MG (except *phot1^CR5^*) and BR stages, it was higher at RR stage than AV. The *psy2* expression was either similar or lower than that of AV at all stages (except at MG in *phot1^CR5^*). The expression of chromoplast-specific *lycopene-β-cyclase* (*cycb*) in *phot1^CR2^, phot1^CR3^,* and *phot1^CR5^* alleles was lower at all stages during ripening than AV (except at MG in *phot1^CR3^*). Ostensibly, the influence of *Nps1*\* on carotenogenesis genes was distinctly different from *phot1^CR^* alleles.

### *Nps1*\* influences gene expression of phototropins and other ripening regulators

The influence of *Nps1*\* on stimulation of carotenoid biosynthesis genes seems to be indirect, as unlike other photoreceptors, phototropins are not localized in the nucleus. Nonetheless, *Nps1*\* ripening fruits showed high transcript levels of both *phot1* and *phot2* (Figure 6). Considering that *Nps1*\* showed delayed onset of ripening and had reduced ethylene synthesis, we examined whether *Nps1*\* affected the expression of key transcriptional regulators regulating the above responses. Consistent with reduced ethylene emission, the expression of *ethylene response factor 6 (erf6)* was lower in *Nps1*.* In contrast, the expression of a key ripening regulator-*ripening inhibitor (rin)* and *tomato agamous-like 1 (tagl1),* was higher in *Nps1*\* (barring BR stage). Conversely, the expression of other ripening regulators-*Nonripening (nor)*, *fruitful 1 (ful1),* and *fruitful 2* (*ful2)* was lower in *Nps1*\* at MG and BR stages. *Nps1*\* also showed lower expression of genes related to chloroplast numbers viz., *golden 2-like (glk2),* and *Arabidopsis pseudo response regulator 2-like (aprr2)* at BR and RR stages. The other carotenoid biosynthesis regulatory genes, *Cys-rich zinc finger domain-containing protein (Or), phytochrome interacting factor1 (pif1)* showed higher expression at MG and BR and lower expression at the RR stage in *Nps1*\*. Ostensibly, the influence of *Nps1*\* was not limited to carotenoid biosynthesis genes, it influenced phototropins, and genes involved in fruit ripening diversely.

**Figure 6.**
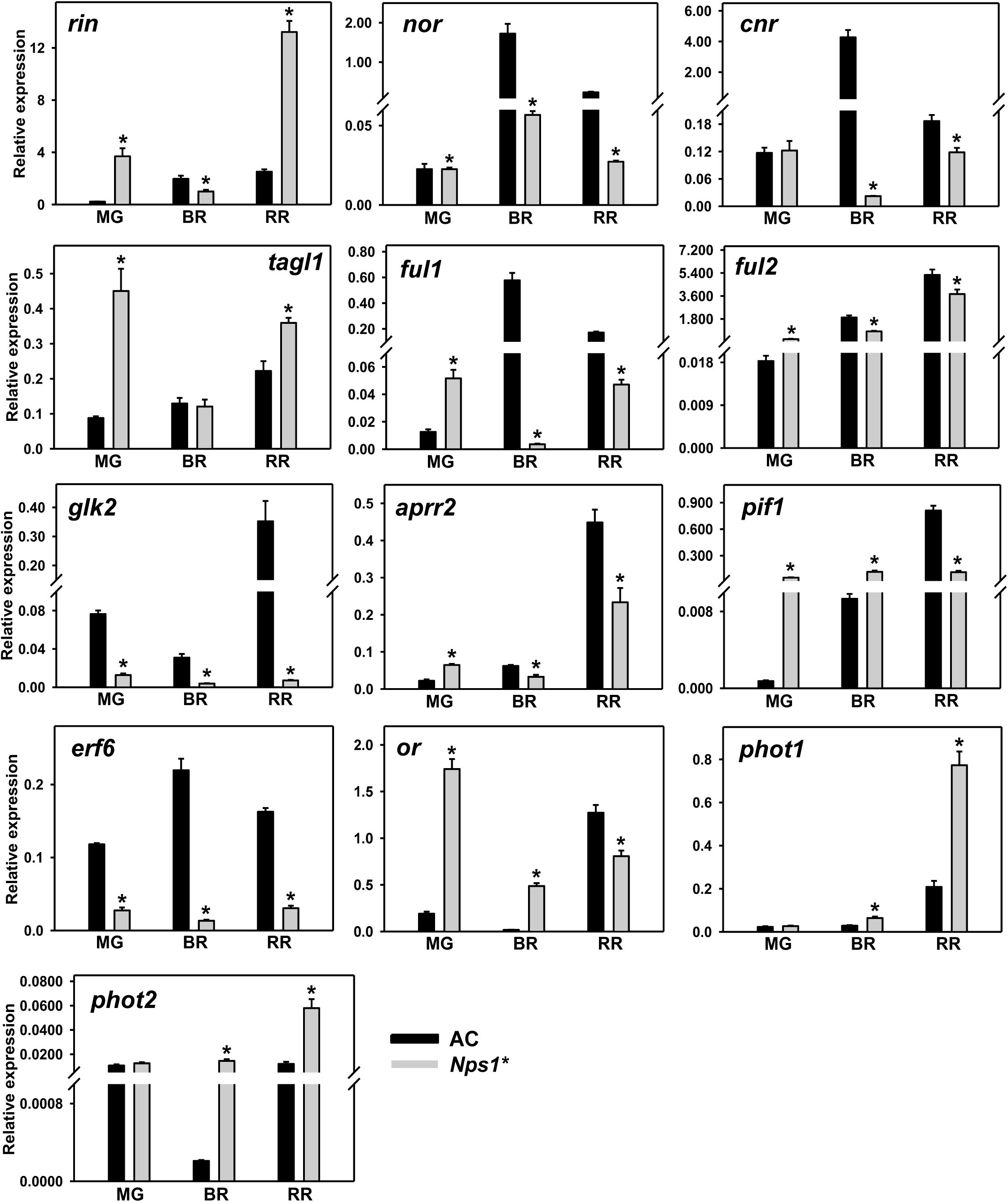
Relative expression of ripening regulators, modulators of carotenogenesis, and *phototropin* genes in ripening fruits of AC and *Nps1*\*. The graphs depict data obtained after normalization with *β-actin* and *ubiquitin*. Data are means ± SE (n=3), *P ≤ 0.05.

### *Nps1*\* upregulates fruit aminome and sugars

Similar to regulating carotenogenesis, *Nps1*\* also influenced the metabolome of ripening fruits in a varied manner. The principal component analysis highlighted that the metabolome of *Nps1*\* was distinct from AC, and the most prominent differences were at the RR stage (Supplemental Figure 8). The majority of amino acids and sugars were high in *Nps1*\* RR fruits (Figure 7; Supplemental Figure 9; Supplemental Table 3). The sucrose level was higher in *Nps1*\* which seems to contribute to the higher Brix. Consistent with reduced CO_2_ emission from RR fruits, the levels of citrate and aconitate contributing to the first two steps of the TCA cycle were lower in *Nps1*\*. In general, the levels of organic acids contributing to the TCA cycle were low in *Nps1*\*.

**Figure 7.**
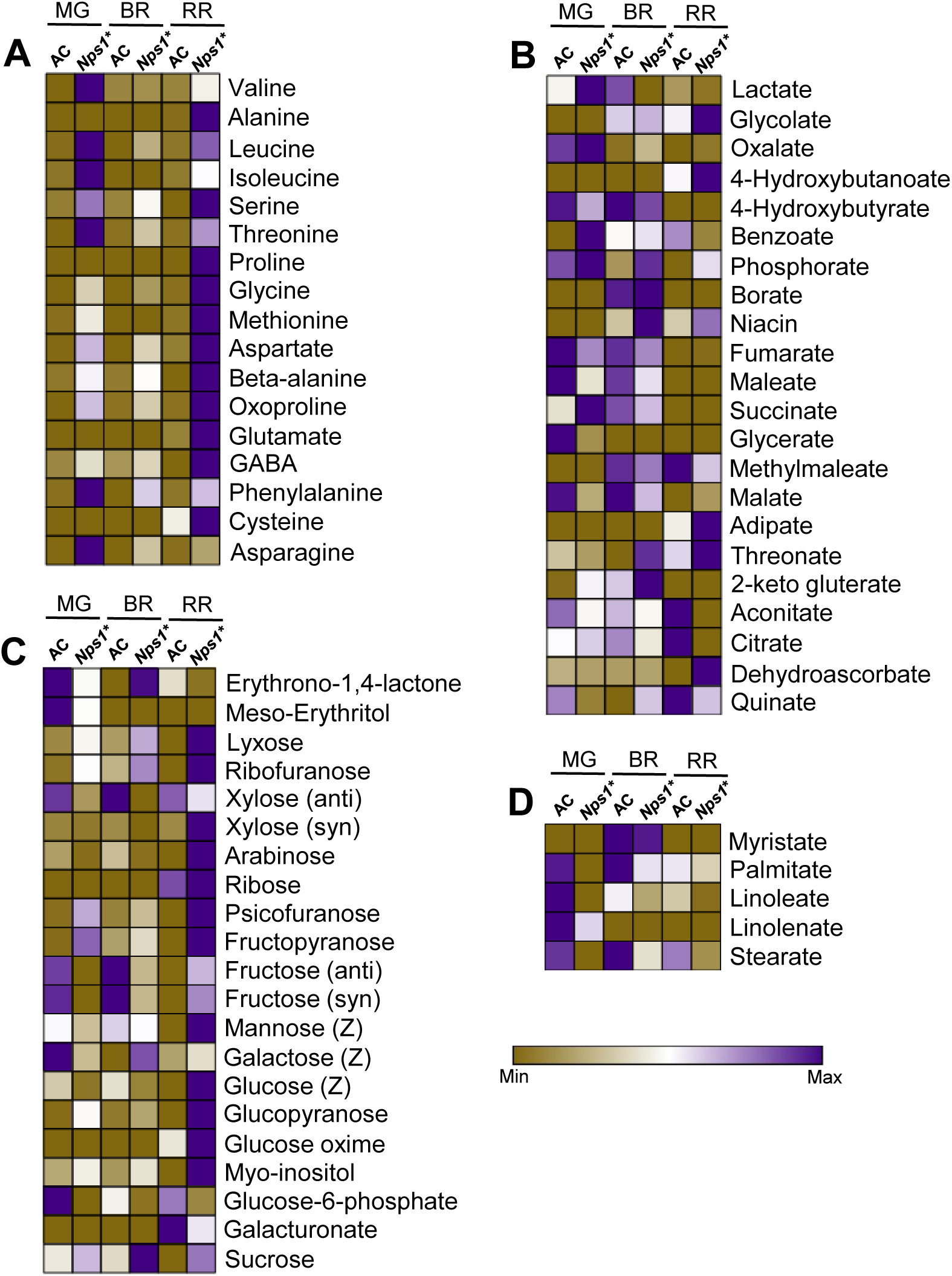
Primary metabolites level in ripening fruits of AC and *Nps1*.* Only significantly different metabolites (≥1.5 fold) are depicted in heat maps. Data are means ± SE (n=5), P≤0.05. **A**, Amino acids. **B**, Organic acids. **C**, Sugars. **D**, Fatty acids.

### Levels of lycopene-derived 6-methyl 5-hepten-2-one are high in *Nps1*\*

The distinct aroma of tomato is contributed by a myriad of compounds contributed from several metabolic pathways including carotenoids. Analogous to primary metabolites, the volatile profile of *Nps1*\* was distinct from AC (Figure 8; Supplemental Figure 10, Supplemental Table 4). Among the volatiles detected in *Nps1*\* and AC, the majority were derived from the phenolics pathway, followed by lipid, terpenoid, and phenylpropanoid pathways. In general, *Nps1*\* showed reduced levels of most volatiles, and only a few were upregulated at one or more stages. Among the carotenoid derived volatiles, only 6-methyl 5-hepten-2-one (MHO) and acetophenone were detected. Consistent with higher lycopene in *Nps1*\* RR fruits, the level of aroma compound, MHO (lycopene derivative) was 2.7-fold high. Intriguingly, the other carotenoid derived volatiles, geranial, pseudoionone, β-ionone, or geranyl acetone were not detected. Among the odor and taste contributing volatiles, methyl salicylate, phenylethanol, hexanal, and *p*-menth-1-en-9-al were lower in *Nps1*\* fruits. Overall, while *Nps1*\* upregulated several metabolites at the RR stage, a similar effect is not perceptible on volatiles, except MHO linked directly with lycopene levels.

**Figure 8.**
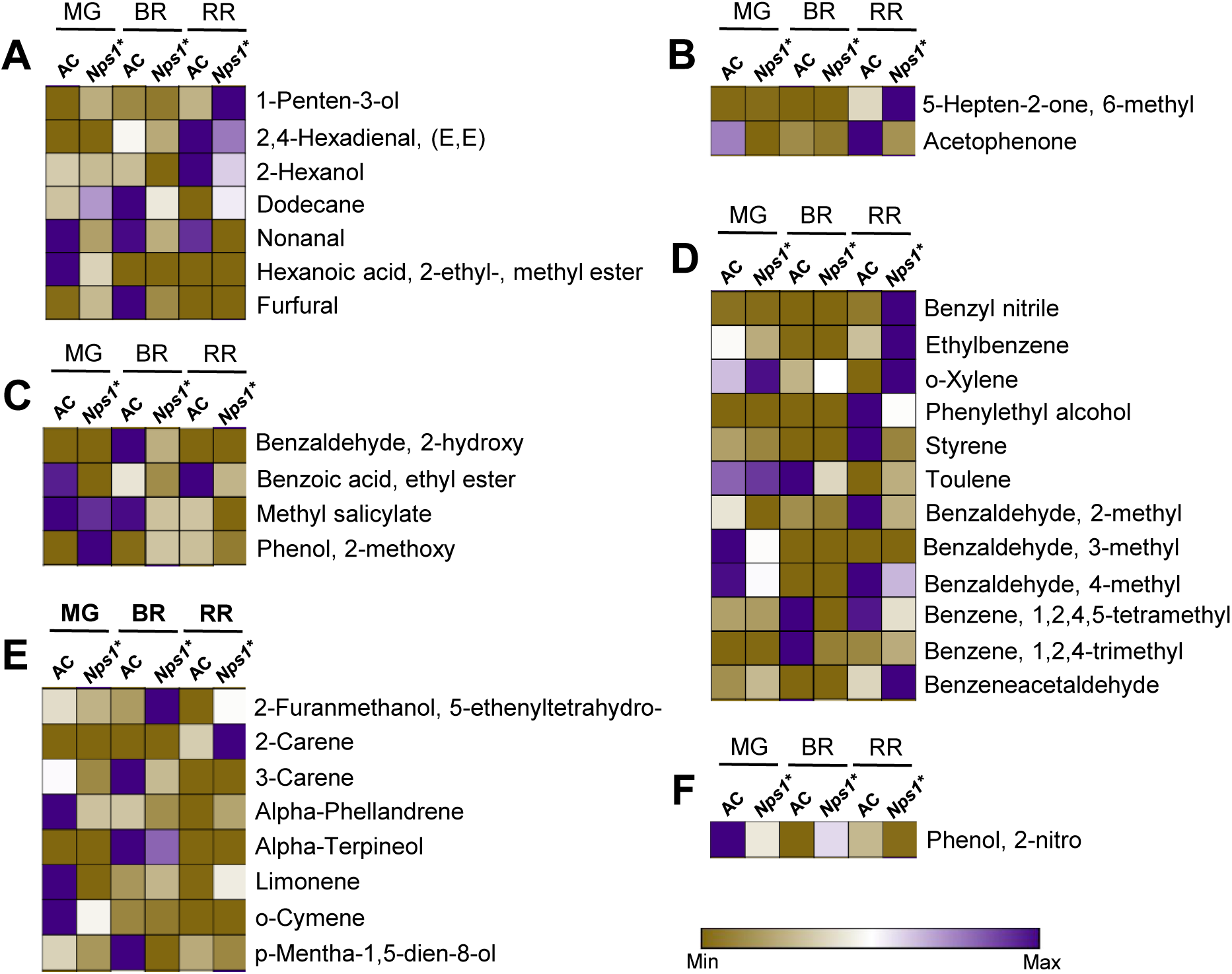
Volatile analysis in ripening fruits of AC and *Nps1\**. **A**, Lipid derived. **B**, Open-chain carotenoid derived. **C**, Phenylpropanoid derived. **D**, Phenolic derived. **E**, Terpenoid derived. **F**, Amino acid derived volatiles. Only significantly different metabolites (≥1.5 fold) are depicted in heat maps. Data are means ± SE (n=8), P≤0.05.

### *Nps1*\* upregulates abiotic stress-related proteins

To decipher a link between elevated lycopene levels in *Nps1*\* and proteins contributing to carotenogenesis, we profiled the proteome of AC and *Nps1*\* fruits. The proteome (with two or more peptide hits) comprised of ca. 2336 (MG), 2375 (BR) and 2318 (RR) proteins (Supplemental Figure 11). The label-free proteome quantification using Scaffold identified several upregulated (MG-190, BR-167, RR-139) or downregulated (MG-113, BR-126, RR-92) proteins in *Nps1\** compared to AC (Supplemental Table 5). Both *Nps1*\* and AC also had unique proteins exclusively associated with them, as no single peptide hits of these proteins were detected vice versa. These proteins may be present in lower abundance in *Nps1*\* and AC thus precluding their detection. The unique proteins identified in *Nps1\** (MG-131, BR-95, RR-72) and in AC (MG-177, BR-117, RR-98) varied in stage-specific fashion (Supplemental Table 6). Relatively, ripening fruits of *Nps1*\* displayed fewer unique proteins but lost several proteins compared to AC.

The functional annotation of the differentially expressed proteins revealed distinct differences between *Nps1*\* and AC at all ripening stages (Supplemental Figure 12). Both up- and down-regulated proteins in *Nps1*\* were distributed across the full spectrum of functions. Of interest was a greater upregulation of the abiotic stress category except for the BR stage, and greater downregulation of the biotic stress category in *Nps1*\*. Consistent with enhanced amino acid levels, the amino acid metabolism category had greater downregulation in *Nps1*\*.

### *Nps1*\* upregulates carotenoid sequestration proteins

Considering that *Nps1*\* shows a substantially high amount of carotenoids in RR fruits, it is reasonable to expect that proteins contributing to carotenoid biosynthesis too are upregulated. Contrarily, proteome profiling revealed the opposite trend. Oddly carotenogenesis proteins viz. GGR, DXS, PSY1, ZISO, ZDS, CRTISO, while detected in AC (RR), were not detected in *Nps1*\* (Figure 9; Supplemental Table 5-6). Moreover, levels of plastidic HDR involved in the formation of carotenoid precursors, isopentenyl diphosphate and dimethylallyl diphosphate, were 5-fold down-regulated (RR) in *Nps1*.* Only a lone protein, isopentenyl diphosphate isomerase (IDI1) involved in dimethylallyl diphosphate synthesis was identified in *Nps1*.* Notably, two proteins involved in the sequestration of carotenoids were upregulated in *Nps1*\*. Prominently, the levels of CHRC, a protein associated with high lycopene content (Kilambi et al, 2013), were 4.7-fold higher in *Nps1*\*. Likewise, CHRD, a carotenoid sequestration protein (Leitner-Dagan et al., 2006a), was exclusively detected in *Nps1*\* RR fruits. Tangibly, the high carotenoids in *Nps1\** fruits seem to be interlinked with a higher amount of sequestration proteins.

**Figure 9.**
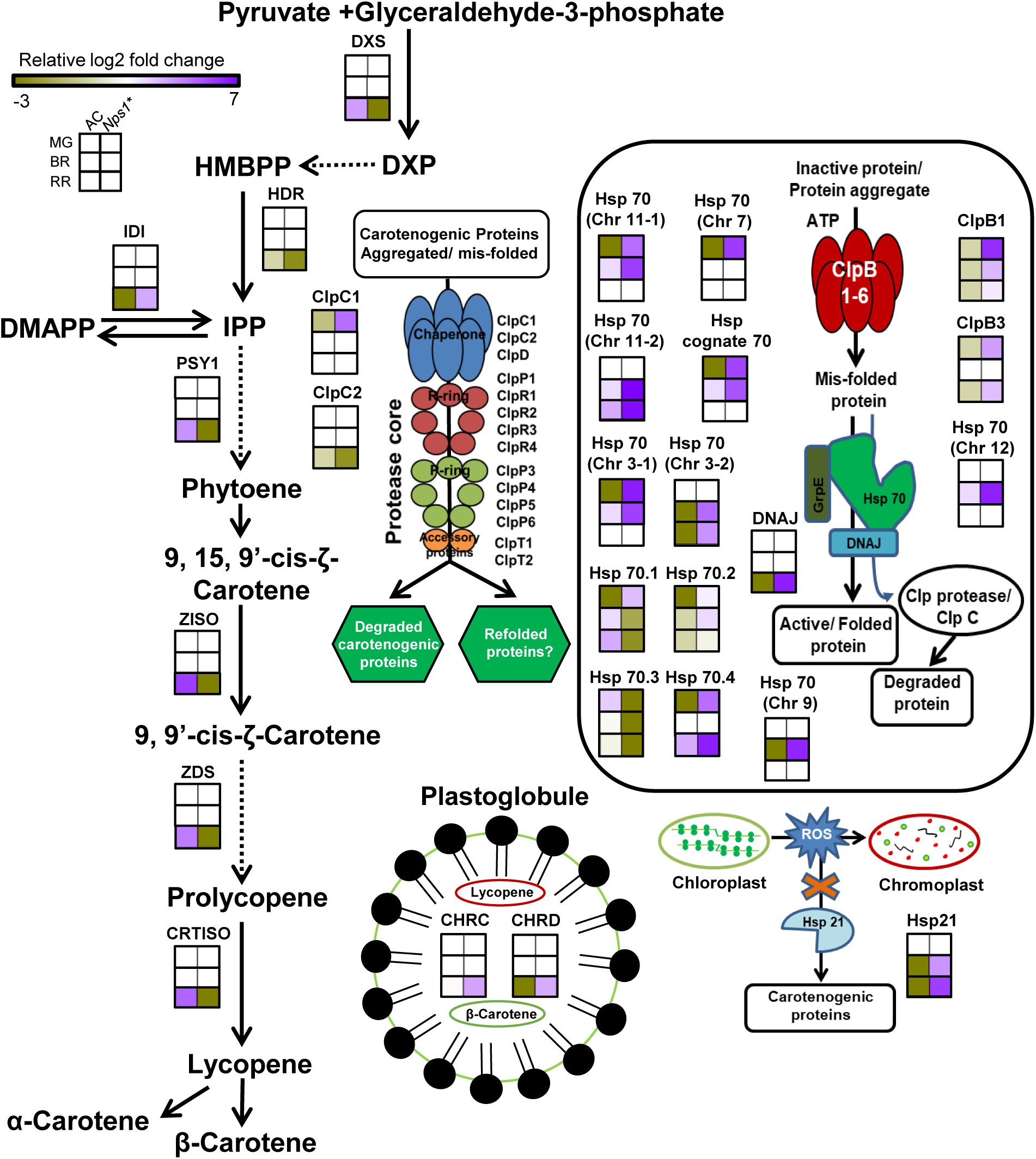
Schematic representation of relative abundances of proteins involved in carotenoid biosynthesis, sequestration, protein folding, degradation and heat shock proteins in AC and *Nps1\** fruits at different stages of ripening. Heat maps were generated using normalized log2 fold changes for differentially expressed proteins. For proteins detected only in *Nps1\** or in AC, the missing values were replaced with 1/10 of the lowest value in the dataset and the heat maps were generated with normalized log2 abundances. Carotenogenic proteins that are aggregated or misfolded can be folded into their active confirmation by Clp B protease system in the presence of ATP together with Hsp 70 and DNAJ proteins (Maurizi and Xia 2004). Alternatively, the above misfolded or aggregated proteins can be degraded by Clp C/Clp protease system or facilitated for refolding (Pulido et al., 2016; Nishimura and van Wijk 2015; Andrea et al., 2018). CHRC and CHRD are plastid lipid associated proteins that can be localized to the plastoglobule and assist in sequestration of carotenoids (Kilambi et al., 2013; Leitner-Dagan et al., 2006a and b). Chloroplast to chromoplast conversion involves generation of reactive oxygen species (ROS) that may induce either degradation or inactivation of carotenogenic proteins. This action of ROS can be prevented by Hsp 21, which may thus protect carotenogenic proteins from degradation (Neta Sharir et al., 2005).

### Molecular chaperones maintaining protein stability are abundant in *Nps1*\*

Comparative proteome profiling of *Nps1*\* highlighted that it influenced a broad spectrum of proteins thus affecting overall protein homeostasis of ripening fruits. Particularly, the functional category of proteins of abiotic stress was upregulated in *Nps1*\*. Most abiotic stress proteins protect the functional proteome by being molecular chaperones, mediating proper folding of proteins, and preventing protein degradation. Consistent with this, *Nps1***\*** fruits showed striking upregulation of molecular chaperones belonging to Hsps during ripening (Supplemental Figure 13, Supplemental Table 7). At all stages of ripening, *Nps1*\* displayed higher levels of several Hsps than AC (MG-17; BR-17; RR-11 Hsps). While many of these Hsps showed stage-specific upregulation, few were high at two or more stages. Specifically, Hsp21 was abundantly present only in *Nps1*\*. The levels of Hsp70.2 in *Nps1*\* was also higher in ripening fruits. Despite Hsp21 protein being present only in *Nps1*\*, the transcript levels of *Hsp21* were low in *Nps1*\* and that of *Hsp70*.2 declined in ripening fruits (Supplemental Figure 14). The above protein versus transcript levels mismatch indicates the operation of posttranscriptional regulation of Hsp21 and Hsp70.2. Likewise, proteins assisting protein-folding such as peptidyl-prolyl isomerase (PPI), 14-3-3 proteins, protein disulfide isomerase (PDI), and methionine sulfoxide reductase (MSR) too were upregulated in *Nps1*\* (Supplemental Figure 13). However, an important chaperone, OR was below detectable level in both *Nps1*\* and AC. The protection of proteins was further aided by the absence of proteases in *Nps1*\*, particularly at the RR stage. Conversely, several proteases were detected only in AC RR fruits (Supplemental Table 6).

The proteome profile of *Nps1*\* showed altered levels of several Clp components, with some purported to regulate the activity of carotenogenic proteins (Figure 9; Supplemental Figure 13; Supplemental Table 7). The abundance of ClpB3 protease (Solyc12g0426060) in *Nps1*\* was 6-fold high during early ripening, whereas ClpB1 (Solyc03g115230.3.1) was upregulated throughout ripening. While downregulation of ClpC2 (Solyc12g0426060) was observed only at RR, the level of ClpC1 (Solyc03g0118340) in *Nps1*\* and AC was nearly similar during ripening. Akin to *Hsp21*, in *Nps1*\* the transcript levels of *ClpC1* and *ClpB3* lacked correlation with respective protein levels during ripening, indicating a posttranscriptional control (Supplemental Figure 14).

## Discussion

The capability to enhance the carotenoid levels in fleshy fruits is an important biotechnological target (Giuliano, 2017). In tomato, analysis of high pigment and photoreceptor mutants including transgenics has demonstrated a causal link between light signaling and carotenogenesis. It alludes that manipulation of photoreceptors and associated signaling partners can be an effective means for boosting carotenoids levels in fruits. In this study, we show a novel function for plant photoreceptor phototropin1, wherein a dominant-negative mutant-*Nps1* considerably enhances the carotenoid levels in tomato fruits, whereas the loss of function alleles reduce the carotenoid content.

### Altered hormonal balance seemingly delays attainment of MG in *Nps1*

The role of photoreceptors-cryptochromes and phytochromes, in tomato fruit development has been largely inferred by comparative phenotyping of mutants with wild type. Relatively few reports are available for functional assay of fruit-localized photoreceptors. Alba et al. (2000) demonstrated the operation of functional phytochromes in detached MG tomato fruits, as red light-stimulated lycopene accumulation in these fruits was reversed by far-red light. Likewise, both phototropin1 and phototropin2 are functional in MG fruits, as these fruits display chloroplast accumulation and avoidance respectively. Contrarily, *Nps1* seems to be compromised in functional phototropins, as blue-light failed to elicit chloroplast movement in mutant fruits.

Unlike tomato *high pigment*-*hp1* and *hp2* mutants that have deep-green colored MG fruits, *Nps1*\* MG fruits were pale-green colored with reduced chlorophyll levels. Emerging evidence indicates that chlorophyll accumulation in tomato fruits is mediated by phytochromes (Bianchetti et al., 2018; Gupta et al., 2014). Reduced chlorophyll levels in *Nps1\** fruits construe an ancillary role of phototropins in this process. Similar to phytochrome mutants (Gupta et al., 2014), *Nps1*\* fruits too show delayed attainment of the MG stage. While the above observations allude a linkage between phototropins and phytochromes, it is limited to the phase preceding the MG stage. Post-MG stage, unlike phytochrome mutants that display delayed progression to BR and RR stages (Gupta et al., 2014), duration to BR and RR stages was similar in *Nps1*\* and AC.

While the involvement of phytohormones in tomato fruit development is well documented, the knowledge regarding the crosstalk between light and phytohormones in the pre-MG phase is fragmentary. Similar to *Nps1*\*, an auxin-resistant (Balbi and Lomax, 2003) and a NO-overproducing mutant (Bodanapu et al., 2016) show prolonged pre-MG phase implying hormonal participation. In tomato fruits, enhanced ethylene emission coupled with high ABA and low IAA levels led to the early attainment of the MG stage (Sharma et al., 2020; Liang et al., 2020). Conversely, reduced ABA and ethylene levels and high auxin levels seem to delay the attainment of the MG stage in *Nps1*\*. Collectively, it can be construed that *Nps1*\* delays attainment of the MG stage by influencing hormonal balance.

### *Nps1*\* dominant-negatively enhances carotenogenesis in tomato

In ripening tomato fruits, carotenoid synthesis does not obligatorily require light, nonetheless light is necessary for maximum carotenoid production (Raymundo et al., 1976; Gupta et al., 2014). On-vine ripened fruits are exposed to natural daylight, which is perceived by multiple photoreceptors such as UVR8, cryptochromes, phototropins, and phytochromes. In general, deficiency of photoreceptors such as UVR8 (Li et al., 2018), or cryptochromes (Giliberto et al., 2005; Liu et al., 2018; Fantini et al., 2019), or phytochromes (Gupta et al., 2014; Bianchetti et al., 2018) lead to reduced carotenoid levels. Conversely, constitutively active phytochromeB2-overexpressor line with a modified GAF domain stimulated the carotenoid level (Alves et al., 2020). Unlike these photoreceptors, *Nps1*\* while impairing normal phototropins functioning such as chloroplast movement, considerably stimulates carotenoid accumulation. It seems that stimulation of carotenoid levels by *Nps1*\* is linked with its constitutively active nature, as genome-edited loss of function *phot1* alleles displayed reduced carotenoids. The reduced carotenoid levels in gene-edited lines signify that a functional phot1 is necessary for normal fruit carotenogenesis, while constitutively active phot1 in *Nps1*\* likely exacerbates carotenoids accumulation. The recreation of tomato Arg495His substitution in Arabidopsis phot1 protein (Arg472His) conferred constitutively active kinase domain in the absence of light (Petersen et al., 2017). Likewise, an equivalent mutation in *Chlamydomonas* phot (Arg210His) also exhibited constitutive kinase activity *in vitro* (Aihara et al., 2012). Collectively above reports support the notion that constitutively active phot1 in *Nps1*\* may have stimulated carotenoid accumulation.

At times, the influence of dominant-negative mutations such as *Nac-Nor* or dominant mutation *Nr* may vary in different genetic backgrounds (Wang et al., 2020; Elkind et al., 1990; Rodríguez et al., 2010; Lanahan et al., 1994). Such background influence was not conspicuous for *Nps1*\*, as in two different genetic backgrounds, it dominant-negatively stimulated carotenoids levels. The influence of *Nps1*\* was not confined to carotenoids, it also influenced the metabolic and volatile shifts associated with ripening. Higher amino acids and sugars coupled with reduced CO_2_ emission indicated its effect on a broad spectrum of responses. A similar broad-spectrum influence was seen in volatile profiles, nonetheless higher levels of MHO in *Nps1*\* were in agreement with higher lycopene levels (Tieman et al., 2017). In tomato *cry2* mutant, a decrease in the lycopene is associated with an increase in fatty acids (Fantini et al., 2019). Contrarily, high lycopene in *Nps1*\* fruits is accompanied by a decrease in fatty acids, indicating a possible inverse relationship between these two pathways. It remains possible that a decrease in fatty acids may be related to diversion to the carotenoid pathway.

### Ethylene emission and carotenogenesis are not linked in *Nps1*\*

Compared to other plant photoreceptors that influence gene expression by associating with transcription factors in the nucleus, phototropins are membrane-bound (Liscum et al., 2020), thus may influence the carotenoids and metabolome indirectly. Lamentably, only a little information is available about constituents linking phototropin triggered signal to altered photoresponses, while those leading to gene expression are not known. Alternatively, phototropins can regulate the carotenoid levels in ripening fruits by altering the hormonal balance. It is known that an asymmetric auxin gradient forms on the activation of phototropins that precedes the manifested hypocotyl curvature (Esmon et al., 2006). Considering that formation of the above auxin gradient is associated with transcription of auxin associated genes (Esmon et al., 2006), it remains plausible that alteration of auxin levels by *Nps1*\* may be responsible for higher carotenoids accumulation. In tomato, evidence indicates that post-MG phase carotenoid accumulation is modulated by auxin-ethylene interaction, wherein high auxin suppresses ethylene action and reduces carotenoid accumulation (Su et al., 2015). Contrarily, in *Nps1*\* though auxin level is high and ethylene emission is reduced, yet fruits accumulate a high amount of carotenoids.

The role of other hormones in upregulating carotenoids levels in *Nps1*\* is also equivocal. *Nps1*\* fruits accumulate a much higher level of carotenoids than the JA-overexpressor line (Liu et al., 2012), ABA deficient mutants (Galpaz et al., 2008) or transgenics (Sun et al., 2012). Therefore, it seems unlikely that reduced ABA, JA, or high MeJA levels in fruits contributed to carotenoid stimulation. Although several studies have shown a strong correlation between ethylene emission and fruit-specific carotenoid biosynthesis (Lanahan et al., 1994; Oeller et al., 1991), reduced ethylene emission not necessarily leads to reduced carotenoids. While light signaling defective *hp1* mutant accumulates high carotenoids, the ethylene emission was less than wild type (Kilambi et al., 2013). Similarly, in *Nps1*\* reduced ethylene emission does not lessen carotenoid accumulation. In entirety, though *Nps1*\* strongly affects the hormonal balance of tomato fruits, its influence on carotenoid accumulation is not strictly dependent on it.

### *Nps1*\* upregulates several MEP and carotenogenesis-related genes

Though the identity of the signal chain components emanating from *Nps1*\* is arcane, it upregulated key ripening regulators, *rin,* and *tagl1*, which are known to promote carotenogenesis. The above cohort also included *Or* and *pif1* known to modulate carotenogenesis. Reduced expression of *glk2* and *aprr2* conformed that high carotenoids in *Nps1*\* is unrelated to chloroplast numbers. Considering that expression of *nor* and *ful1, 2* were lower, they seem not to participate in *Nps1*\* mediated carotenogenesis.

Though phototropins mediate chloroplast movements, they seem not to affect photosynthesis or chlorophyll levels in vegetative tissue (Gotoh et al., 2018). Consistent with this, no discernible difference in chlorophyll levels was observed in leaves of *Nps1*\* and AC. Nonetheless, reduced chlorophyll levels in *Nps1*\* MG fruits suggest that phototropin action in fruit is distinct from leaves. While the onset of fruit ripening triggers chlorophyll catabolism (Kilambi et al., 2017), in parallel it stimulates chloroplast to chromoplast transformation. The chromoplast-localized carotenoid biosynthesis is distinct from leaves as it is mediated by chromoplast specific *psy1* and *cycb*. Its precursor-GGPP is also derived from the plastid-localized MEP pathway, which in general has no specificity towards chromoplasts except GGPPS2, which is localized in chromoplasts (Ament et al., 2006). Since chemical inhibition of MEP or carotenoid pathway leads to reduced carotenoid levels in fruits (Rodriguez-Concepcion, 2010), evidently coordination between these two pathways ensures the supply of precursors. While it remains to be determined how this coordination is accomplished, it is suggested that the activity of key enzymes of these two pathways, DXS1 and PSY1 seem to have some coordination, though their transcript levels are not commensurate (Fraser et al., 2007; Lois et al., 2000).

Unlike chloroplasts, after the onset of ripening, major flux from the MEP pathway is primarily directed towards carotenoids, as the levels of other metabolites dependent on MEP pathway such as side chain of chlorophylls, and phylloquinones are not essential for chromoplast functioning. In *Nps1*\*, while *dxs1* and *ggpps2* genes of the MEP pathway are downregulated, the intermediate genes-*dxr*, *hdr*, and *idi1* are upregulated. Considering that *hdr* and *dxr* are single-copy genes and their overexpression boosts the carotenoids levels (Botella-Pavia et al., 2004; Estévez et al., 2001), their upregulation can contribute to elevated carotenoids in *Nps1*\*. Proteome profile revealed that in *Nps1*\*, DXS and DXR proteins are reduced and IDI1 is exclusively detected. Since IDI1 disruption reduces carotenoid content in fruits (Pankratov et al., 2016), it is plausible that high IDI1 in *Nps1* also directs the flux towards carotenoids. It appears that in *Nps1*\*, a complex modulation of multiple MEP pathway constituents determines precursor supply towards the carotenoids pathway.

In ripening tomato fruits, the first committed step of carotenoid formation is mediated by a fruit-specific PSY1 (Fray and Grierson, 1993) which is distinct from leaf-specific PSY2 (Fraser et al., 1999), with no overlap in their functions. Consequently, mutants defective in PSY1 fail to accumulate carotenoids in fruits (Fraser et al., 1999). Similar to the MEP pathway, barring *psy1,* the expression of other genes upstream of lycopene was upregulated in *Nps1*\*. While higher expression of *psy1* is associated with stimulation of lycopene levels (Fraser et al., 2007), the lowered expression does not inevitably lower the lycopene level. In *phot1^CR^* loss of function alleles too, though *psy1* expression was high at RR stage, the lycopene content was low. A negative correlation between *psy1* expression and lycopene content was observed in the *hp1* mutant (Kilambi et al., 2013) and *DET1-*underexpression lines of tomato (Enfissi et al., 2010). Oddly, in *Nps1*\* the expression of *psy2* was higher. Current evidence indicates that *psy2* does not contribute to fruit specific carotenogenesis (Fraser et al., 1999; Fantini et al., 2013). However, it may have a hitherto undiscovered role in fruits.

While PSY1 is involved in condensation of GGPP to phytoene, other genes downstream to it leading to lycopene formation catalyze structural modifications of C-40 phytoene, comprising of desaturation and isomerization. Emerging evidence indicates that the stimulation of the lycopene level in tomato fruits is not limited to *psy1*, other intermediary genes also contribute to this process (Fantini et al., 2013). Consistent with this, overexpression of *pds* in tomato enhances lycopene and β-carotene in the fruits (McQuinn et al., 2018). The normal function of *zds* is also critical to lycopene accumulation, as its suppression by RNAi reduces lycopene level (McQuinn et al., 2020). Higher expression of intermediary genes in *Nps1*\* signifies that these genes can also contribute to high lycopene levels. Nonetheless, for most carotenogenesis genes, the correlation was amiss between transcript levels and the respective protein abundances in *Nps1*\*. The above lack of correlation signifies that enhanced carotenoids in *Nps1\** likely arise from the maintenance of their function rather than their relative abundance. Current evidence favors that the protection of carotenogenic proteins is also an essential part of higher carotenoids synthesis in tomato fruits (D’Andrea and Rodriguez-Concepcion, 2019).

Similar to upstream genes, the genes downstream to lycopene also showed higher expression levels. Nonetheless, unlike genes upstream of lycopene, the higher expression of these genes showed little correlation with the levels of downstream carotenoids. The carotenoid catabolism is mediated by CCDs releasing volatiles and bioactive apocarotenoids such as β-cyclocitral and strigolactone (Hou et al., 2016). Among CCDs, CCD1 degrades lycopene releasing volatiles such as MHO (Vogel et al., 2008; Ilg et al., 2014). Markedly, the higher expression of *ccd1b* gene in *Nps1*\* seems to correlate with high MHO levels.

### Levels of carotenoid sequestration proteins are higher in *Nps1*\*

Considering that increased gene expression often does not lead to a commensurate increase in carotenoids, it entails that the posttranscriptional steps such as mRNA modification (Tan et al.,2017; Li et al., 2018), protein abundance and stability (Yazdani et al., 2019; D’Andrea et al., 2018), also govern carotenoids accumulation. Despite genetic and transgenic intervention to decipher the regulation of carotenogenesis, the knowledge about the contribution of proteins and enzymes in fruits is far from complete. Notwithstanding the high expression of several carotenogenic pathway genes and also phototropins in *Nps1*\*, yet none of the corresponding proteins were detected in proteome profiles. It is plausible that unlike MEP pathway proteins, carotenogenic pathway proteins may be of low abundance, therefore escape detection. More than the abundance, the enzyme activity of respective proteins is more important for controlling flux into the pathway. Considering that increase in PSY1 activity despite low expression in *hp1* mutant elevates carotenoids levels in fruits (Cookson et al., 2003), a similar activation of pathway enzymes may underlie the elevation of carotenoids in *Nps1*\*.

Unlike carotenogenic proteins, with increased carotenoid levels it is expected that carotenoid sequestration proteins too exhibit a comparable increase. Such an association between high carotenoid levels and increased levels of carotenoid sequestration proteins has been observed in tomato fruits (Kilambi et al., 2013, 2017; Nogueira et al., 2013). Since the levels of CHRC and CHRD are elevated in *Nps1*\*, it is reasonable to assume that a higher amount of these proteins contributes to high carotenoid content. Contrasting to carotenogenic proteins, the increased level of CHRC protein was also associated with increased transcript levels.

### High Aminome seems to be linked to reduced levels of TCA cycle proteins

Parallel to the increase in carotenoids, *Nps1*\* considerably influences the aminome of fruits, increasing the levels of several amino acids. Considering that these amino acids are derived from glycolysis and TCA cycle intermediates, their elevation would reduce flux into the TCA cycle. Reduced CO_2_ emission from *Nps1*\* fruits also implied a subdued TCA cycle. Consistent with this, levels of citrate, aconitate, and α-ketoglutarate levels were low in *Nps1*\* fruits. Proteome profile also revealed that several proteins related to the TCA cycle or its derivatives were affected in *Nps1*\*. Since the level of citrate catabolizing enzyme aconitase was higher in *Nps1*\*, it may rapidly transform citrate to isocitrate. At the next step, the low level of isocitrate dehydrogenase in *Nps1*\* may cause lower α-ketoglutarate (2-oxoglutarate) levels. Besides, due to reduced levels of α-ketoglutarate dehydrogenase in *Nps1*\*, a substantial part of α-ketoglutarate maybe diverted to amino acids. Consistent with this, levels of amino acids derived from α-ketoglutarate such as glutamate and proline were substantially high in *Nps1*\*.

In consonance with reduced flux in the TCA cycle, the proteome profiles revealed an increased abundance of proteins involved in the amino acid synthesis. The reduced flux in the TCA cycle seems to also involve pyruvate, which is converted by pyruvate dehydrogenase (PDH) to acetyl CoA, an immediate precursor for the TCA cycle. Ostensibly reduced abundance of PDH isoforms in *Nps1*\* fruit during ripening may limit the availability of pyruvate for the TCA cycle. In corollary, increased pyruvate levels boost levels of pyruvate-derived amino acids like alanine, valine, leucine, which were abundant in *Nps1*\*. Increased malate level in *Nps1*\* can also contribute to pyruvate. In *Nps1*\* reduced levels of two isoforms and absence of one isoform of malate dehydrogenase, may lead to a build-up of malate. Since *Nps1*\* also shows an abundance of few isoforms of malic enzyme (ME) catalyzing the conversion of malate into pyruvate (Osorio et al., 2013), this could be an additional source for pyruvate. In parallel, the MEP pathway may also be stimulated by the increased level of pyruvate, thus boosting the carotenoids level in *Nps1*\*.

### Upregulation of protein protection seems to be related to higher carotenogenesis in *Nps1*\*

Remarkably, the proteome profile of *Nps1*\* revealed a substantial enrichment of proteins belonging to molecular chaperones, protein folding, and proteinase inhibitors. Emerging evidence indicates that modulation of stability of proteins by chaperones and proteases plays an important role in maintaining the activity of rate-limiting enzymes of MEP pathway such as DXS and DXR (Flores-Perez et al., 2008; Pulido et al., 2016, 2017). The reduction in Clp and Hsp70.2 activities by mutations or by transgenic means leads to higher accumulation of enzymatically active forms of DXS and DXR (Flores-Perez et al., 2008; Pulido et al., 2016) and high carotenoids (D’Andrea et al., 2018; D’Andrea and Rodriguez-Concepcion, 2019). Consistent with this, the high abundance of ClpB3 protease and Hsp70.2 and low abundance of ClpC1 and ClpC2 proteases in the *Nps1*\* may help in the reactivation of DXS and other carotenogenic proteins (D’Andrea and Rodriguez-Concepcion, 2019).

The role of chaperones in chromoplast formation and enhanced carotenoids levels have been revealed by *Hsp21* (Neta-Sharir et al., 2005; Carvalho et al., 2012), and *Or* (Yazdani et al., 2019) overexpression in tomato fruits. While OR protein was below detection limits, Hsp21 was abundant throughout ripening and present solely in *Nps1*\*. Therefore, it is plausible that high levels of Hsp21 contribute to enhanced lycopene levels in *Nps1*\*. Considering the importance of chaperones and protein folding in maintaining enzyme activity of MEP pathway and carotenogenic proteins, the enrichment of these proteins in *Nps1*\* is suggestive of a similar role. It is plausible that these proteins in *Nps1*\* provide an overall protective microenvironment that allows more efficient functioning of carotenogenic proteins leading to higher levels of carotenoids.

It remains to be determined how *Nps1*\* influences broad spectrum responses encompassing increased carotenogenesis, metabolic and proteomic shifts. Within the constraints of limited knowledge about phototropin signaling, we can only make a conjecture. Phototropins function as light and temperature sensors and loss of this sensing capability by dominant-negative mutation may create a stressful situation. While other regulatory mechanisms may alleviate the stress in vegetative tissues, during fruit ripening absence of such compensation may amplify the stress. Since this stress is within, it is construed as abiotic stress, leading to the upregulation of abiotic stress-related proteins and protein protection mechanisms.

In summary, our results highlight that phototropins play an important role in the regulation of carotenoid biosynthesis during fruit ripening. Besides bringing a new photoreceptor regulating carotenogenesis in tomato, our study unfolds a potential biotechnological tool to enhance the carotenoid levels in fruits. The dominant-negative feature of *Nps1*\* makes it an attractive tool for enhancing carotenoids levels in heterozygous fruits, thus has considerable value for the hybrid industry. Our work adds to current impetus to generate tomato with health-promoting carotenoids and advances the repertories available for making more nutritious tomato.

## Materials and methods

### Plant material and growth conditions

Tomato (*Solanum lycopersicum)* cultivar Ailsa Craig (LA2838, TGRC, Davis, Ca), Arka Vikas (IIHR, Bengaluru, India) and *non-phototropic seedling 1* mutant, *Nps1* (Sharma et al., 2014) were used. Seeds germination and low-fluence blue light-induced phototropic curvature analysis of parents, and back-crossed progeny was carried out as described in Sharma et al (2014). After the phototropic screening, seedlings were transferred to white light (100 µmol m^-2^s^-1^) for one week. Thereafter plants were transferred to greenhouse maintained at 28±1°C during the day and ambient temperatures at night.

Reciprocal crosses were made between wild type (AC and AV) and *Nps1* plants. The BC_1_F_1_ plants were self-pollinated to generate BC_1_F_2_ plants. The progeny was backcrossed again to raise BC_2_F_2_ progeny. The homozygous BC_2_F_2_ (*Nps1*: *Nps1*) plants were then advanced to BC_2_F_3_ and BC_2_F_4_ progeny. During backcrossing, the zygosity of progeny for *Nps1* at each stage was identified by low-fluence phototropism as described above and CELI endonuclease assay as described below.

### Mismatch Endonuclease screening

In parallel with phototropic screening, the homozygosity and heterozygosity of mutant, AV, AC, and backcross progeny were carried out using a CELI endonuclease assay (Mohan et al., 2016). In brief genomic DNA was extracted from the leaves of respective plants using an in-house protocol (Sreelakshmi et al., 2010). The *phot1* gene encompassing the mutated base (guanine-1484-to-adenine transition) was PCR-amplified from genomic DNA using unlabeled M13 tailed primers (Supplemental Table 8). The PCR products after heteroduplex formation were digested with CELI (Mohan et al., 2016). The digested fragments were resolved using agarose gel electrophoresis and visualized for the presence of the mutation. The homozygosity and heterozygosity of plants were also confirmed by Sanger sequencing of PCR-amplified *Phot1* gene products (Macrogen, Korea).

### Chloroplast relocation

The MG fruits after harvest were dark-adapted for 12 h. The fruits were cut into two identical halves under green safe light. For chloroplast accumulation, after obtaining the dark position of chloroplasts, the slices were exposed to low-fluence blue light (5 µmol/m^2^/s) for 30 min. Thereafter, slices were exposed to high-fluence (100 µmol/m^2^/s) blue light for 30 min to induce chloroplast avoidance. The positioning of chloroplasts was observed in a Zeiss confocal microscope as described in Sharma et al. (2014).

Chloroplast movement in tomato leaf was monitored by the measurement of red light transmittance through leaf discs at 25°C using a microplate reader (Biotek, Synergy HT) as described by Wada and Kong (2011). Briefly, from overnight dark-adapted leaves 0.3-0.4 cm size discs of (8 discs/line) were punched and overlaid on 0.5% (w/v) agar under green safe light in a 96 well plate. Thereafter the transmittance of light through leaf discs monitored at 1 min at 660 nm. The irradiation schedule consisted of 30-min measurement in darkness (for the baseline), followed by weak blue light (∼3.2 µmol/m^2^/s for accumulation response) irradiation for (80 minutes, thereafter high-intensity blue light (∼80 µmol/m^2^/s for avoidance response) exposure for 60 minutes. The darkness baseline value was subtracted from weak/strong light values to obtain the Δ transmittance.

### Generation of gene-edited lines of *phot1* by CRISPR-Cas9

Double guide RNAs (sgRNA) for the *phot1* gene (Solyc11g072710) with GC content between 40-60% were selected using the CRISPR-P web tool (http://cbi.hzau.edu.cn/crispr; Lei et al., 2014). Each sgRNA was folded *in silico* by RNAfold server (http://mfold.rna.albany.edu/?q=mfold/RNA-Folding-Form2.3), to check dangling spacer and stable hairpin structure. The gRNAs chosen to develop CRISPR vectors are Phot1-sgRNA-1: GAGAAGGTTAATTCGAAGG and Phot1-sgRNA-2: GGAACTCCGACCAGAGTTGC.

The human codon-optimized CAS9 with 2×35S CaMV promoter and Nos terminator was amplified from pAGM4723 plasmid (Addgene# 49772) using *Kpn*I forward and *Pac*I reverse primers (Supplemental Table 9). The amplified products were cloned into the pCAMBIA2300 vector. The gRNA expression cassette was developed through overlap extension PCR. Briefly, the AtU6 promoter, sgRNA scaffold (with target region), and AtU6 terminator (Nekrasov et. al, 2013) was synthesized (Macrogen, Korea). Two long primers of 130 bp for sgRNA expression cassette were annealed through overlap extension PCR. Using *Pac*I forward and *Spe*I reverse primers, the Phot1-sgRNA-1 and with *Kpn*I forward and *Kpn*I reverse primers, the Phot1-sgRNA-2 expression cassettes were amplified and cloned into a pCAMBIA2300-CAS9 vector. All the fragments were gel eluted, ligated, and confirmed. The final vector pCAMBIA2300-CAS9-Phot1 was verified by sequencing and the primers information is listed in Supplemental Table 9.

The pCAMBIA2300-CAS9-Phot1 vector was transferred into *Agrobacterium* strain LBA4404 using the freeze-thaw method. *Agrobacterium tumefaciens*-mediated transformation of the standard processing tomato cultivar, AV was performed according to Van Eck et al. protocol (2006). Transgenic plants were recovered in the medium supplemented with kanamycin and transferred into the soil. Genomic DNA was isolated according to Sreelakshmi et al. (2010) from 8-12 week-old plants. The CRISPR/Cas9 induced mutations were screened by amplifying a region of about 546 bp surrounding both the gRNAs in the *phot1* gene using *Taq* DNA polymerase (Supplemental Table 9). To detect mutations, the amplified mutated and the wild-type DNA fragments were separated by agarose gel electrophoresis. Sanger sequencing was performed after cloning the PCR product into InsTAclone PCR Cloning Kit (Thermo Scientific). Six individual clones/editing event were sequenced using the M13 forward primer (Supplemental Table 9). The confirmed lines for editing were advanced to T_1_ generation and the homozygous mutants were identified by PCR followed by sequencing. The confirmed homozygous edited lines were advanced to T_2_ generation and Cas9-free lines were identified. These Cas9-free homozygous lines were further characterized for fruit characteristics as described below.

Off-targets for the *phot1* sgRNAs were identified using CRISPR-P and CRISPOR web tools (http://cbi.hzau.edu.cn/crispr; http://crispor.tefor.net) (Lei et al., 2014; Haeussler et al., 2016). A list of 33 potential off-targets was predicted for the target locus with up to 4 mismatches. Further, based on high cutting frequency distribution (CFD) off-target score (Haeussler et al., 2016) seven potential off-targets were selected (four for Phot1-sgRNA-1; three for Phot1-sgRNA-2; Supplemental Table 10). The genomic DNA surrounding the potential off-target locus was amplified using specific primers (Supplemental Table 9) and the PCR products were analyzed by Sanger sequencing.

### Fruit ripening

For fruit ripening study, AC, AV, *Nps1*,* and *phot1^CR1-5^* plants were grown in three randomized blocks (each block constitutes one biological replicate) as described previously (Kilambi et al., 2013; Supplemental Figure 15). The flowers were tagged at anthesis for chronological monitoring of fruit development. The time point of attainment of different ripening stages viz, MG, BR, and RR were identified. Fruits were harvested at respective stages, flash-frozen in liquid nitrogen, and stored at -80°C until further use as described previously (Kilambi et al., 2013).

### Estimation of carotenoids, transcripts, hormones, metabolites, °Brix, firmness and respiration rate

The carotenoid and chlorophyll levels were determined from leaves and fruits using the Gupta et al. (2015) and Wellburn (1994) protocols respectively. RT-PCR analysis and ethylene emission estimation were carried as described in Kilambi et al. (2013). For RT-PCR analysis, RNA extraction from the fruits, cDNA conversion, DNAase treatment, and analysis was carried out as described previously. RT-PCR was carried out using *β*-*actin* and *ubiquitin* as reference genes (Supplemental Table 11). The hormone estimation and metabolite analysis were carried out as described by Bodanapu et al. (2016). Metabolites were extracted from the fruits of AC and *Nps1\** (BC_2_F_2_ homozygous; five biological replicates each) and identified by GC-MS. Sugar content, pH, and firmness of fruits were determined as described by Gupta et al. (2014). For respiration rate, tomato fruit was enclosed an air-tight container and a time course of CO_2_ emission was recorded using a CO_2_ Sensor (Vernier Carbon Dioxide Gas Sensor, CO_2_-BTA) at 30-sec interval for 10 min at 23°C, 75% RH and calculated using preinstalled Graphical AnalysisTM 4 software.

### Proteome profiling

Protein isolation, separation, and digestion were carried out as described earlier by Kilambi et al. (2013, 2016, and 2017). Proteins (70 µg) were first separated by one dimensional SDS-PAGE. Thereafter in-gel digestion was carried out using trypsin, and peptides were extracted. After desalting by C18 spin columns (Thermo Scientific), peptides were aliquoted, dried, and stored at −80°C.

### Mass spectrometry, data analysis, and label-free quantification

Peptides were analyzed using Easy nanoLC-II coupled with LTQ Velos Pro mass spectrometer (Thermo Scientific). Peptides (350 ng) were separated using a setup consisting of a trap column (Integrafrit 10µm × 2.5cm, 5µm, C18 New Objective, USA) and a Biobasic C18 picofrit column (7µm × 10cm, 5µm, New Objective, USA) by solvent A (95/5-water/ACN with 0.1% (v/v) formic acid) and solvent B (95/5-ACN/water with 0.1% (v/v) formic acid) at a flow rate of 350 nL/min. For this fractionation, a gradient of 5%-30% B (0-100 min), 30%-95% B (101-103 min), 95% B hold (104-106 min), 95%-5% B (107-113 min), 5% B hold (114-118 min) was employed for a 118 min run time. All the other mass spectrometric parameters used for this fractionation were exactly as mentioned in Kilambi et al. (2016).

Data analysis was done using Sorcerer (version 5.1 release, Sage-N Research). *S. lycopersicum* iTAG3.2 proteome sequence (ftp://ftp.solgenomics.net/tomato_genome/annotation/ITAG3.2_release/ITAG3.2_proteins.fasta, downloaded on July 18, 2017, 35673 sequences and 12210990 residues) was used as the database. The searches were done with Sequest with the parameters- a peptide mass tolerance and fragment mass tolerance of 5 ppm and 0.8 Da respectively, trypsin as the protease, with a maximum of two allowable missed cleavages, carbamidomethylation of cysteine as fixed and oxidation of methionine as variable modifications. Peptides were filtered for high confidence (95% protein and peptide probabilities, assigned through Protein prophet algorithm; Nesvizhskii et al., 2003) and these were used for assigning protein IDs. Peptides with FDR of 1% were selected, which was estimated based on the number of decoy hits using a local FDR database. The mass spectrometry proteomics data are available via ProteomeXchange (Vizcaíno et al., 2014) with the dataset identifier PXD018150. Label-free quantitation was carried out using Scaffold software (version 4.7.5, Proteome Software) as described previously in Kilambi et al. (2016).

### Extraction of volatile compounds and their identification

The volatile compounds from the fruits (8 biological replicates) were extracted as described in Rambla et al. (2016). The extracted volatiles were adsorbed to a 50 μm divinylbenzene/Carboxen/Polydimethylsiloxane (DVB/CAR/PDMS) SPME fiber assembly (Supelco) for 20 min at 50°C under continuous agitation. The fiber was inserted into GC 7890A (Agilent Technologies, Palo Alto, CA, USA) coupled with MS Pegasus 4D with Time of Flight as the detector (LECO Corporation, MI, USA). The volatiles were desorbed at 250°C for 1 min. The volatiles were then separated using the following program: 5 min at 45°C with a linear ramp of 5°C/min to 250°C and held at 250°C for 5 min. Both the injector and detector temperatures were set at 260°C and 250°C respectively. Helium was used as carrier gas at a flow rate of 1.5 mL min^-1^. Ionization energy (EI) of 70 eV was used for mass spectrometry detector, with a source temperature of 250°C, a scan range of 35-600 m/z and the mass spectra were recorded at a scan speed 2 scans/sec.

The raw data were processed by ChromaTOF software 2.0 (Leco Corporation, USA) and further analyzed using the MetAlign software package (Lommen and Kools, 2012; www.metalign.nl) with a signal to noise ratio of ≥ 2, for baseline correction, noise estimation, alignment and extraction of the ion-wise mass signal. The mass signals that were present in less than three biological replicates were discarded. The Metalign results were processed with MSClust software for the reduction of data and compound mass extraction (Tikunov et al., 2012). The mass spectra extracted by MSClust were searched in NIST MS Search v 2.2 software for the identification of compound names within the NIST (National Institute of Standard and Technology) Library, and Golm Metabolome Database Library. The compound hits which showed maximum matching factor (MF) value (>600) and least deviation from the retention index (RI) was used for metabolite identity.

### Statistical analysis

Unless otherwise specified, a minimum of three biological replicates was used for every experiment. A student’s *t*-test was performed to determine significant differences using Sigma plot (version 11.0). Where required, the P values <0.05, <0.01, <0.001, are indicated by one, two, or three asterisks respectively. Heat maps were made using publicly available Morpheus software (https://software.broadinstitute.org/morpheus/).

## Author contributions

The whole study was conceived and designed by YS. HVK performed genetic and phenotypic segregation, ethylene evolution, fruit carotenoid, transcript profiling, proteome analyses, AD characterized the gene-edited lines, KS performed metabolite, volatile and hormone analyses, NRN designed the CRISPR constructs, off-target analysis, NG generated edited plants and performed initial screening for editing in tissue culture explants, NPT performed initial proteome analyses, Brix, firmness, pH and leaf carotenoid profiling, AJD performed sequencing and transcript profiling of selected genes, SS and KT performed fruit chloroplast assay and initial fruit ripening, KT made the first cross between AV and *Nps1*, SK performed leaf chlorophyll estimation and carbon dioxide evolution from fruits. Data were interpreted by HVK, KS, NRN, AD, RS, and YS. YS, and RS wrote the manuscript with inputs from HVK and NRN. All authors read and approved the manuscript.

## Acknowledgments

We thank Ms. Hemalatha for technical help with GeLCMS and Dr. Eros Kharshiing for critical reading of the manuscript. This work was supported by the Department of Biotechnology (DBT), India (BT/PR11671/PBD/16/828/2008, BT/PR/6983/PBD/16/1007/2012, and BT/COE/34/SP15209/2015) to YS and RS, University Grants Commission, India, research fellowship to HVK, Council of Scientific and Industrial Research, India research fellowship to KT, Department of Biotechnology research fellowship to AD, NPT and AJD.

---

**Supplementary Figure 1.**
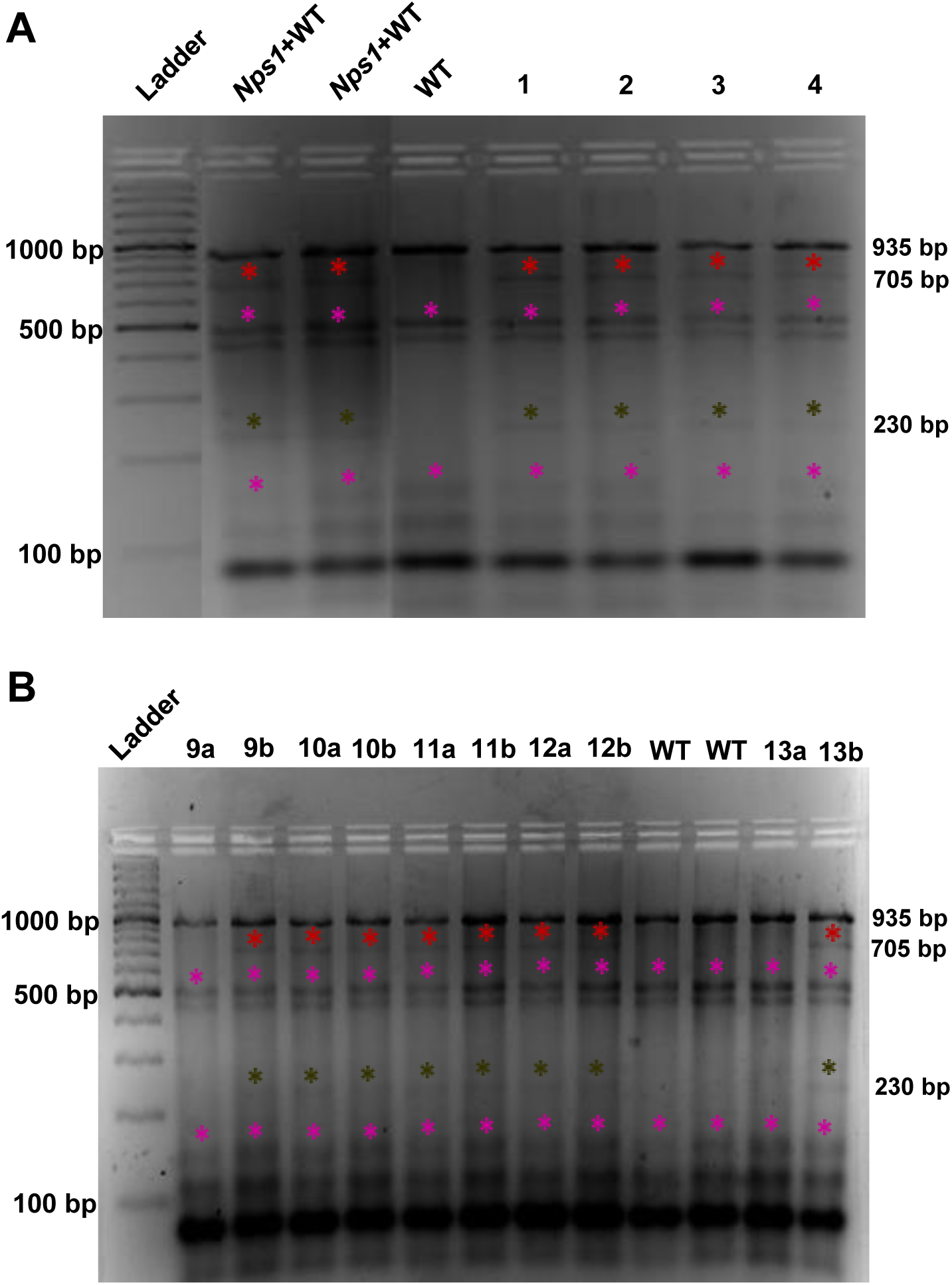
Representative figure showing CELI endonuclease assay used for detection of the *phot1* mutation and its zygosity in backcrossed plants. Genomic DNA from F_1_ and F_2_ plants was amplified using primers encompassing the *phot1* mutation site. The PCR product from the backcrossed plants was digested with CELI endonuclease either without mixing (homoduplex) or after mixing with WT (AC or AV) PCR product; heteroduplex) to ascertain the zygosity of plants. The digested products were separated on 1.2% (w/v) agarose gel and analyzed. The PCR product (935 bp) having *phot1* mutation in heterozygous condition upon CELI digestion generates two fragments of 705 bp and 230 bp size. Note that if digested fragments are observed in the homoduplex condition, it indicates that the *phot1* mutation is heterozygous. If the digested fragments are observed only in heteroduplex condition, then the plant is homozygous for the *phot1* mutation. *Nps1+*WT (heteroduplex) and WT (homoduplex) digested samples are shown as controls. **A**, Confirmation of introgression of *phot1* mutation in F_1_ through CELI digestion. Lanes 1 and 3 are loaded with homoduplex of F_1_ plants and 2, 4 are loaded with heteroduplex (F_1_*+*WT). **B**, CELI assay to identify and confirm segregation in F_2_ population obtained from WT♀x*Nps1♂* cross. Lanes 9a, 10a, 11a, 12a and 13 a are loaded with homoduplex of F_2_ plants; and lanes 9b, 10b, 11b, 12b and 13b are loaded with heteroduplex (F_2_*+*WT). Note that F_2_ lines- 10, 11 and 12 are heterozygous for *phot1* mutation and the lines 9 and 13 are homozygous for *phot1* mutation. Red* -705 bp product, black* -230 bp product, pink* - non specific products.

**Supplemental Figure 2.**
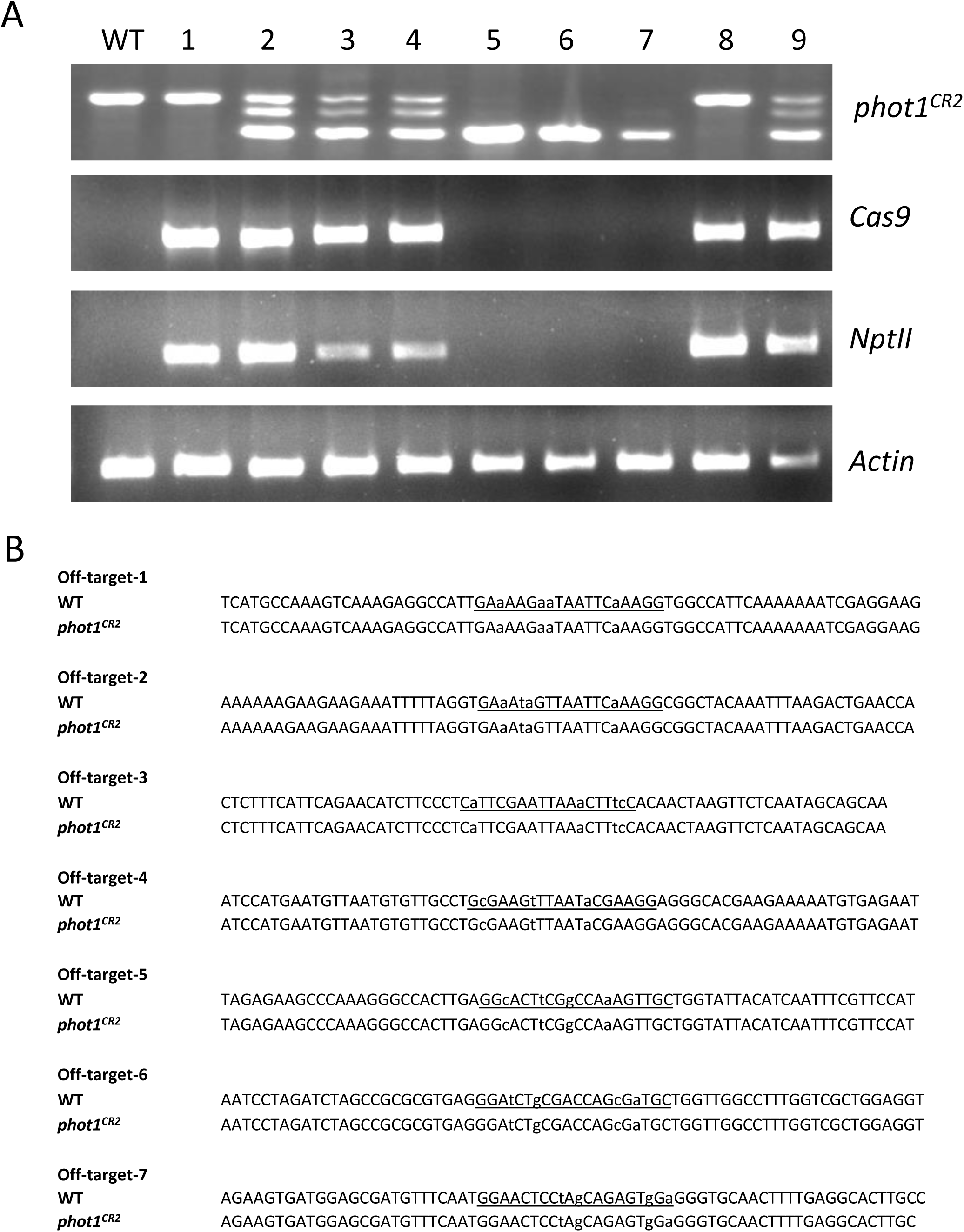
A representative image showing the identification of homozygous and CAS9-free plants in T_1_ and T_2_ generations. **A,** Identification of homozygous *phot1^CR2^* mutant in T_1_ followed by screening of Cas9- free plants in T_2_ generation. Examination of *phot1^CR2^*, *Cas9*, *nptII* genes in the T_1_ and T_2_ plants by PCR. Plant number 5, 6 and 7 are putative homozygotes and free of T-DNA insertion. **B,** Mutation patterns were examined in 7 most potential off-target sites for sgRNA1 (1-4) and sgRNA2 (5-7). Target sequence of potential off-target sites was underlined, with lowercase letters representing mismatches. Note that no off-target editing events were detected.

**Supplemental Figure 3.**
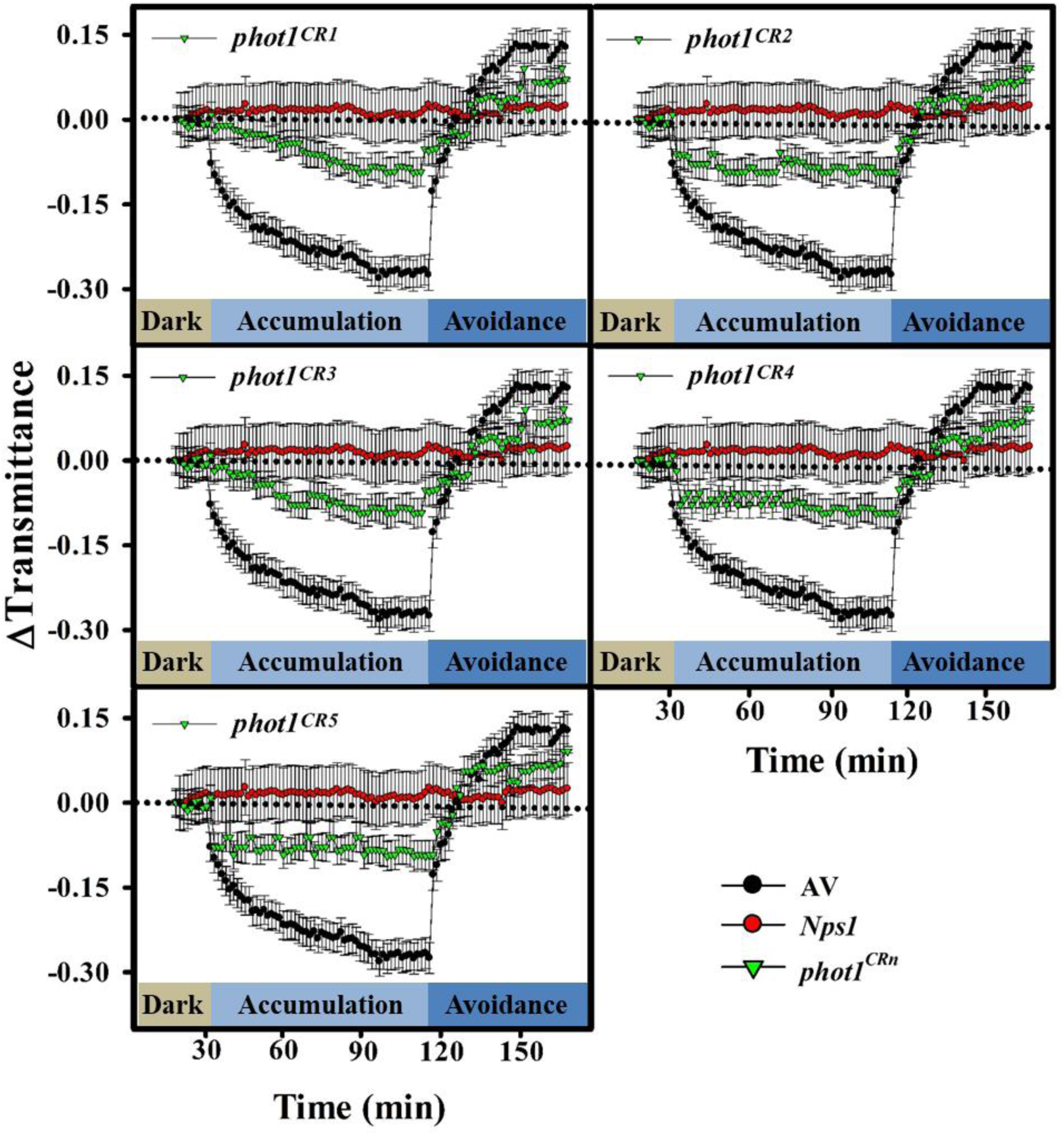
Chloroplast re-location response in the leaf discs of AV, *Nps1* and *phot1^CR^* alleles. The leaf discs were dark-adapted for 12 h and exposed to low fluence blue light (3.2 µmol/m^2^/s) to induce chloroplast accumulation response. Thereafter, the leaf discs were exposed to high fluence blue light (80 µmol/m^2^/s) to induce chloroplast avoidance response. Note that compared to total loss of chloroplast relocation response in *Nps1*, *phot1^CR^* alleles display reduced chloroplast accumulation and avoidance responses. Data are means ± SE (n=8).

**Supplemental Figure 4.**
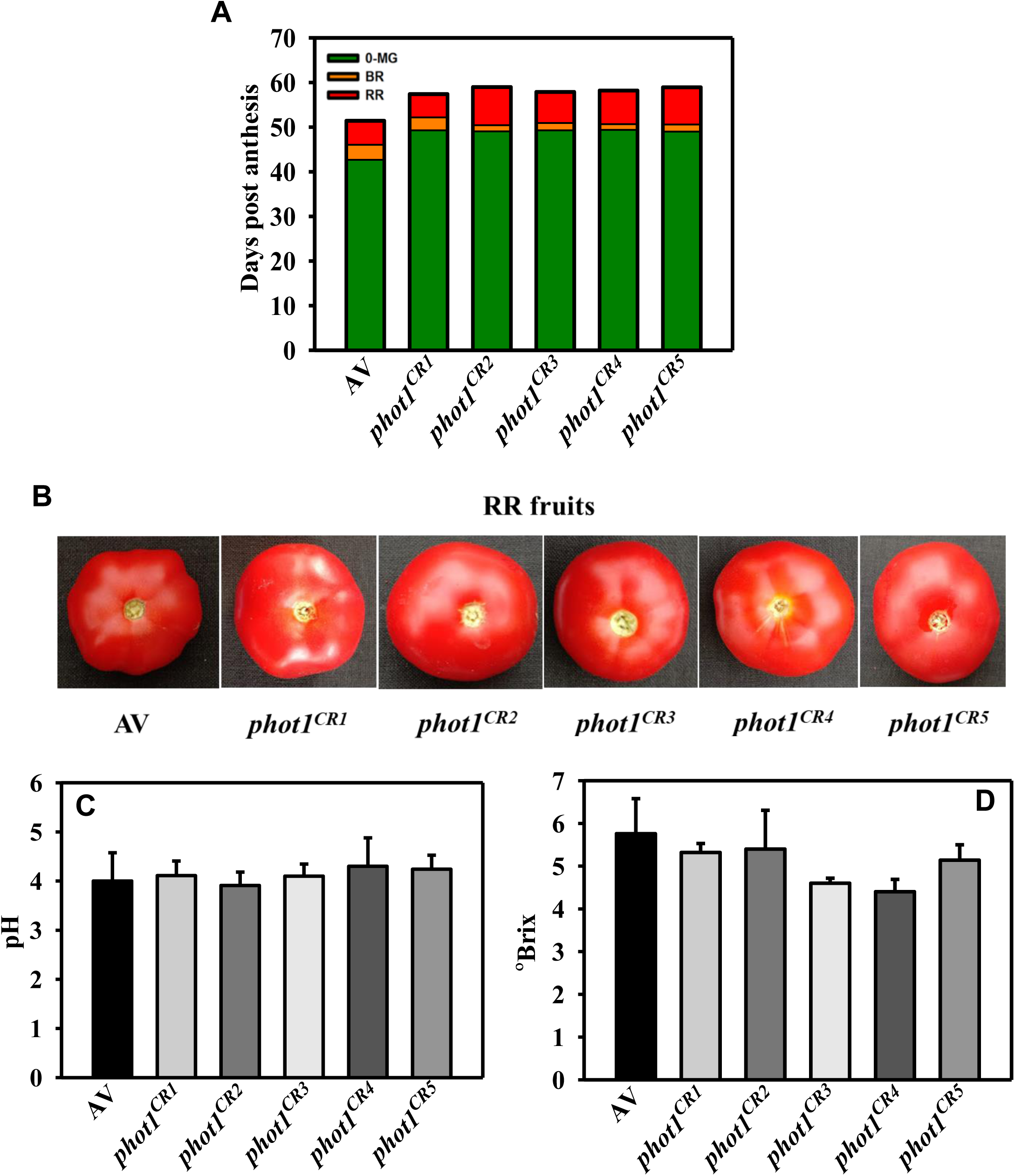
Chronological development of fruits during ripening in AV and *phot1^CR^* lines, pH, °Brix and colour of their ripe fruits. **A**, Chronological development of fruits through different ripening stages. Note that the onset of ripening is delayed and the transition to subsequent ripening stages is varied in *phot1^CR^* lines than AV. **B**, Red ripe fruits of *phot1^CR^* lines. Note the lighter red colour in *phot1^CR^* lines than the wild type progenitor, AV. **C-D**, pH and °Brix of ripe fruits. Data are means ± SE (n=15 for fruit development and n=5 for others), *P ≤ 0.05.

**Supplemental Figure 5.**
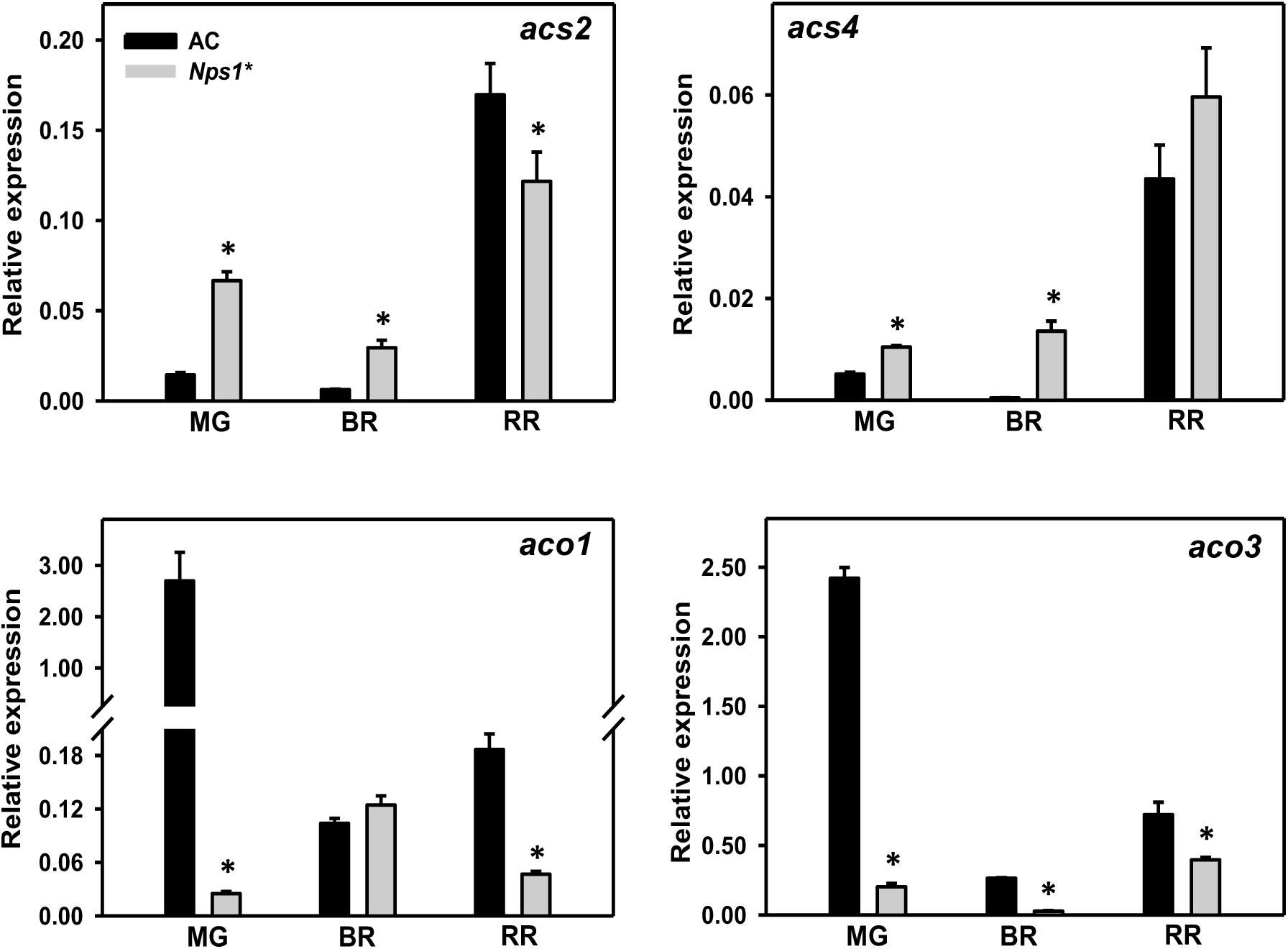
Relative transcript levels of *acs* and *aco* genes mediating system II ethylene biosynthesis in ripening fruits of AC and *Nps1*\*. The graphs depict data obtained after normalization with *β-actin* and *ubiquitin*. Data are means ± SE (n=3), *P ≤ 0.05.

**Supplemental Figure 6.**
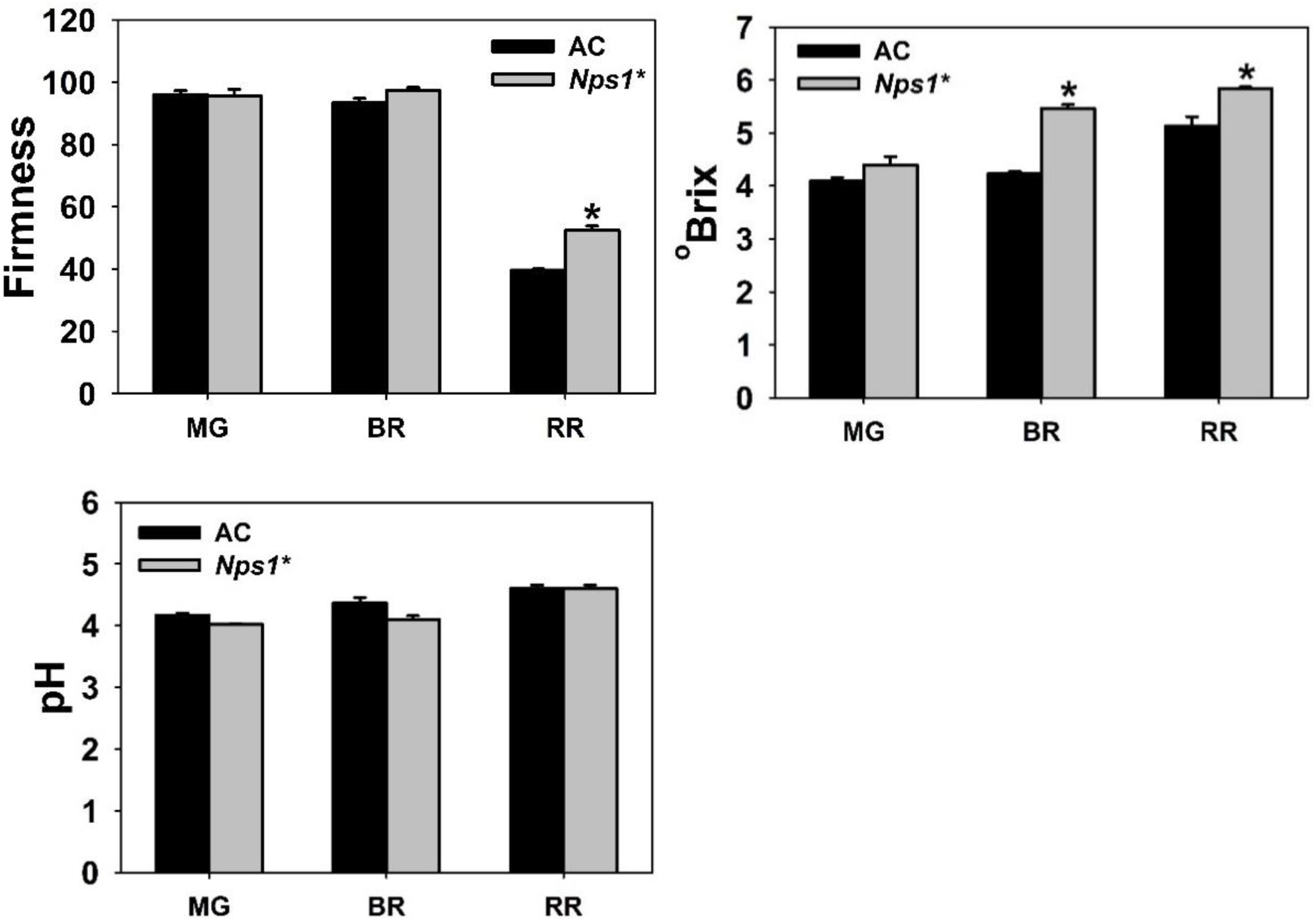
Fruit firmness, °Brix and pH of ripening fruits of AC and *Nps1*.* Data are means ± SE (n=3), *P ≤ 0.05.

**Supplemental Figure 7.**
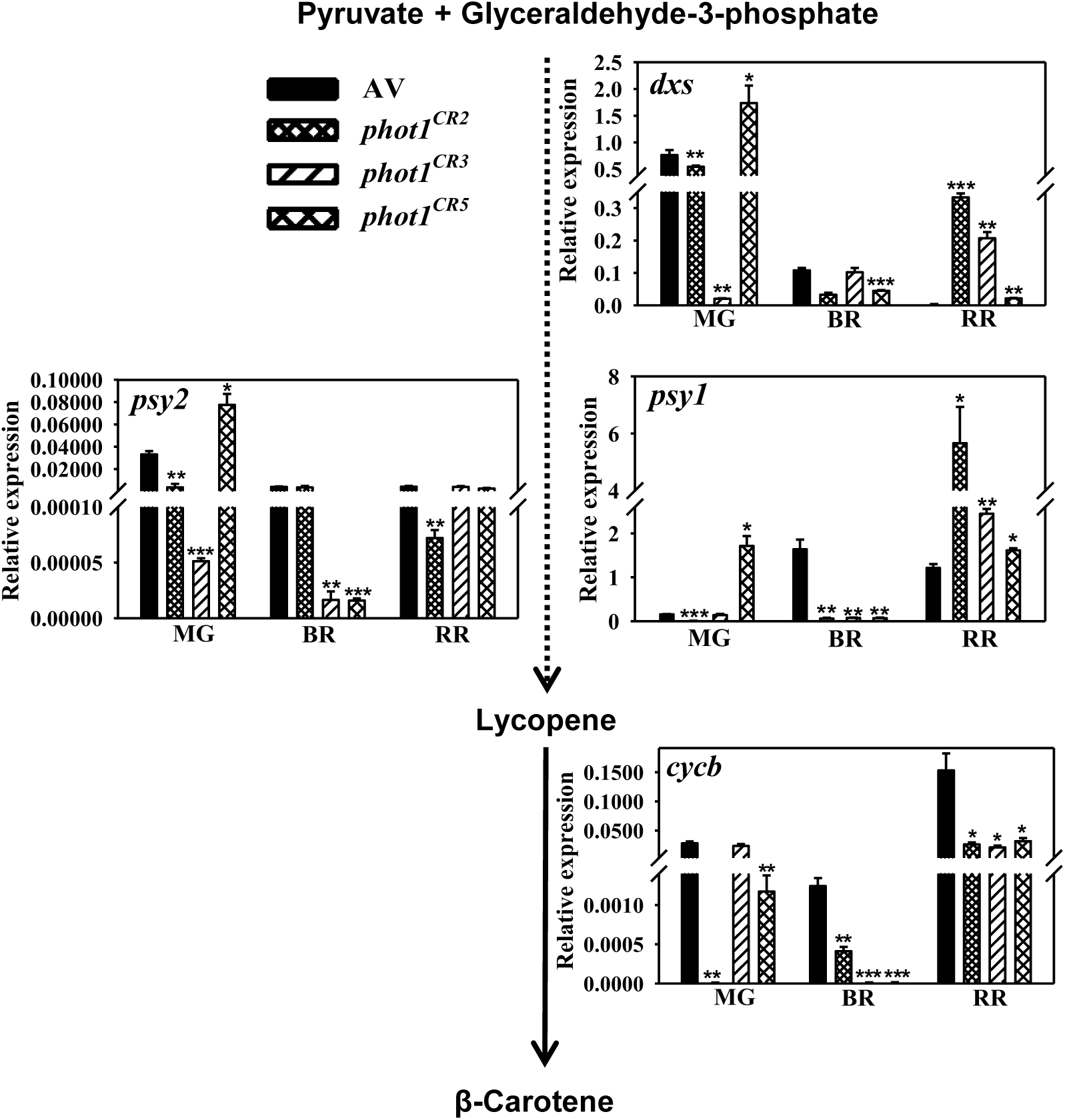
Relative expression of rate limiting genes mediating carotenoid biosynthesis in ripening fruits of AV and *phot^CR^* lines. The *phot^CR2^, phot^CR3^* and *phot^CR5^* lines are homozygous for editing and Cas9-free. Dotted arrows indicate multiple steps in the carotenoid biosynthesis pathway. The graphs indicate relative expression of genes obtained after normalization with *β-actin* and *ubiquitin*. Data are means ± SE (n=3), *P ≤ 0.05, **P ≤ 0.01 and ***P ≤ 0.001.

**Supplemental Figure 8.**
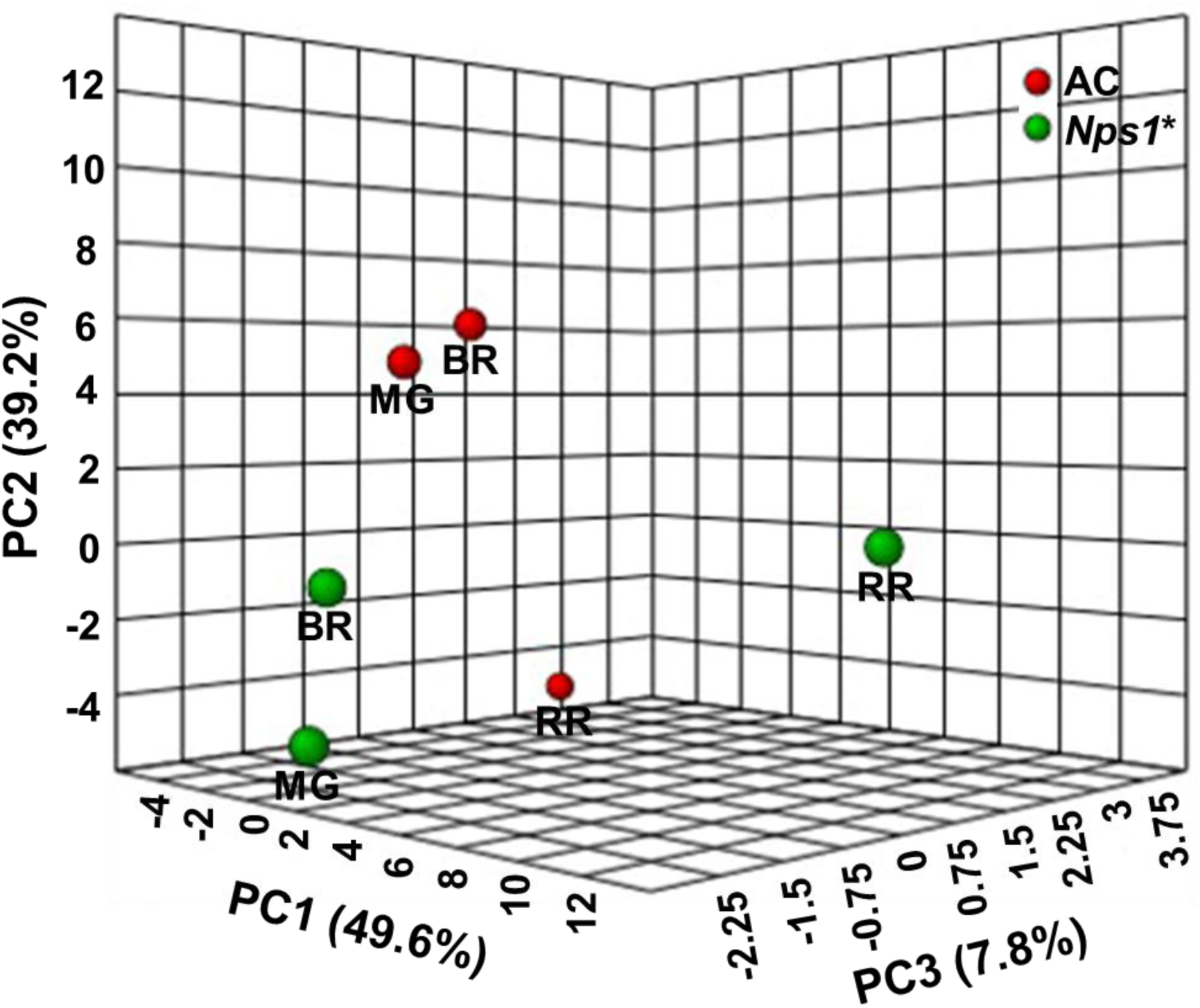
The principal component analysis of primary metabolites detected in ripening fruits of AC and *Nps1\**. Data are means ± SE (n=5), *P≤0.05.

**Supplemental Figure 9.**
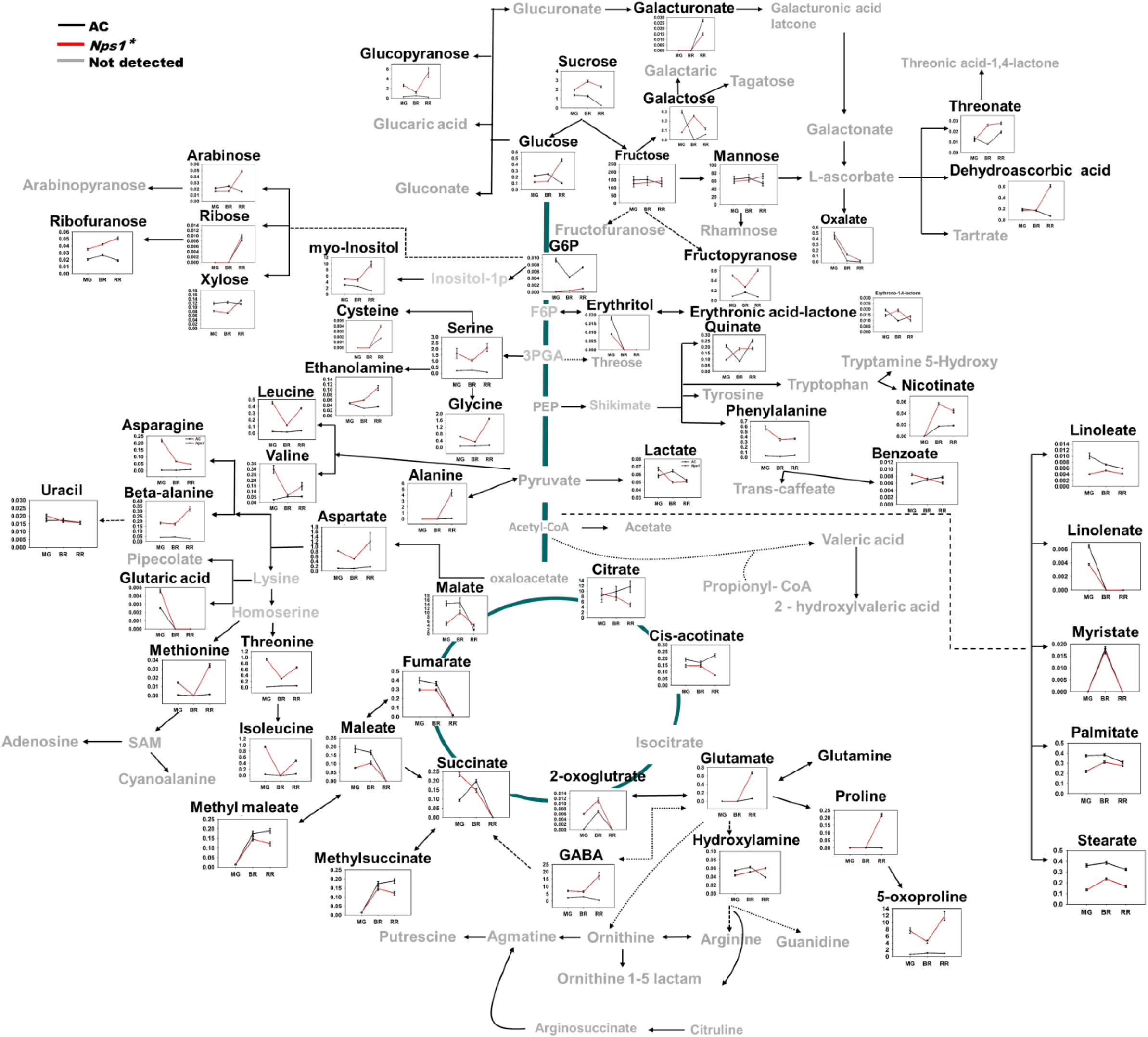
The metabolic shifts in AC and *Nps1\** fruits during ripening. An overview of the metabolic pathways representing relative abundance of metabolites at different ripening stages in fruits of *Nps1\** and AC. The Y-axis represents the relative levels of metabolites with reference to internal standard ribitol. X axis denotes metabolite levels at MG-mature green, BR-breaker, and RR- red-ripe stages.

**Supplemental Figure 10.**
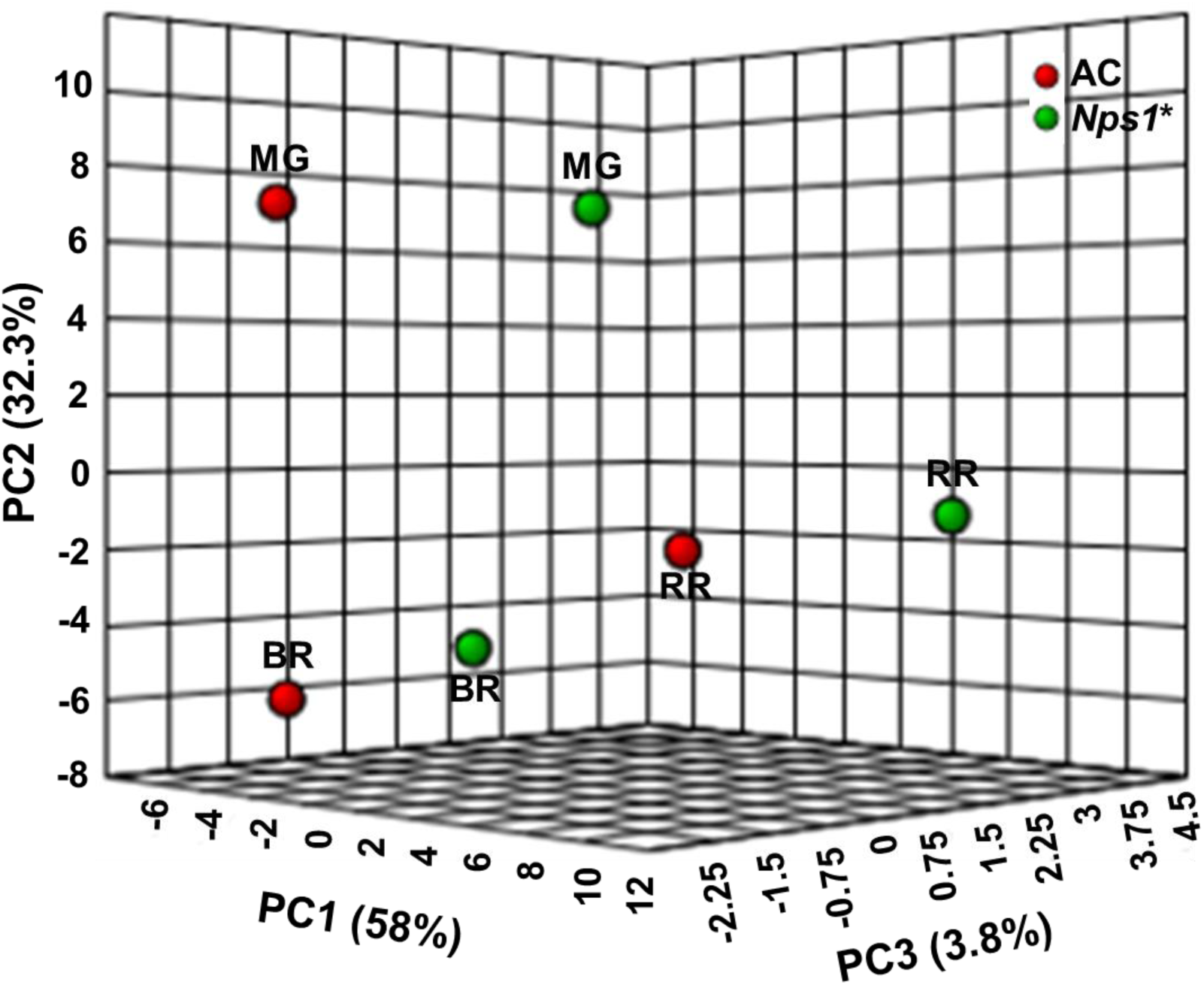
The principal component analysis of volatiles detected in ripening fruits of AC and *Nps1\**. Data are means ± SE (n=8), *P≤0.05.

**Supplemental Figure 11.**
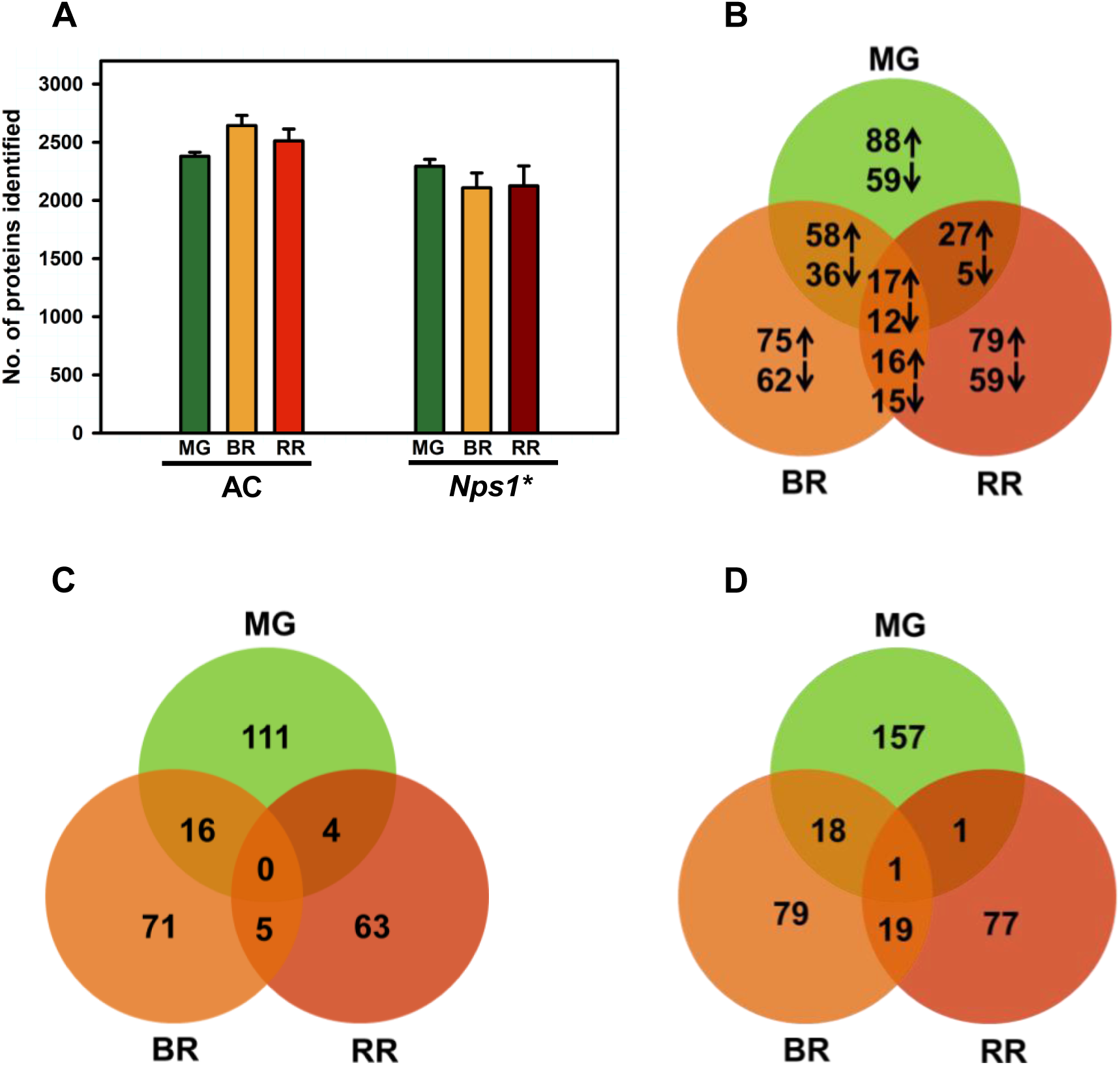
Proteome profiling in ripening fruits of AC and *Nps1*.* Data are means ± SE (n=3), P≤0.05; up-regulation (↑), down-regulation (↓). **A**, Number of proteins identified in AC and *Nps1\** fruits. **B**, Differentially expressed proteins in *Nps1\** in comparison with AC. **C**, Proteins identified solely in *Nps1\**. **D**, Proteins identified only in AC.

**Supplemental Figure 12.**
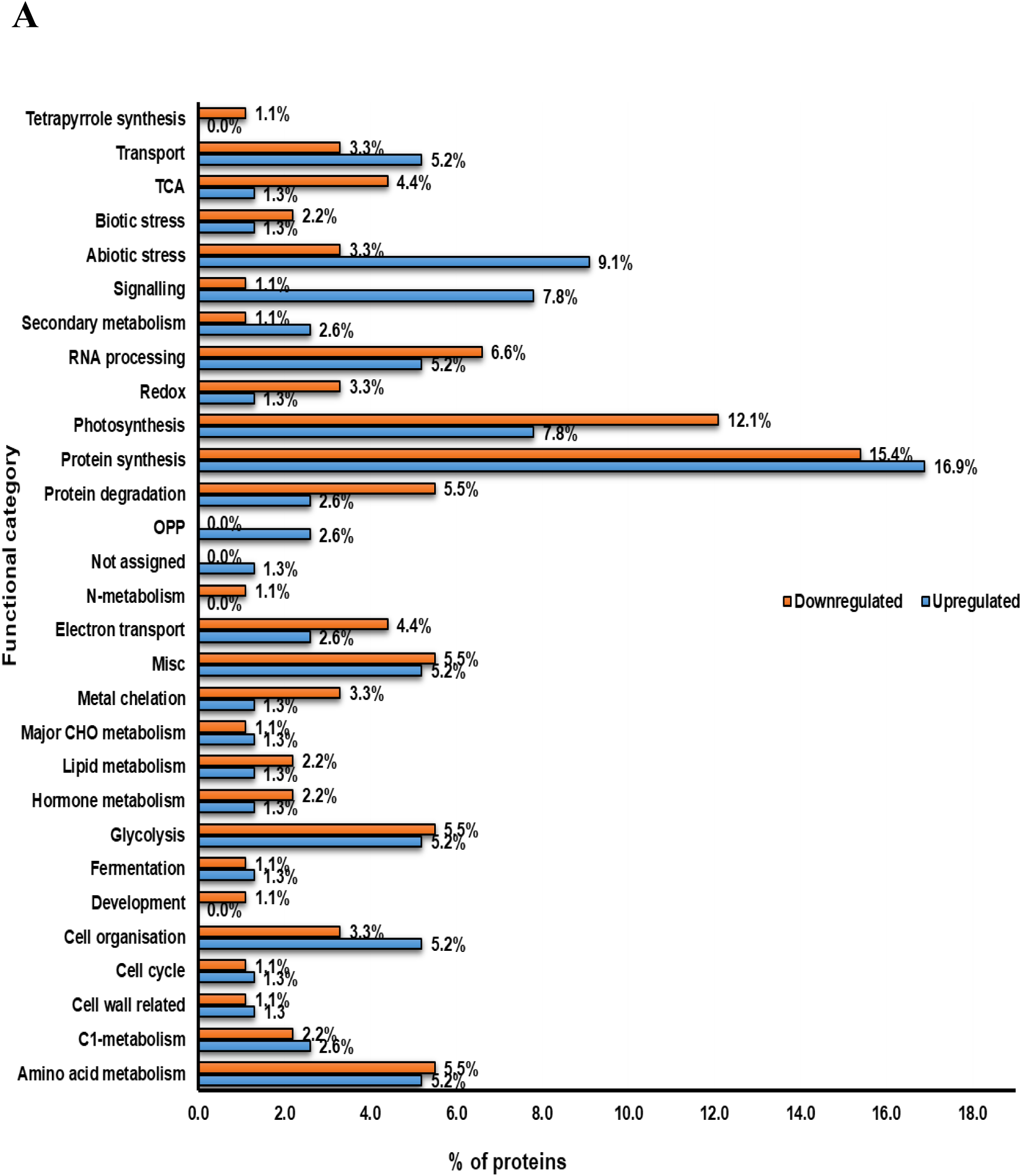
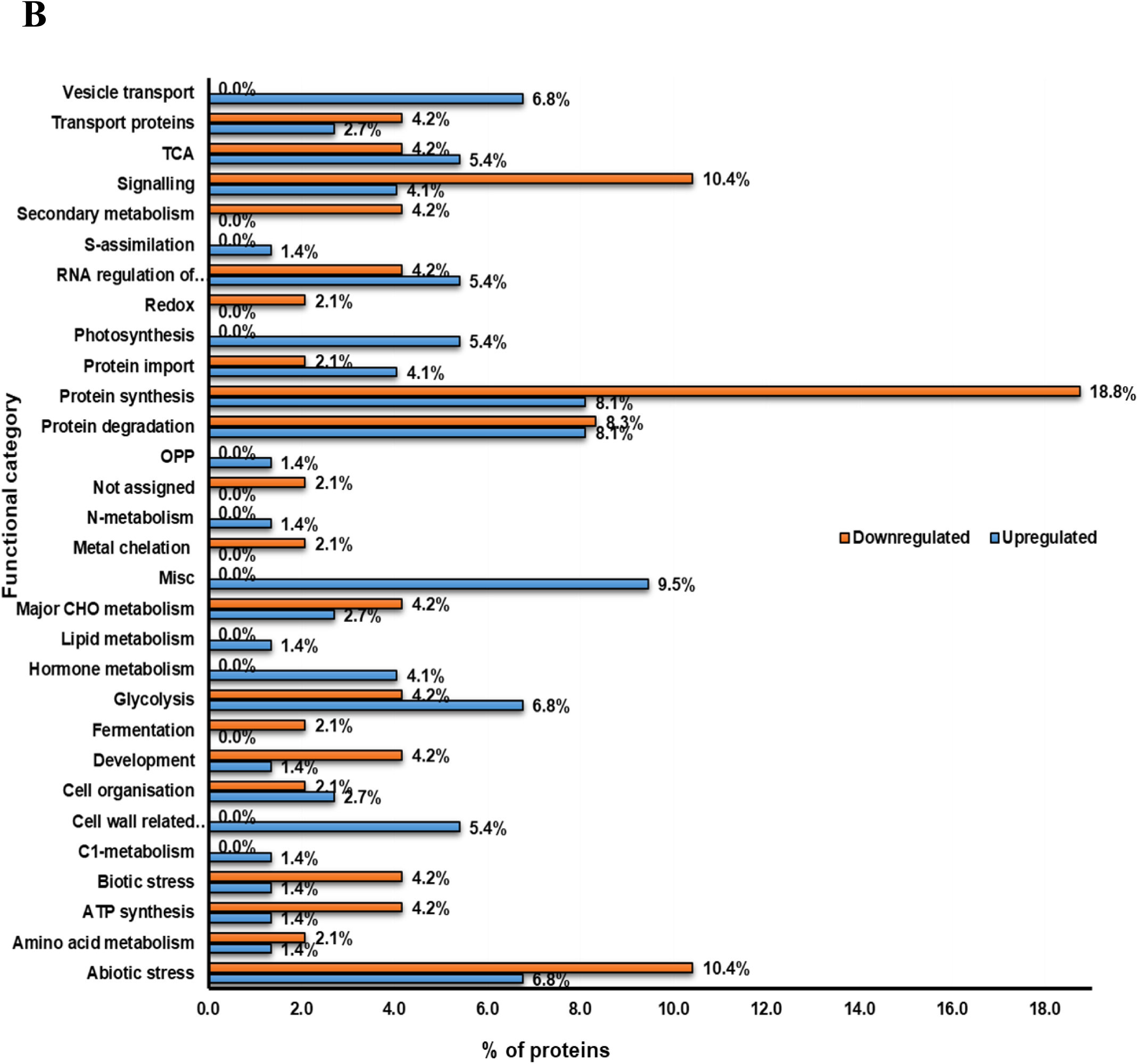
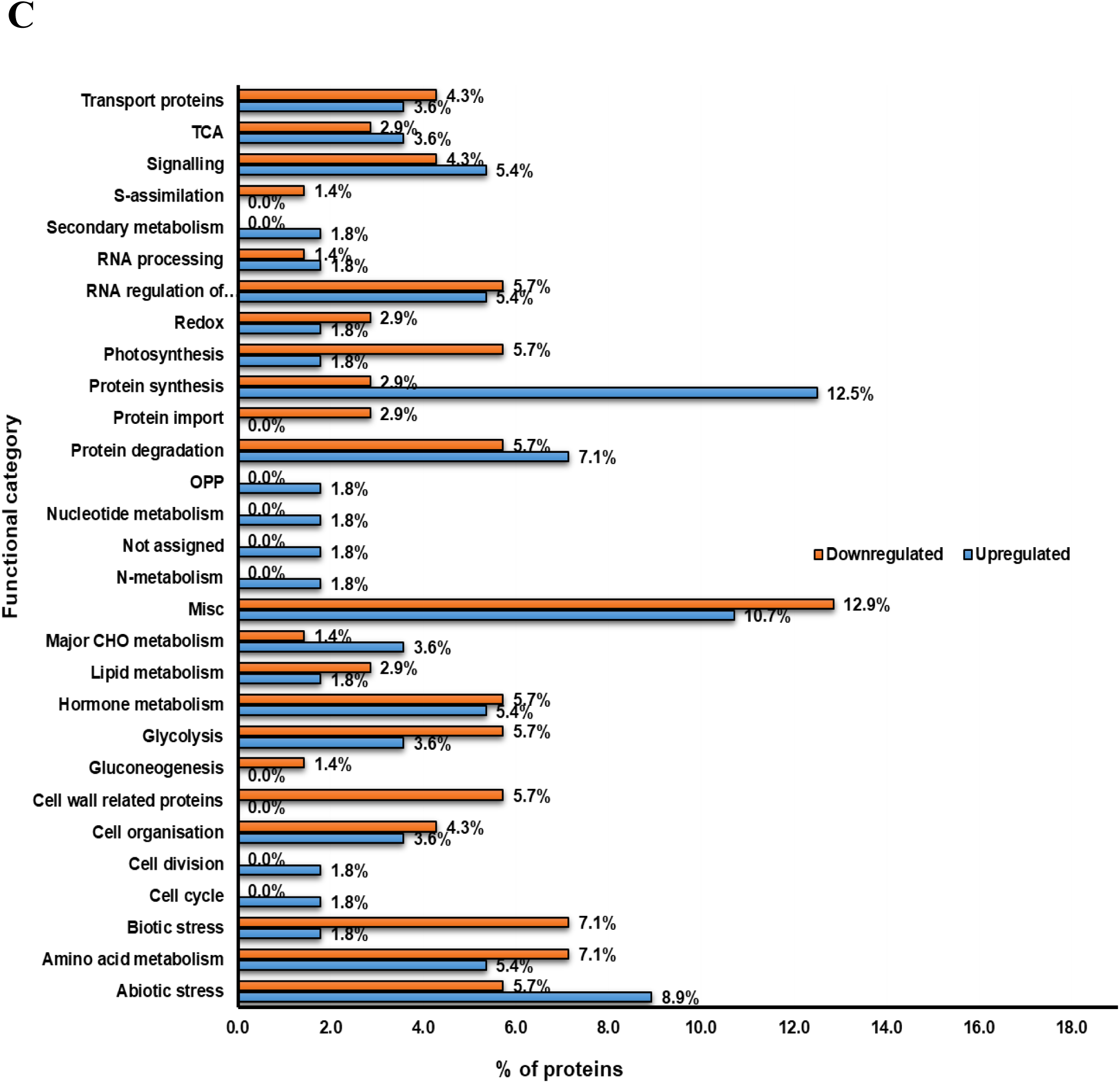
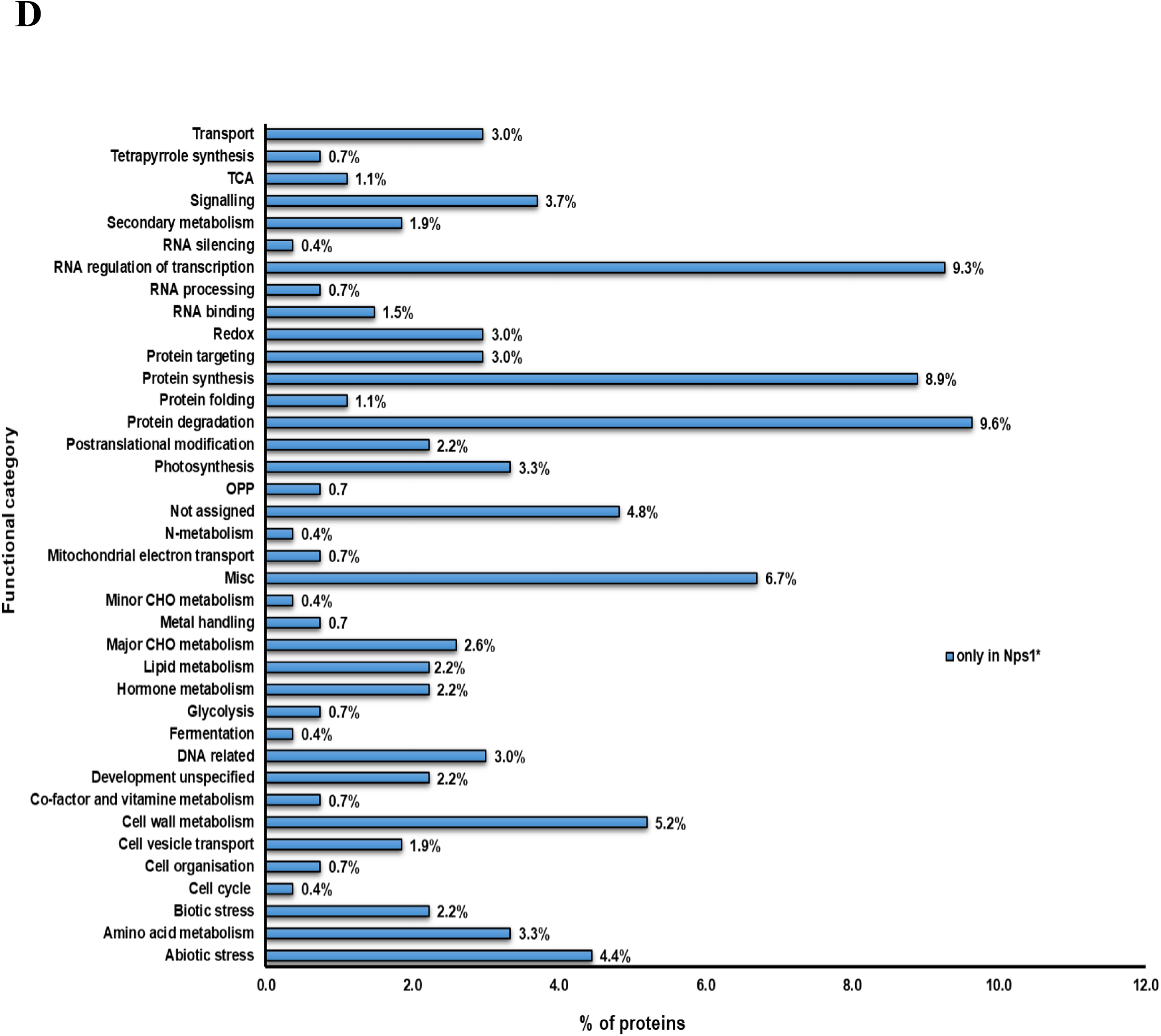
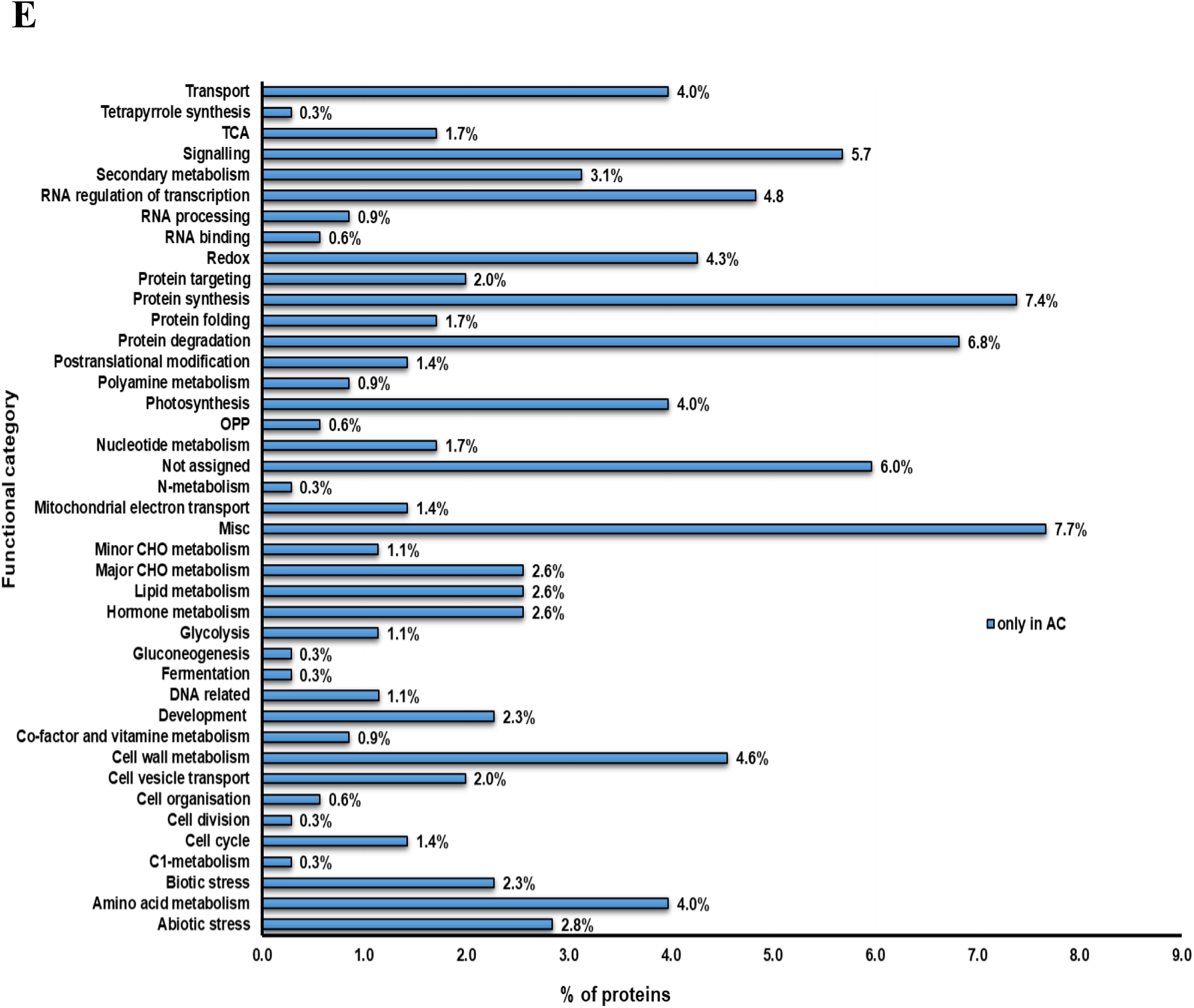
Functional classification of differentially expressed proteins in the fruits of *Nps1*\* and AC. The differentially expressed proteins in the fruits of *Nps1*\* compared to AC were functionally classified using MapMan. These proteins are also listed in Supplemental Table 5 and 6. **A,** MG stage; **B,** BR stage; **C,** RR stage; **D.** Proteins present only in *Nps1*\*; **E,** Proteins present only in AC.

**Supplemental Figure 13.**
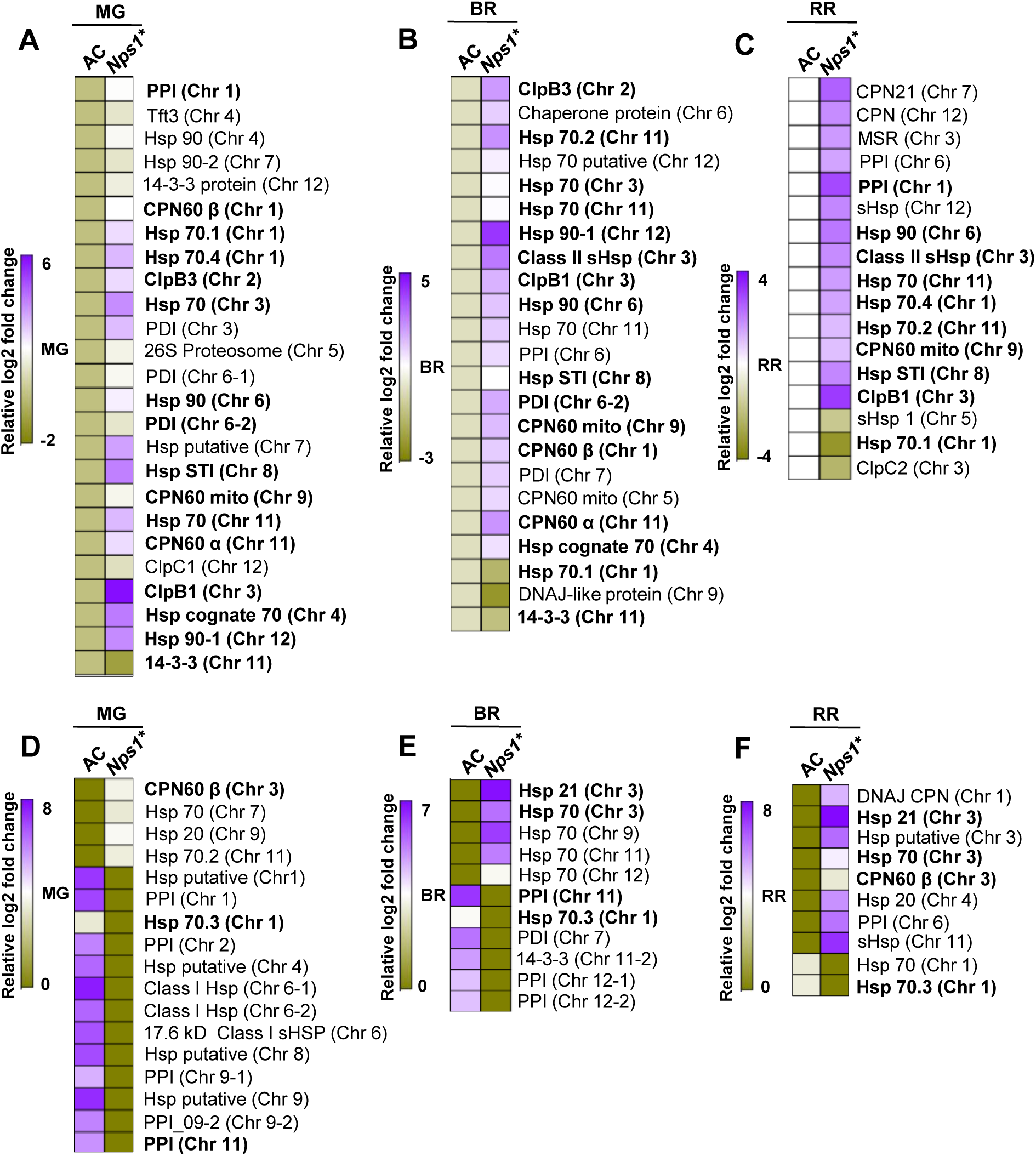
Relative abundances of heat shock proteins and proteins involved in protein folding and degradation in AC and *Nps1\** fruits at different stages of ripening. Heat maps were generated using normalized log2 fold changes for differentially expressed proteins. For proteins detected only in *Nps1\** or in AC, the missing values were replaced with 1/10 of the lowest value in the dataset and the heat maps were generated with normalized log2 abundances. **A-C**, Differentially expressed proteins in *Nps1\** in comparison with AC. **D-F**, Proteins detected only in AC or *Nps1\**. Protein names in bold represents proteins identified in two or all three stages. The chromosome number for each protein is given in parenthesis. The abbreviations are Hsp-Heat shock protein; PPI-Peptidyl-prolyl cis-trans isomerase; CPN60-Chaperonin60; PDI-Protein disulfide isomerase; 26S P-26S proteasome non-ATPase regulatory subunit-like protein; Hsp STI-Heat shock protein Stress-Induced; sHSP-small heat shock protein; MSR-methionine sulfoxide reductase.

**Supplemental Figure 14.**
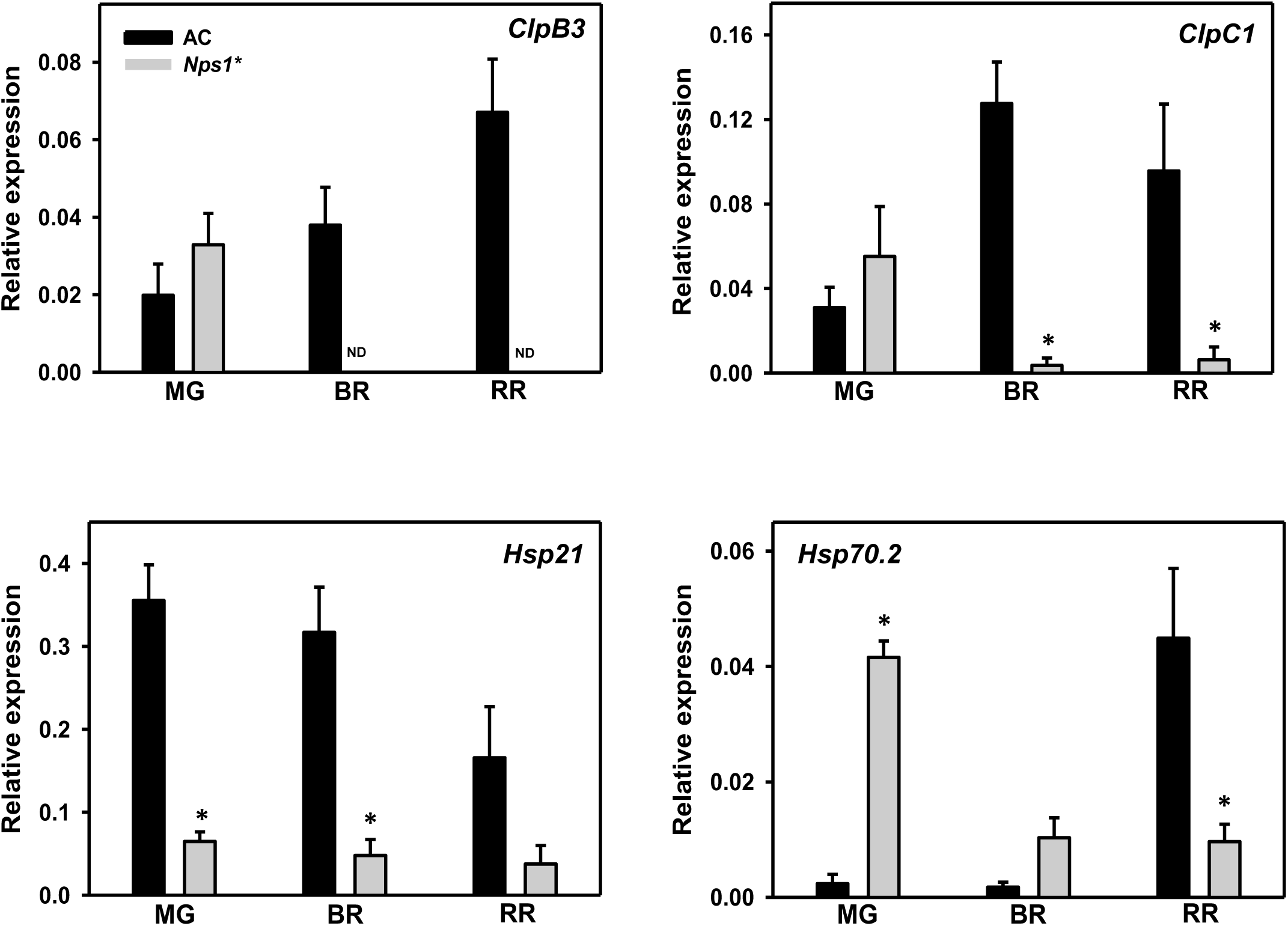
Relative expression of selected *Clp* proteases and *Hsps* in ripening fruits of AC and *Nps1*\*. The graphs depict data obtained after normalization with *β-actin* and *ubiquitin*. Data are means ± SE (n=3), *P ≤ 0.05; ND-not detected.

**Supplemental Figure 15.**
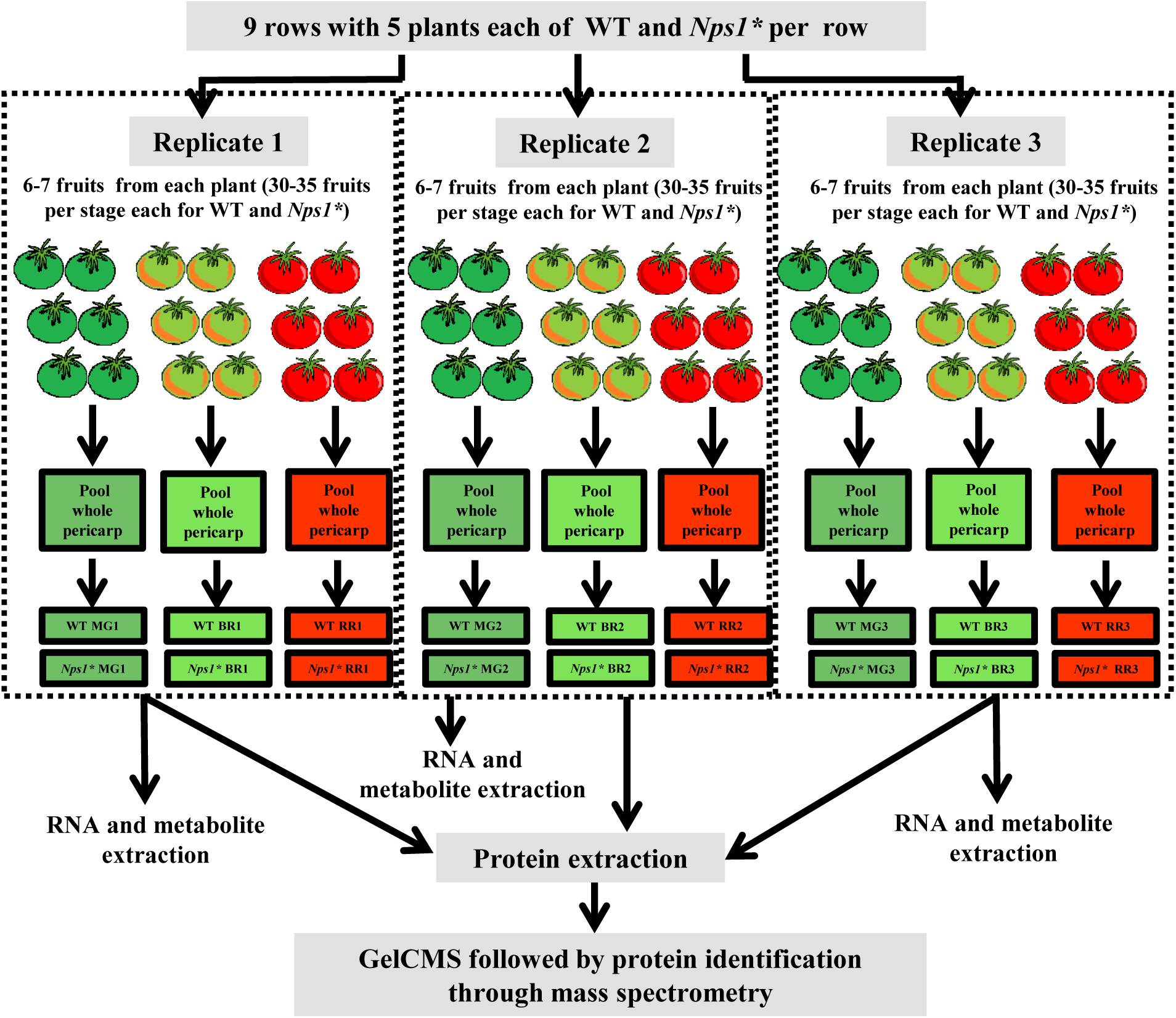
Experimental design describing growth of plant material, collection of fruit tissue at different stages of ripening in AC and *Nps1*\* for carotenoid, metabolite, RNA and protein extraction.

**Supplemental Table 1.**
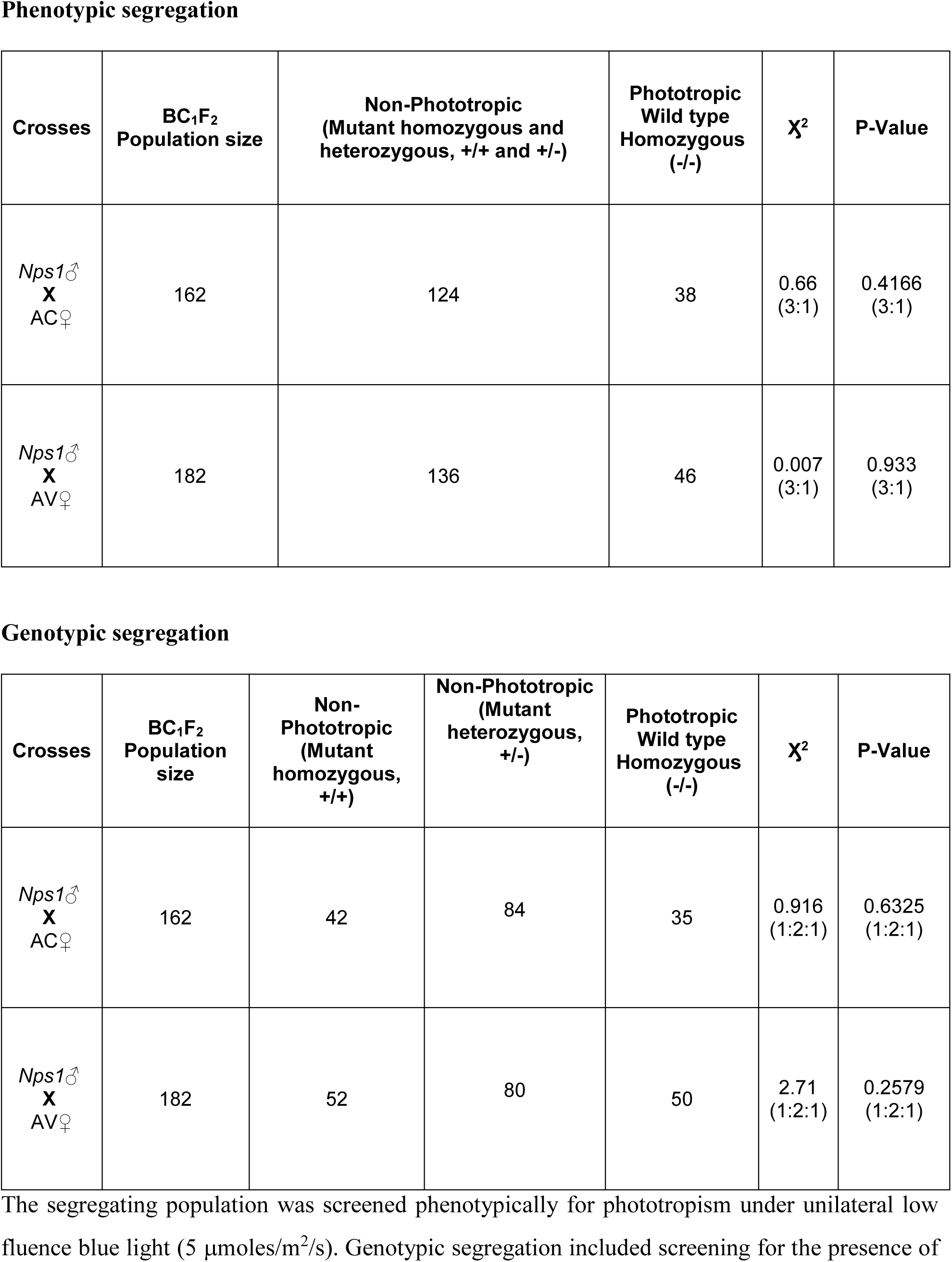
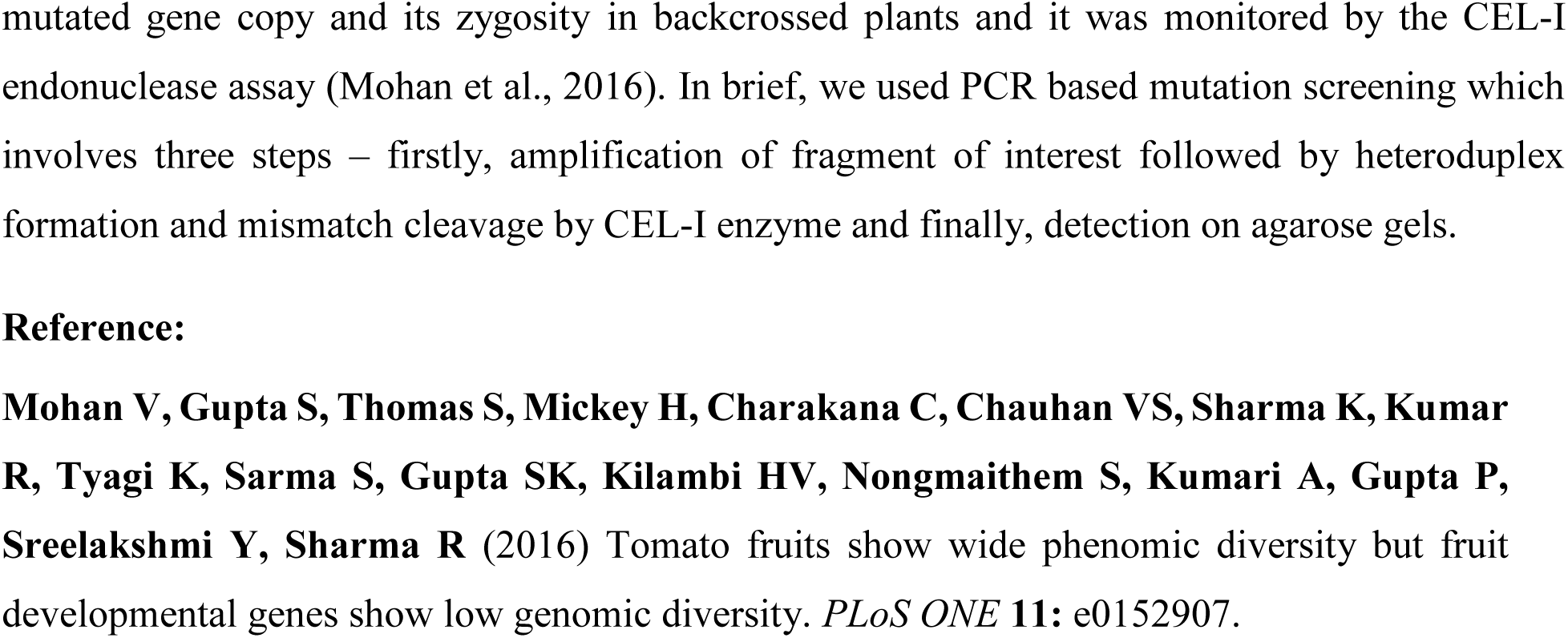
The genetic segregation of *Nps1* in BC_1_F_2_ generation in AC and AV backgrounds.

**Supplemental Table 2.** Carotenoid and chlorophyll contents in the leaves of 70 day-old plants of AC, *Nps1* and *Nps1*.* Carotenoid and chlorophyll contents are presented as μg/g fresh weight. Data are means (n=3) ± SE. Chl, chlorophyll; Car, carotenoid. Student’s t-test was used to determine the significant differences in the metabolites between AC and the mutant. Values in bold indicate significant differences: *P<0.05.

| Metabolite ($\mu\text{g gFW}^{-1}$) | AC | *Nps1* | *Nps1\** |
| --- | --- | --- | --- |
| **β-Carotene** | 37.46 ± 2.04 | 43.26 ± 1.74 | 45.45 ± 3.96 |
| **Lutein** | 70.95 ± 4.81 | 76.93 ± 3.32 | 79.23 ± 6.74 |
| **Violaxanthin** | 5.40 ± 1.3 | 5.91 ± 0.94 | 7.68 ± 1.99 |
| **Neoxanthin** | 24.94 ± 3.34 | 26.03 ± 1.09 | 30.90 ± 4.94 |
| **Total Car** | 138.77 ± 10.64 | 152.14 ± 4.80 | 163.26 ± 17.32 |
| **Chlorophyll a** | 455.64 ± 4.04 | 468.24 ± 17.56 | 450.97 ± 1.77 |
| **Chlorophyll b** | 520.88 ± 2.01 | **454.76 ± 0.66\*** | 454.18 ± 10.77 |
| **Chl a: Chl b** | 0.9763 ± 0.043 | 0.8658 ± 0.005 | 1.0024 ± 0.0234 |
| **Total Chl** | 923.87 ± 14.74 | **971.85 ± 2.38\*** | 908 ± 10.87 |
| **CHL:CAR** | 6.09 ± 0.28 | 7.09 ± 0.52 | 5.69 ± 0.60 |

**Supplemental Table 8.**
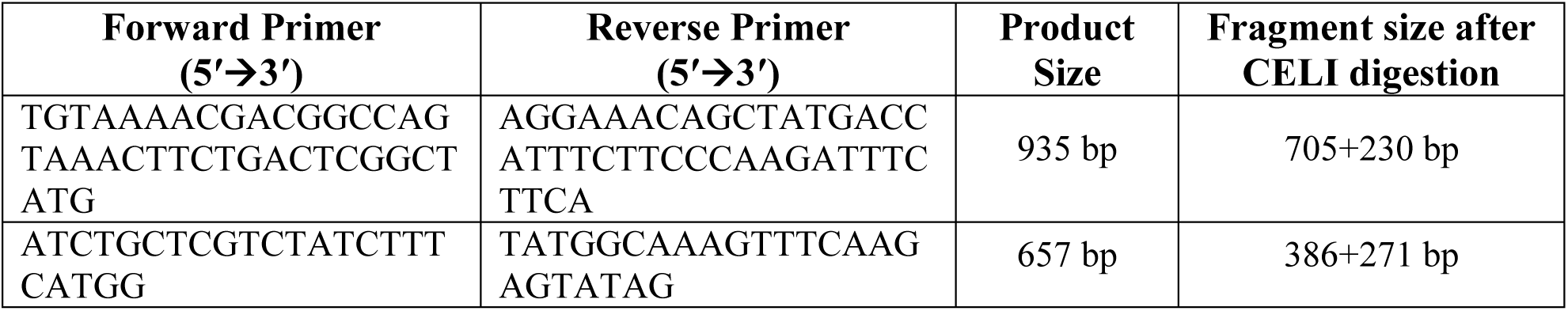
Primers used for detection of mutation, zygosity and sequencing of PCR products in *phot1* gene in segregating population.

**Supplemental Table 9:**
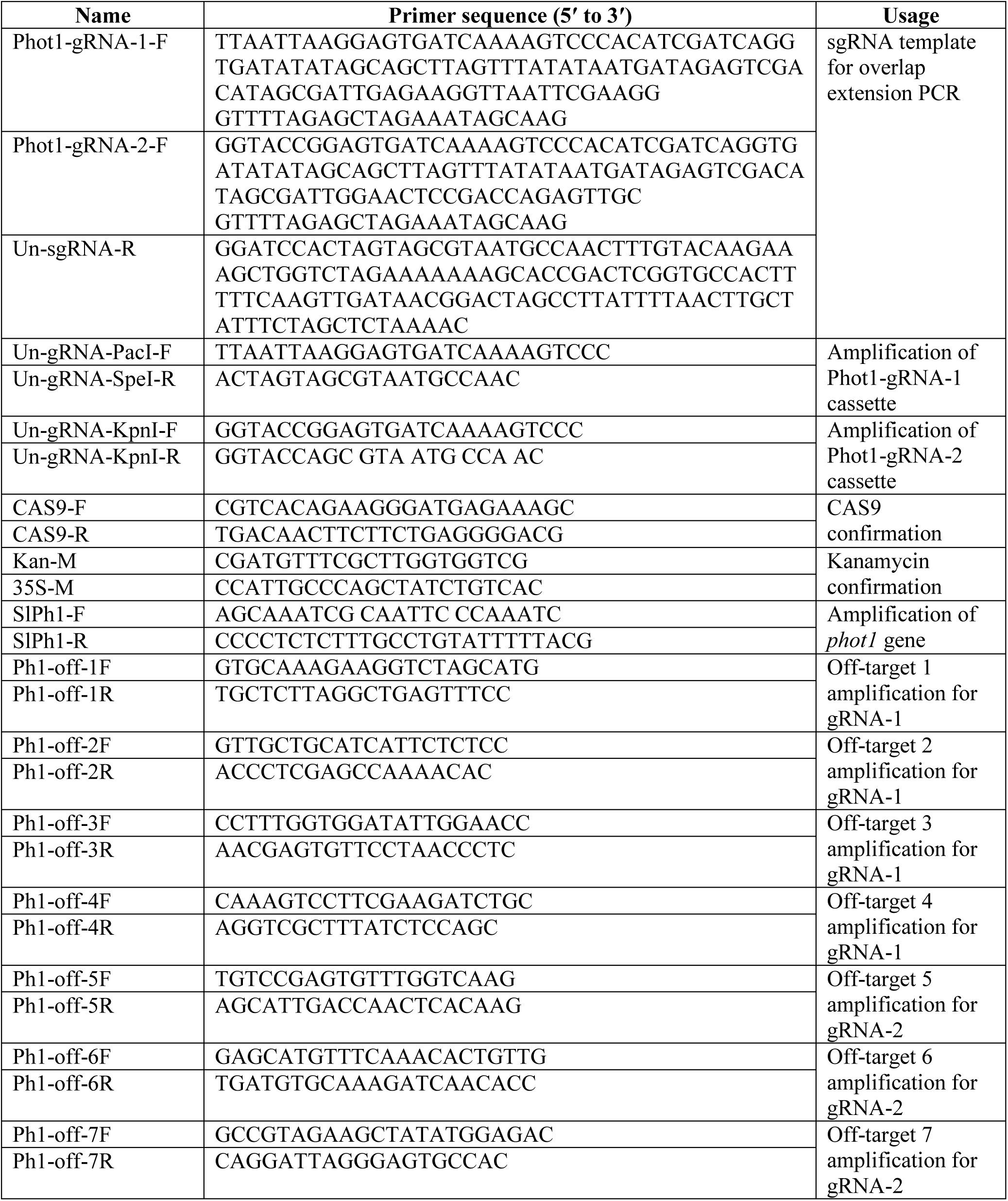
List of primers used for making CRISPR constructs for *phot1* gene, for sequencing the editing events and the primers used for detecting off-target effects in the current study.

**Supplemental Table 10:**
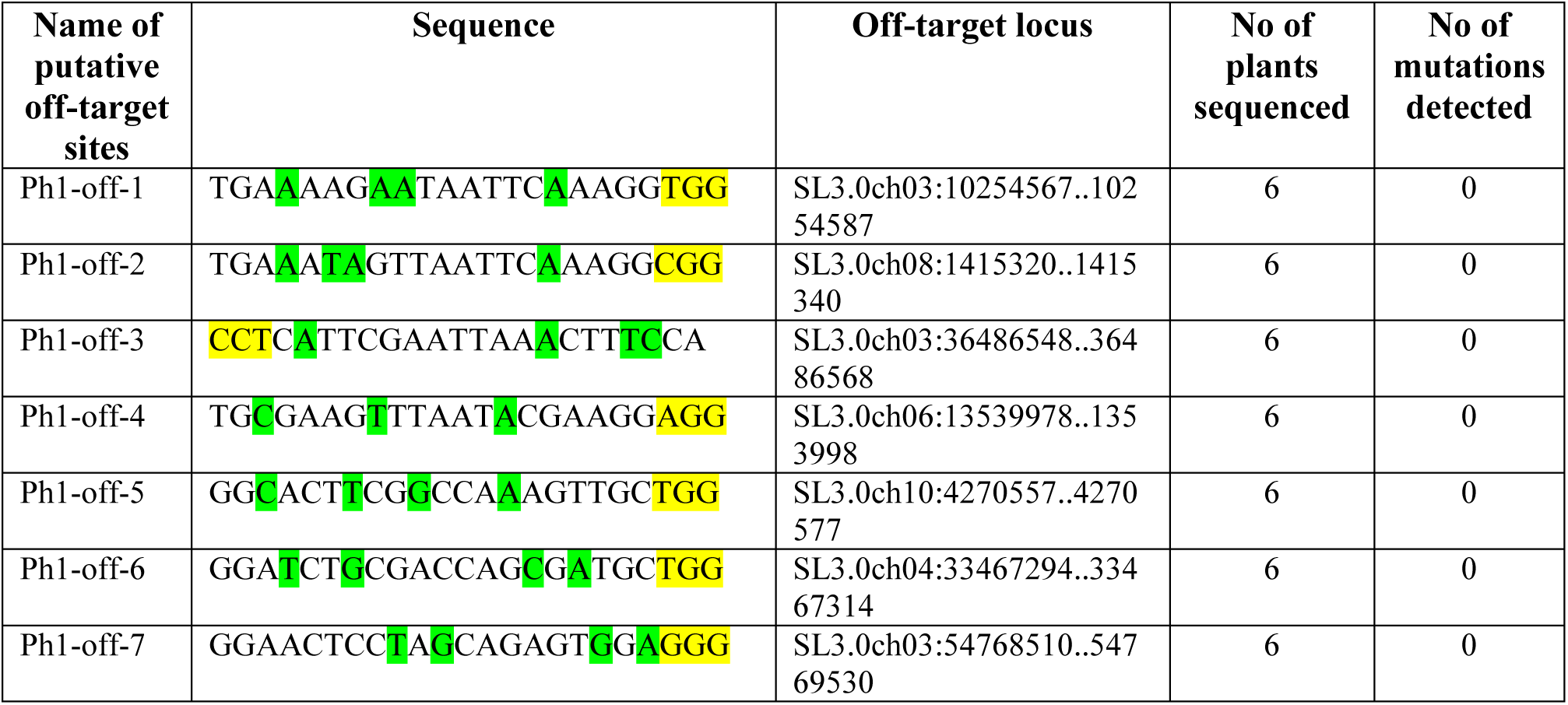
Identification of potential off-target sites and mutation detection. Off-target sites for *phot1gRNA1* and *phot1gRNA2* were predicted using CRISPR-P and CRISPOR web tools (http://cbi.hzau.edu.cn/crispr; http://crispor.tefor.net). Mismatch nucleotides are marked in green and PAM sequence (NGG) is highlighted in yellow. The *phot1^CR2^*, *phot1^CR3^* and *phot1^CR5^* lines that are homozygous and Cas9-free in T_2_ generation were used for off-target analysis. The genomic DNA surrounding the potential off-target locus was amplified using specific primers and the PCR products were analysed by Sanger sequencing.

**Supplemental Table 11.**
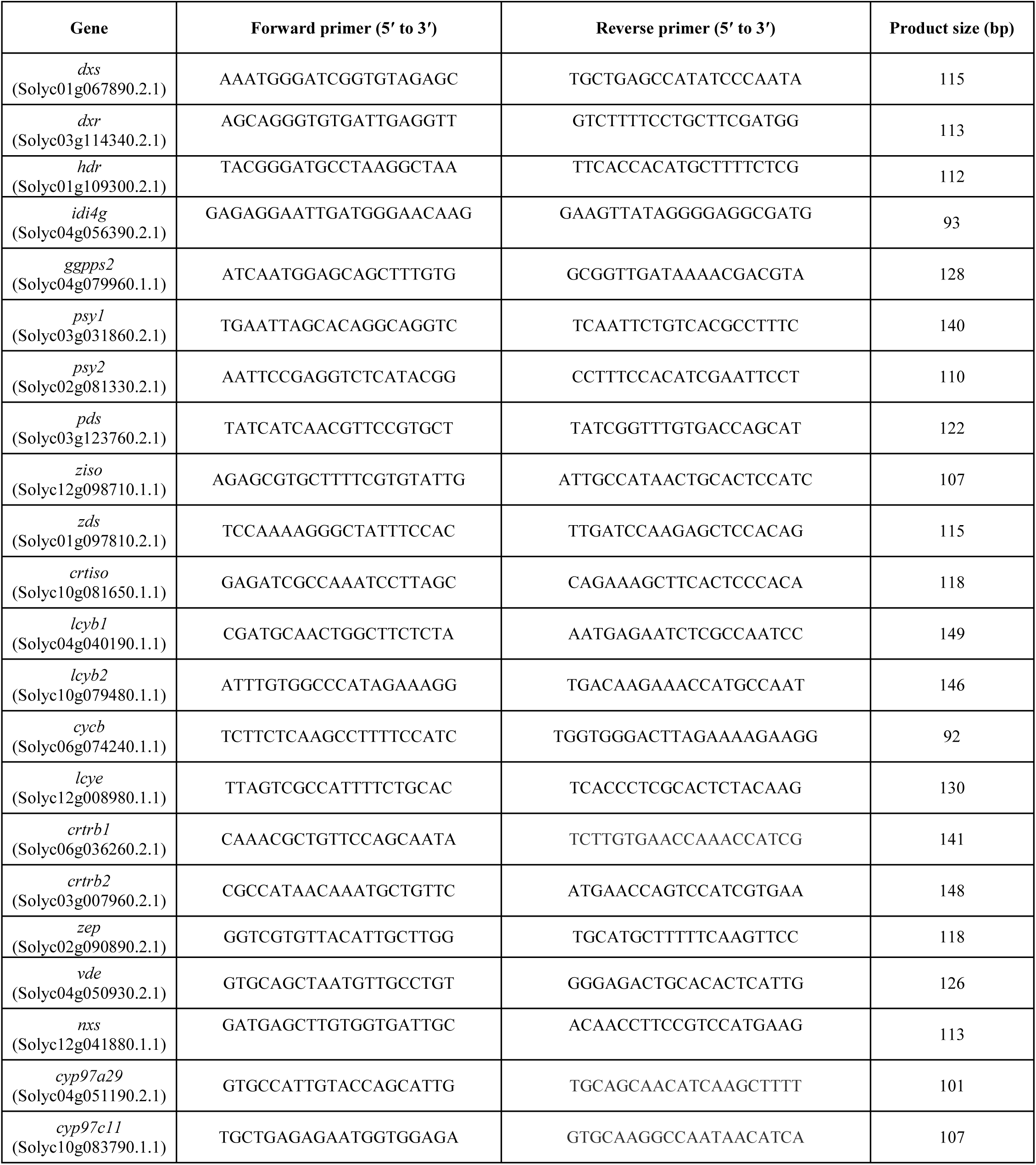
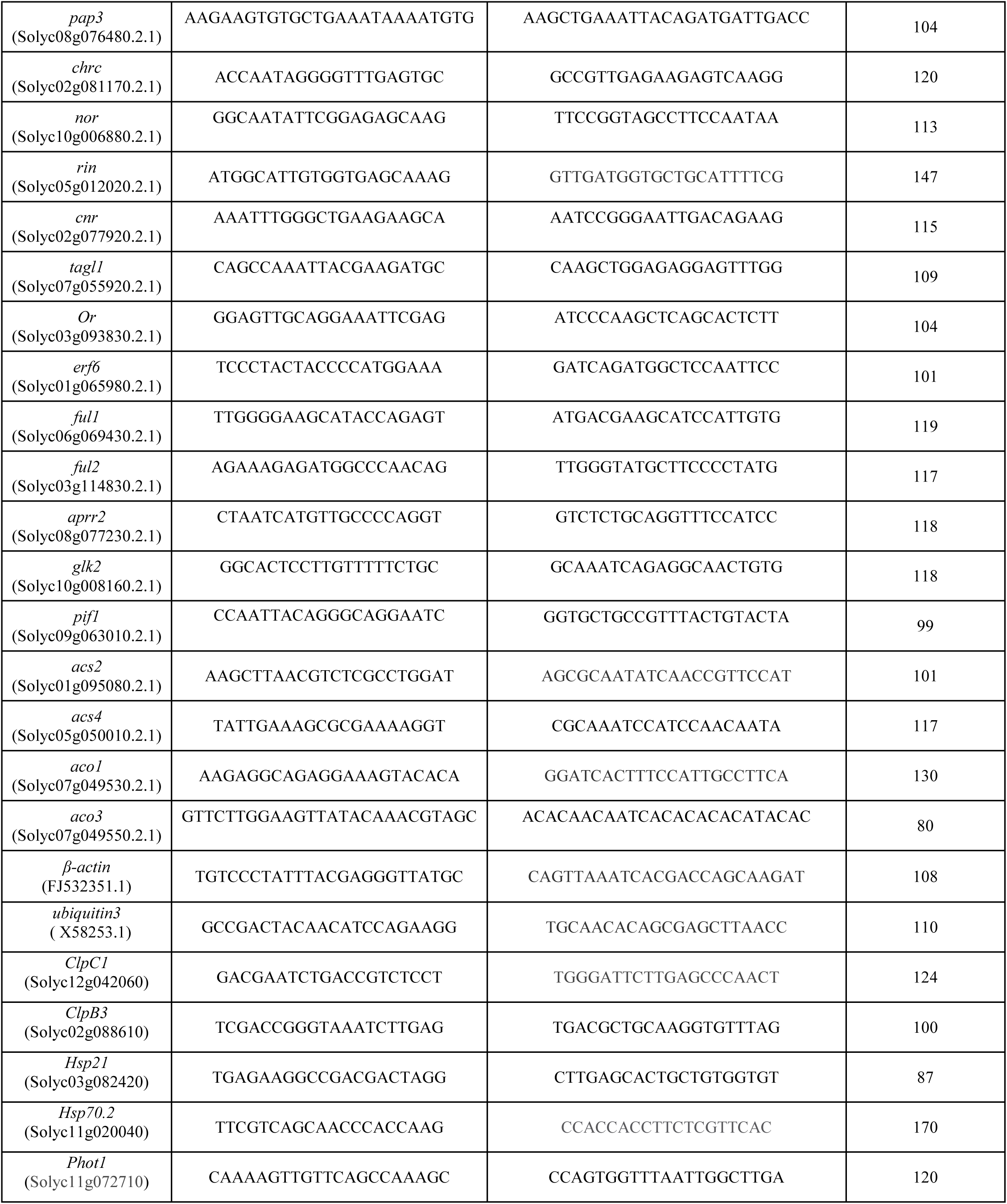
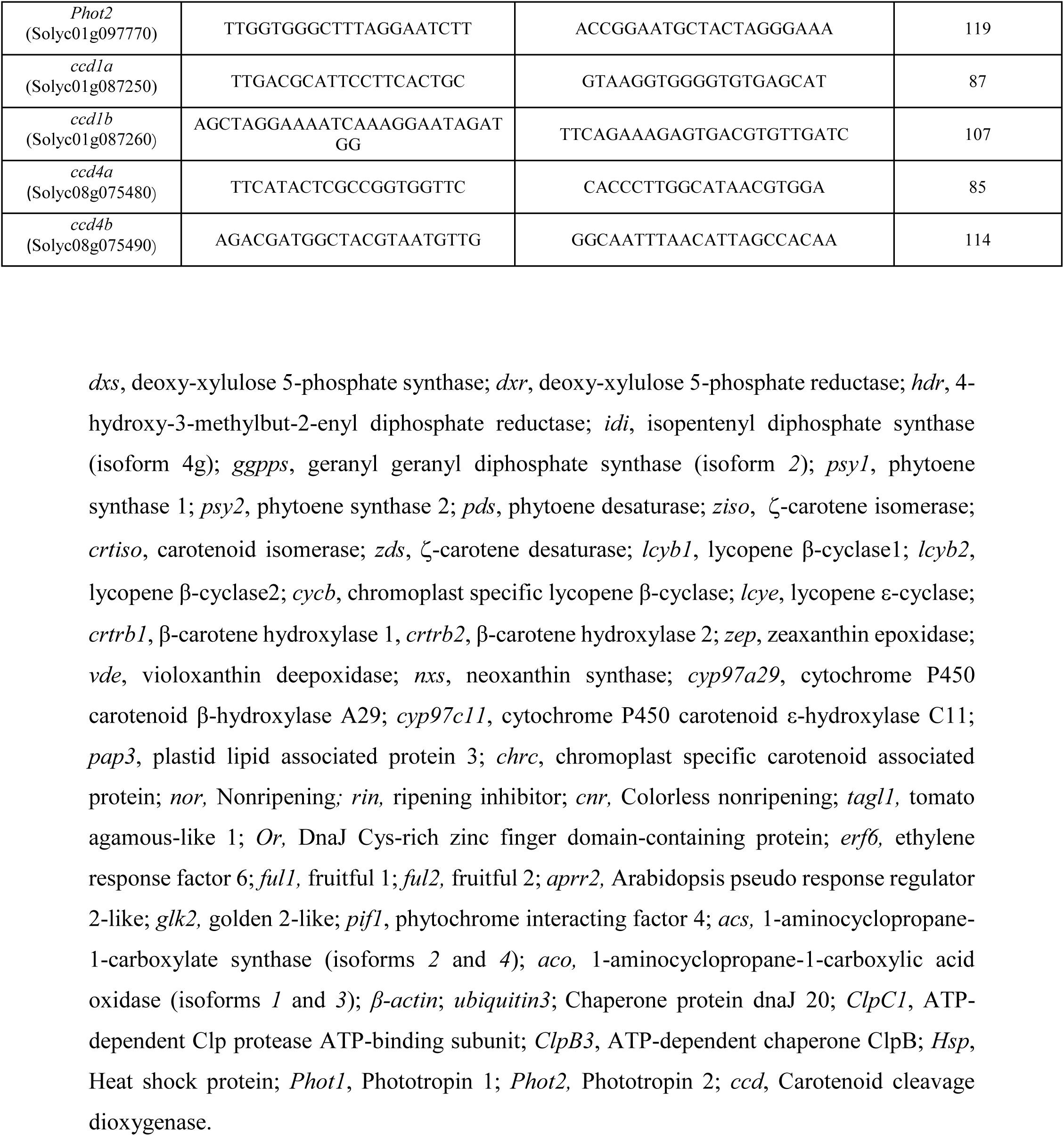
Primers used for quantitative real-time PCR analysis in this study.

